# Acyl amides derived from a bacterial symbiont control development of its nematode host

**DOI:** 10.64898/2026.06.15.732422

**Authors:** Li Su, Daniela V. Barahona, Michelle A. Diano, Krishna P. Patel, Sebastian Kaiser, Kevin Hoffman, Edna Bode, Rogelio Hernández Tamayo, Ali Khoshouei, Richard F. Collins, Martin Thanbichler, Jesko Koehnke, Helge B. Bode

**Affiliations:** Department of Natural Products in Organismic Interactions, Max Planck Institute for Terrestrial Microbiology, 35043 Marburg, Germany; Center for Research in Cellular and Molecular Biology (CIBCM), University of Costa Rica, San José 11501-2060, Costa Rica; Escuela de Zootecnia, University of Costa Rica, San José 11501-2060, Costa Rica; Institute of Food Chemistry, Leibniz University Hannover, Hannover, Germany; Center for Synthetic Microbiology (SYNMIKRO), Philipps University of Marburg, 35043 Marburg, Germany; Institute of Phototrophic Microbiology, Heinrich-Heine-Universität Düsseldorf, 40225 Düsseldorf, Germany; Faculty of Biology, Medicine and Health, University of Manchester, Oxford Road, Manchester, M13 9PT, UK; Department of Biology, Phillips University Marburg, 35043 Marburg, Germany; Department of Chemistry, Philipps University of Marburg, 35043 Marburg, Germany

## Abstract

Symbiotic bacteria frequently regulate host development through small-molecule signals. In the entomopathogenic *Steinernema* nematode *– Xenorhabdus* bacterium symbiosis, which is widely used in biological pest control, the signals that trigger nematode development have remained unknown. Using a phenotypic screen of *Xenorhabdus* mutants, we identified bacterial acyl amides, which we named stripteamides, as signals required for efficient recovery of *Steinernema* infective juveniles. Stripteamides are synthesized from phenylethylamine and activated fatty acid substrates by StaS, a previously uncharacterized amide synthase. Mechanistic and structural analyses indicate that StaS evolved from the initiation module of a nonribosomal peptide synthetase (NRPS) and was repurposed to generate signaling metabolites for host development. Stripteamide signaling is conserved across *Xenorhabdus* species and operates in different *Steinernema* hosts. These findings reveal a new function for widespread bacterial acyl amides that previously have been shown as cytotoxic and affecting bacterial quorum sensing, highlighting the context dependence of microbial natural product language.

## Introduction

Symbiotic microorganisms regulate many aspects of host development and physiology through small-molecule signals^1-7^. In animals, bacterial metabolites can shape immune maturation, epithelial homeostasis, behavior, and developmental transitions, illustrating that microbial chemistry can function as a molecular interface between symbiotic partners^2,8^. However, even in well-studied mutualisms, the identities, biosynthetic origins, and biological roles of the metabolites that trigger specific host transitions often remain poorly understood.

Entomopathogenic nematodes (EPNs) of the genera *Steinernema* and *Heterorhabditis* form obligate mutualistic associations with bacteria of the genera *Xenorhabdus*^9,10^ and *Photorhabdus*^11,12^, respectively^13^. Together, these partners act as highly effective insect pathogens and have been widely exploited as biological control agents in agriculture^14,15^. The free-living infective juveniles (IJs) carry their bacterial symbionts in a developmentally arrested state and invade insect hosts in search of suitable environments for reproduction. Once inside the insect hemocoel, the nematodes release their symbiotic bacteria^16^, which rapidly proliferate and remodel the host environment through the production of diverse specialized metabolites and enzymes^17^ (Supplementary Figure 1).

**Figure 1.**
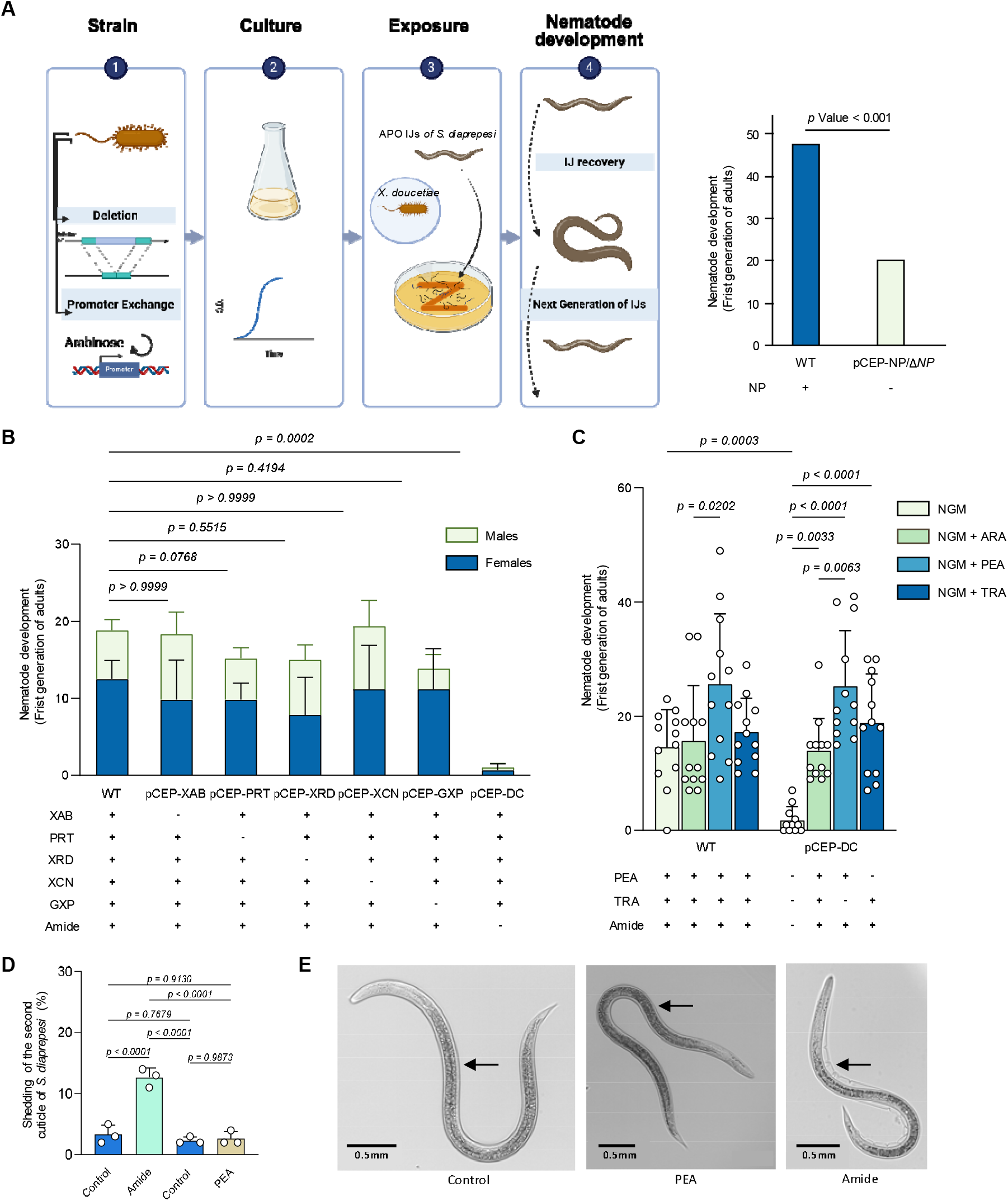
A phenotypic screen of *Xenorhabdus* strains identifies stripteamides as essential signals for nematode development. **(A)**, Methodology used to analyze the effect of different NPs on the development of nematode. Aposymbiotic *Steinernema diaprepesi* infective juveniles (APO IJs) were exposed to mutant strains of *X. doucetiae* harboring deletions or arabinose (ARA)-inducible promoter exchanges in biosynthetic gene clusters (BGCs) for specific natural products. After 4 days of incubation at 24°C, the number of individuals that developed into adults was determined by counting females and males to analyzed in the absence of which NPs this value decreased (Created with BioRender.com). **(B)**, Evaluation of nematode development in the presence and/or absence of XAB (xenoamicin), PRT (protegomycin), XRD (xenorhabdin), XCN (xenocoumacin), GXP (GameXPeptide), and Amide (phenylethylamides and tryptamides) by exposing them to mutant strains deficient in their synthesis. **(C)**, Analysis of nematode development in the presence of *X. doucetiae* WT or pCEP-DC in which the amide biosynthetic pathway was restored by supplementing the medium with PEA (phenylethylamine) and TRA (tryptamine) or by ARA induction. **(D)**, Process of removing the second cuticle (second shell) retained by the infective stage of the nematode as a method of protection when developing as a free-living stage. (**E**), Representative phase-contrast microscopy images of infective juveniles of *S. diaprepesi* under control, PEA, and amide treatments. Arrows indicate the second cuticle in all pictures. Scale bars are shown in each panel. Data in **B** (*n* = 6) and **c** (*n* = 12) represent biologically independent experiments. Bars in **B** and **C** indicate mean ± s.d., *P* values calculated with the Tukey’s multiple comparisons test.

Following bacterial proliferation, the nematode must transition from the growth-arrested infective juvenile stage carrying a second cuticle for environmental resistance to active development and reproduction^18^. In the *Heterorhabditis–Photorhabdus* symbiosis, the bacterial metabolite isopropylstilbene (IPS)^19^ functions as a key chemical cue that triggers this “nematode recovery”^20^. In contrast, the molecular signals that initiate development in *Steinernema* nematodes have remained unknown, leaving a major gap in our understanding of chemical communication within this commercially relevant bacterium–nematode–insect interaction.

Here we identify bacterial acyl amides, termed stripteamides, as symbiotic signals that promote the developmental recovery of *Steinernema* infective juveniles and trigger shedding of the second cuticle. Using genetic, chemical, and biochemical approaches in the *Steinernema diaprepesi – Xenorhabdus doucetiae* symbiosis, we show that stripteamides are synthesized from phenylethylamine (PEA) and fatty acids by a previously uncharacterized amide synthase, StaS. Functional and comparative analyses reveal that StaS evolved from the initiation module of the nonribosomal peptide synthetase (NRPS) XfpS, illustrating how NRPS-derived enzymatic scaffolds can be repurposed for small-molecule signaling.

## Results

### Symbiotic amides are identified as critical signals for nematode development

To identify bacterial metabolites that control nematode development, we examined the recovery of aposymbiotic (APO) *Steinernema diaprepesi* infective juveniles (IJs) in the presence of mutants of the symbiotic bacterium *Xenorhabdus doucetiae* DSM 17909 (Figure 1A, Supplementary Figure 2). APO IJs lack their cognate bacterial symbiont and therefore provide a tractable system to test which bacterial metabolites restore developmental recovery.

We first tested whether major classes of bacterial natural products contribute to this developmental transition. Using promoter-exchange mutants generated via the easyPACId (easy Promoter Activation for Compound Identification) system^21^, which puts specific biosynthetic pathways under the control of L-arabinose (ARA), we evaluated the impact of several prominent *Xenorhabdus* metabolites (Supplementary Figure 3). Exposure of APO IJs to wild-type *X. doucetiae* grown on NGM (nematode growth medium) agar plates resulted in recovery and development of an average of 18.4 adults per 50 IJs (Figure 1B). Comparable recovery rates were observed for easyPACId mutants without ARA, deficient in the production of xenoamicin (XAB)^22^, protegomycin (PRT)^23^, xenorhabdin (XRD)^24,25^, xenocoumacin (XCN)^26^, and GameXPeptide (GXP)^27,28^. Thus, loss of these major natural product classes did not impair nematode recovery under the assay conditions.

In contrast, APO IJs exposed to the pCEP-DC mutant without ARA exhibited a dramatic reduction in adult recovery compared to the wild-type control (Figure 1B). Because DC represents the main decarboxylase that provides the amines required for phenylethylamide and tryptamide formation^29^, this phenotype suggested that amine-derived metabolites play a critical role in nematode development.

To test this hypothesis, we performed supplementation assays using either the inducer ARA, which restores expression of the decarboxylase gene, or the amine precursors themselves. In the wild-type background, nematode recovery was largely unaffected by supplementation with ARA, PEA, or TRA (tryptamine). In contrast, the developmental defect of the pCEP-DC mutant was fully rescued by either ARA induction or supplementation with PEA or TRA (Figure 1C). These results demonstrate that disruption of amine-derived metabolite production abolishes nematode recovery, whereas restoration of this pathway restores normal development.

Since complementation of the pCEP-DC mutant with PEA does not distinguish whether PEA itself or its downstream amide products drive the observed effect, we directly compared PEA with a medium-chain length amide (C12-acyl, **11**) in IJs assays. **11** is an abundant amide in *X. doucetiae* cultures^29^. IJs were incubated with each compound in distilled water for 1h, transferred to NGM agar plates and monitored by microscopy. While amide **11** induced shedding of the second cuticle after 60 min, neither PEA nor the control had this effect (Figure 1D and E, Supplementary Movies 1-3). As the phenylethylamides are involved in stripping off the cuticle, we named them stripteamides.

### Biosynthesis of stripteamides by the amide synthase StaS

Having established that amine-derived metabolites are required for nematode recovery, we next sought to identify the bacterial enzyme responsible for their biosynthesis. Chemical analysis of *X. doucetiae* cultures revealed a series of PEA- and TRA-derived fatty acid amides with varying acyl chain lengths (Figure 2)^29^. The TRA derived stripTRAmides were produced at lower abundance and with fewer detectable variants compared than the PEA-derived stripteamides (Supplementary Figure 3 and 4). All compounds were identified by LC–MS/MS analysis and comparison with synthetic standards^30^.

**Figure 2.**
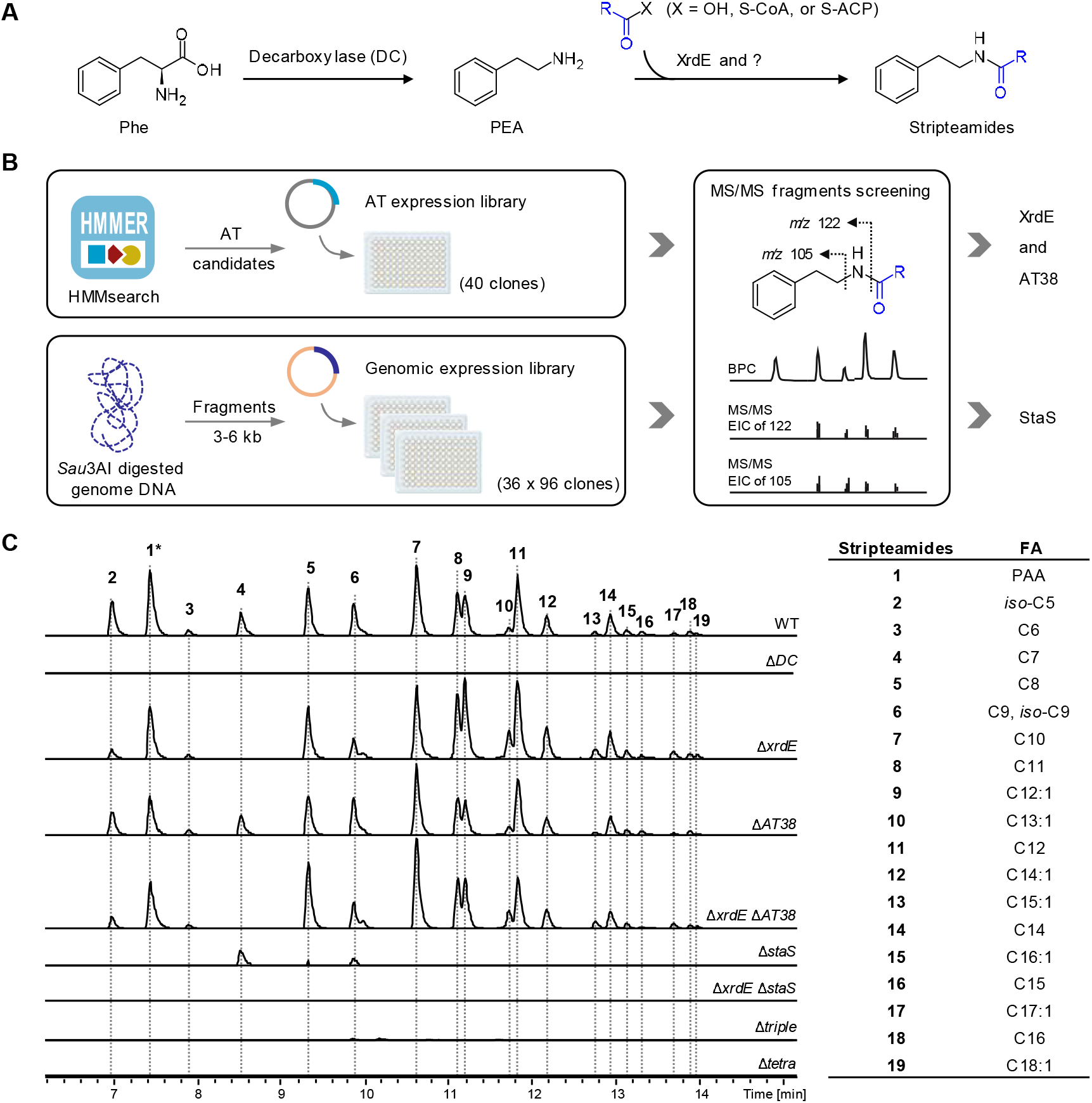
Identification of the stripteamide biosynthetic pathway and the key amide synthase StaS. **(A)** Proposed biosynthetic pathway for stripteamide initiates with decarboxylation of phenylalanine (Phe) to phenylethylamine (PEA) catalyzed by the decarboxylase DC. Subsequent acylation of PEA with various fatty acid chains (either as free acids or activated thioesters) is mediated by XrdE and at least one additional, unidentified enzyme. **(B)**, Strategy for identifying the unknown enzyme. Two parallel screening approaches were employed: (i) a targeted acyltransferase (AT) expression library constructed based on hidden Markov model (HMM) searches of the *X. doucetiae* genome, and (ii) an unbiased genomic expression library. Clones cultured in medium with PEA supplementation were screened for stripteamide production using the diagnostic fragment ions *m/z* 105 and 122 ([M+H]^+^). **(C)**, Extracted ion chromatograms (EICs) showing the production profiles of stripteamides **1**–**19** in the *X. doucetiae* WT strain and its derivative mutants. Mutants analyzed include Δ*DC*, Δ*xrdE*, Δ*AT38*, Δ*xrdE* Δ*AT38*, Δ*staS*, Δ*xrdE* Δ*staS*, and the triple mutant Δ*xrdE* Δ*AT38* Δ*staS* (denoted Δ*triple*), and the quadruple mutant Δ*DC* Δ*xrdE* Δ*AT38* Δ*staS* (denoted Δ*tetra*). The asterisk (*) indicates that the peak intensity for stripteamide **1** is shown at 10-fold reduction for better visualization. Fatty acyl chain structures were validated in our previous studies^24,25^.

Our previous work identified XrdE^29,31^, the *N*-acyltransferase (NAT) involved in the biosynthesis of the antibiotic XRD, as capable of transferring specific medium-chain fatty acids (C6, *iso-*C7, C8 and C9) to PEA^29^, generating stripteamides **3, 4, 5** and **6** (Figure 2C and Supplementary Figure 5). However, deletion of *xrdE* in the WT strain abolished production of stripteamide **4** and reduced stripteamide **6**, but did not eliminate the broader stripteamide profile (Figure 2C). This suggested that additional acyltransferases (ATs) contribute to stripteamide biosynthesis.

To identify such ATs, we performed a hidden Markov model (HMM) search against the *X. doucetiae* genome, yielding an initial set of 40 non-redundant AT candidate genes (including XrdE) (Figure 2B and Supplementary Table 2). Heterologous expression of these 40 AT candidates individually in *E. coli* revealed that only AT38, besides XrdE, could catalyze the formation of stripteamides **3, 5, 7, 9**, and **11** upon the feeding of substrate PEA in medium (Supplementary Figure 5). However, deletion of *AT38* did not impair overall amide production in either the WT or the Δ*xrdE* mutant strain (Figure 2C). Collectively, these results indicate that the key enzyme responsible for stripteamide biosynthesis likely does not belong to the canonical NAT family. We therefore performed a gain of function screen by expressing a genomic library of 3-6 kb fragments from *X. doucetiae* in *E. coli* analyzing 36 × 96 clones for stripteamide production in PEA supplemented medium (Figure 2B).

We identified several clones capable of producing stripteamides besides the already identified ATs XrdE and AT38 (Supplementary Figure 6A). Sequencing of the inserts from positive clones revealed that two independent clones, P2R6C7 and P5R7C7, carried *XDD1_1031*, which encodes an NRPS-derived condensation (C) domain specifically connecting two L-amino acids (^L^C_L_) (Supplementary Figure 6A). To assess its biochemical function, we cloned and expressed *XDD1_1031* (construct pSUL58) in *E. coli* DH10B. When PEA was supplied in the medium, the pSUL58 expression strain produced a spectrum of stripteamides closely resembling that of *X. doucetiae* WT (Supplementary Figure 6B). Genetic validation further confirmed the role of *XDD1_1031* in stripteamide biosynthesis, since the production profile of stripteamide was consistently and significantly altered by deleting the *XDD1_1031* gene across multiple genetic backgrounds, including WT, Δ*xrdE*, and Δ*xrdE* Δ*AT38*. Notably, inactivation of *XDD1_1031* in the WT background restricted the production of stripteamides to only **4, 5**, and **6**, a phenotype mirroring that of the Δ*xrdE* mutant (Figure 2C). These findings establish that the protein encoded by *XDD1_1031* is the principal enzyme responsible for stripteamide biosynthesis in *X. doucetiae*, catalyzing the amidation of medium- to long-chain (C5 to C18) fatty acids with PEA. We therefore named this enzyme <u>st</u>ripte<u>a</u>mide <u>s</u>ynthase (StaS).

### PEA and stripteamides jointly promote nematode development

Based on the clarified the biosynthetic pathway, we could now determine the contribution of the amine precursor (PEA) or the amide products (stripteamides) for optimal nematode development. To this end, we examined the nematode development in four strains: WT, Δ*DC* (PEA-deficient), Δ*triple* (Δ*xrdE* Δ*AT38* Δ*staS*, stripteamide-deficient) and Δ*tetra* (Δ*DC* Δ*xrdE* Δ*AT38* Δ*staS*, deficient in PEA and stripteamide) (Figure 3A). The strains were cultured on NGM medium with or without 1 mM PEA supplementation. In the WT control group, the number of adults developed from IJs was comparable under both conditions. For the Δ*DC* mutant, PEA supplementation fully restored nematode development to WT levels. In contrast, the Δ*triple* mutant supported only partial development regardless of PEA supplementation. Notably, the Δ*tetra* mutant was virtually incapable of supporting nematode development in the absence of PEA. However, upon PEA supplementation, it behaved similarly to the Δ*triple* mutant, achieving only partial developmental recovery. These results demonstrate that PEA can partially support development, but full recovery requires an intact stripteamides biosynthetic pathway. Thus, efficient nematode development depends on the combined action of the amine precursor and StaS-dependent amide products.

Given that *X. doucetiae* produces a suite of stripteamides with varying acyl chain lengths (Figure 2C), we aimed to identify which chain lengths are functionally relevant for nematode development. For this, WT and Δ*tetra* were cultured on NGM and NGM supplemented with 1 mM of individual synthetic stripteamide variants bearing acyl chains of C3 (stripteamide **20**), C6 (**3**), C8 (**5**), C12 (**11**), C14 (**14**), C16 (**18**), and C22 (**21**) (Figure 3B). For the WT strain, no significant differences in nematode development were observed across most supplemented conditions. However, supplementation with the long-chain stripteamide **21** (C22) markedly inhibited development compared to all other treatments, consistent with previous finding suggesting that long-chain stripteamides may exert cytotoxic effects^29,30^. For the Δ*tetra* mutant, supplementation with short-chain (C3) or long-chain (C16 and C22) stripteamides failed to rescue development. Nevertheless, medium-chain variants (C6, C8, C12, C14) significantly restored the development relative to the unsupplemented control. These findings indicate that the medium-chain stripteamides (C6–C14) are the active species responsible for promoting nematode recovery and development to the first-generation adult stage.

**Figure 3.**
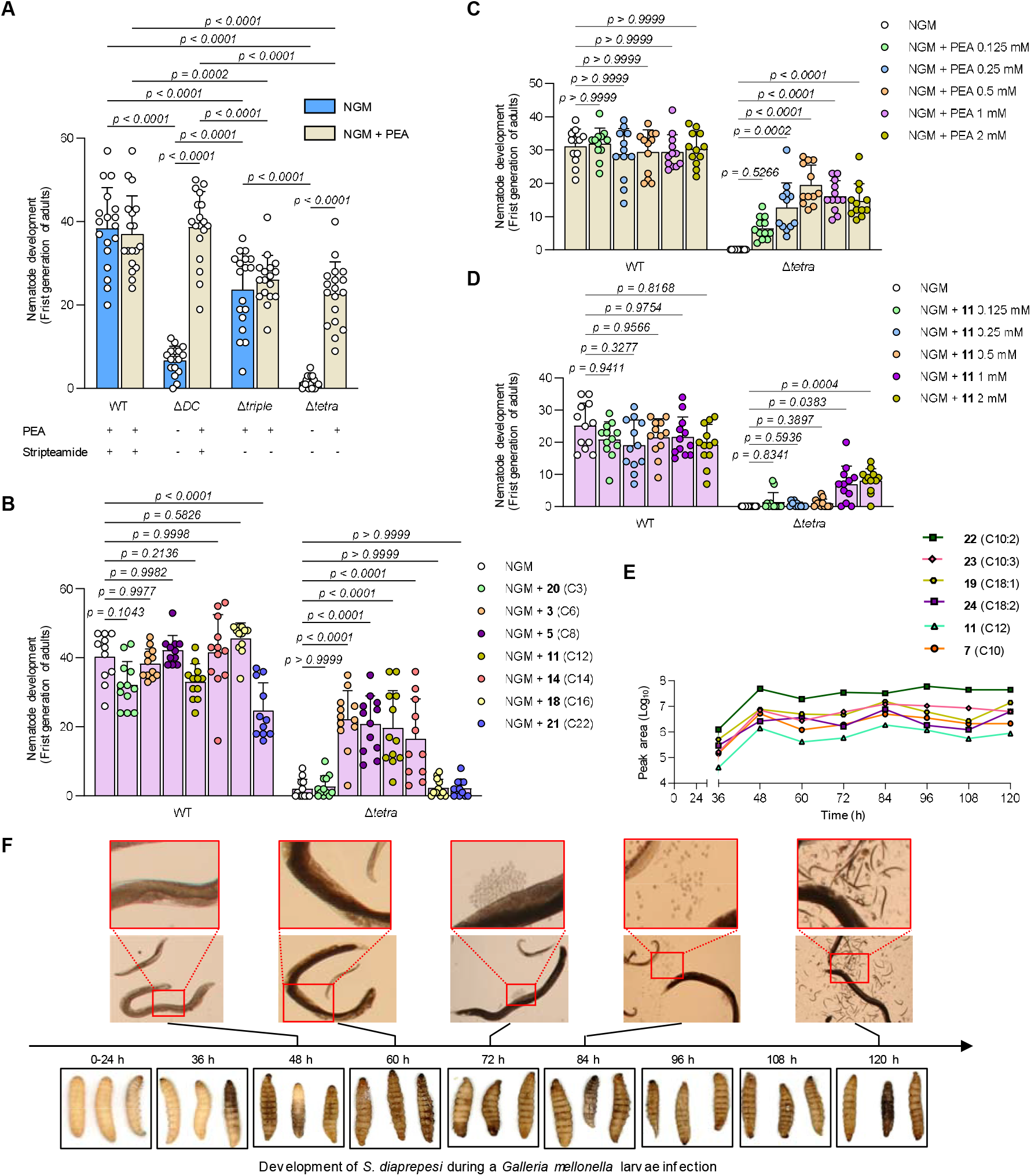
PEA and stripteamides are required for *Steinernema diaprepesi* development, with chain length-dependent activity and temporal correlation with insect infection. **(A)**, Evaluation of the nematode development of *S. diaprepesi* APO IJs after 4 days of incubation at 24°C exposed to *X. doucetiae* WT, Δ*DC*, Δ*triple*, and Δ*tetra* in NGM and NGM supplemented with PEA at a final concentration of 1mM. (**B**), Evaluation of the nematode development of *S. diaprepesi* APO IJs after 4 days of incubation at 24°C exposed to *X. doucetiae* WT and Δ*tetra* in NGM and NGM supplemented with 1 mM of individual synthetic stripteamides **20** (C3), **3** (C6), **5** (C8), **11** (C12), **14** (C14), **18** (C16) and **21** (C22) bearing straight-chain fatty acids of indicated lengths. (**C, D**), Determination of the optimum concentration of the compounds for the development of nematodes from IJs to first generation adults in supplemented media with ascending concentrations of PEA and stripteamide **11** respectively. **(E)**, Time-course quantification of stripteamide production during infection. Hemolymph was collected from *Galleria mellonella* instar larvae at indicated time points following infection with *S. diaprepesi* carrying its native symbiont *X. doucetiae*. Peak area of these most abundant derivatives was measured by HPLC-MS. Data represent mean peak area from three independent infections (*n* = 3 per time point). **(F)**, Development of *S. diaprepesi* during *Galleria mellonella* larvae infection at different time points. The nematodes stages and developmental steps are shown: females and males (48–60 h), egg deposition (72 h), egg hatching and juveniles (84–96 h), and a mixture of different stages (120 h). Representative images show the symptoms of *G. mellonella* larvae at the indicated times after infection by the *S. diaprepesi*-*X. doucetiae* symbiotic complex. Data in **a** (*n* = 18), and **B**–**D** (*n* = 12) represent biologically independent experiments. Bars in **A**–**D** indicate mean ± s.d., *P* values calculated with the Tukey’s multiple comparisons test.

Dose–response assays with Δ*tetra* supplemented with varying concentrations (0.125, 0.25, 0.5, 1, and 2 mM) of either PEA or stripteamide **11** (C12) revealing that slightly lower concentrations of PEA (0.5 mM) are needed (Figure 3C) compared to 1-2 mM of **11** (C12) (Figure 3D).

To understand these findings within the process of natural infection, we proceeded to analyze the onset timing and accumulation of stripteamides at various stages of nematode infection of insects, alongside nematode development within the insect host. Larvae of the model insect *Galleria mellonella* were infected with 50 IJs of the nematode *S. diaprepesi*, which contained its cognate bacterial symbiont *X. doucetiae*. Subsequently, hemolymph samples were collected from the anterior section of the insect at 12-hour intervals post-infection to analyze the synthesis of stripteamides while parallel dissections were performed to monitor nematode development. Stripteamide derivatives became detectable during the 36–48 h window post-infection and remained present until 120 h (Figure 3E, Supplementary Figure 8A and B). Interestingly, the stripteamide profile diverged from that observed in lab-cultured *X. doucetiae*: In insect hemolymph, the C10:2 derivative (**22**, *m/z* 272.20) was the most abundant, followed by C18:1 (**19**, *m/z* 386.34), C10:3 (**23**, *m/z* 270.18), C18:2 (**24**, *m/z* 384.32). By contrast, C10 (**7**, *m/z* 276.23) and C12 (**11**, *m/z* 304.26), the most abundant species in bacteria-only cultures were present at much lower levels in hemolymph samples. These observations reveal a predominance of long-chain unsaturated fatty acid amides within the infected host hemolymph (Figure 3E, Supplementary Figure 8A & B) as described previously for whole-insect extracts^29^. The corresponding stripTRAmides also favored long unsaturated chains, with C18:1 and C18:2 derivatives being dominant (Supplementary Figure 8A,B). Another sampling every 3 h between 36 and 48 hours post-infection revealed initiation of stripteamide production around 36-39 h, followed by linear accumulation which plateaued at 48 h (Supplementary Figure 8C). Parallel assessment of insect symptoms showed that larvae remained mobile with no visible changes until 24–36 h, at which point one of three individuals displayed sign of melanization, evident as dark coloration spots^32^. By 48 h post-infection, all insects had died, displaying progressive melanization and loss of turgor (Figure 3F). Dissection of infected insects at matched time points revealed that nematodes had developed to the young adult (J4) stage at 48 h, with females and males being comparable in size (Figure 3F). Females reached gravidity by 60 h, with egg deposition and hatching observed from 72 to 108 h (Figure 3F). Taken together, these findings place stripteamide production within the temporal window of nematode developmental recovery and insect mortality, consistent with a role for StaS-dependent metabolites during the transition from IJ to reproductive adult within the host environment.

### Evolutionary origin of the stripteamide synthase StaS

StaS has an NRPS-like domain architecture but deviates from canonical NRPS organization. Specifically, StaS consists of an NRPS Condensation (C) domain fused to a truncated adenylation (A)-like region (Supplementary Figure 9), suggesting an evolutionary origin from an ancestral NRPS enzyme. Phylogenetic analysis showed that the StaS C domain branches close to other ^L^C_L_ domains but forms its own monophyletic group. Notably, the StaS C domain branches inside the first C domains of Xefoampeptide synthetases (XfpS)^33^ (Figure 4A). XfpS is a canonical multimodular NRPS responsible for the biosynthesis of xefoampeptides, cyclic lipopeptides composed of a tripeptide linked to a β-hydroxy fatty acid^34^ (Supplementary Figure 10). Examination of *staS* and *xfpS* distribution across the *Xenorhabdus* phylogeny (Supplementary Figure 11A) revealed that the two genes are adjacent in several species, including *X. vietnamensis* (Supplementary Figure 11B), consistent with duplication followed by functional divergence. Overall, *staS* seems to occur in specific subclades, which is additionally supported by stripteamide detection^33^.

**Figure 4.**
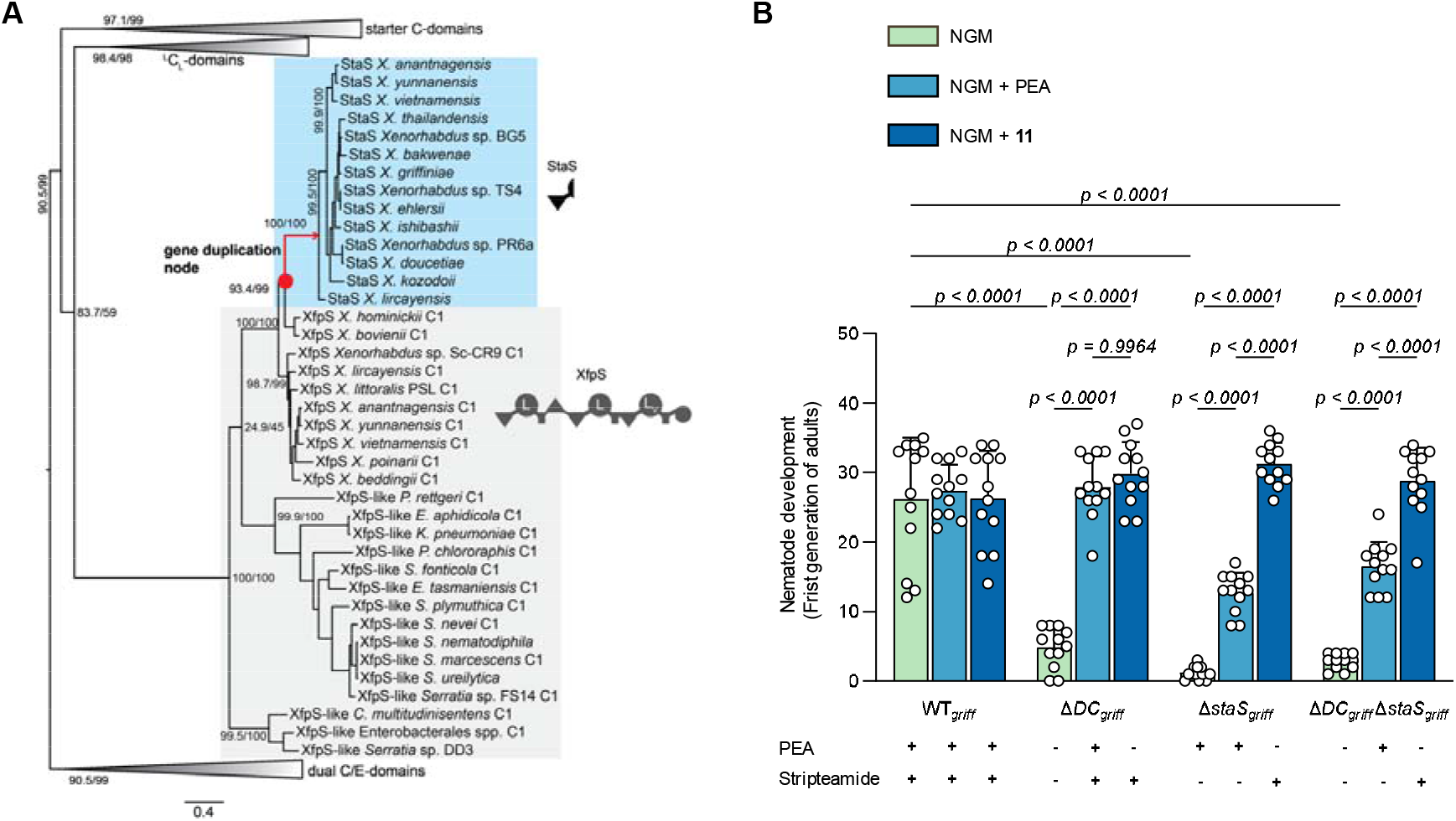
Evolution and function conservation of StaS. **(A)**, Phylogenetic analysis of StaS C-domain. The tree was inferred from 352 C-domains extracted from various NRPSs, including ^L^C_L_, ^D^C_L_, dual C/E and starter C-domains. StaS branches within the starter C⍰domains of the xefoampeptide synthetases (XfpS) and forms a distinct monophyletic group. NRPS domain representation is as follows: Condensation (triangle pointing downwards), epimerization (triangle pointing upwards), adenylation (large [partial] circle), thioesterase (small circle), thiolation (rectangle). **(B)**, Conserved stripteamide signaling observed in *Steinernema hermaphroditum–Xenorhabdus griffiniae* system. Development of aposymbiotic infective juveniles (IJs) to adults of *Steinernema hermaphroditum* was assessed after 4□days of incubation at 24□°C on nematode growth medium (NGM) seeded with *Xenorhabdus griffiniae* WT, Δ*DC*_*griff*_, Δ*staS*_*griff*_ *and* Δ*DC*_*griff*_ Δ*staS*_*griff*_ mutants. Media were supplemented with 1□mM PEA or 1□mM stripteamide **11** (C12). Data shown here represents biologically independent experiments (*n* = 12). Bars indicate mean ± s.d., *P* values calculated with the Tukey’s multiple comparisons test.

To determine whether XfpS-containing strains retain the ability to produce stripteamides and whether StaS homologs generate related metabolites, we analyzed culture extracts from representative *Xenorhabdus* strains by LC–MS. These included strains carrying only StaS (*X. griffiniae*) (Supplementary Figure 12), only XfpS (*X. hominickii, X. bovienii, X. poinarii*, and *X. beddingii*) (Supplementary Figures 13–16), or both (*X. vietnamensis*) (Supplementary Figure 17). Stripteamides were detected only in strains harboring *staS*, indicating that StaS represents a specialized derivative of a more broadly distributed NRPS ancestor likely linked to its role in nematode development.

We next asked whether stripteamide signaling is unique to *X. doucetiae* or reflects a conserved feature of *Xenorhabdus* symbioses. To address this, we turned to the *Steinernema hermaphroditum*– *Xenorhabdus griffiniae* system as a second model^35^. Consistent with our findings in the *S. diaprepesi– X. doucetiae* system, development of *S. hermaphroditum* was severely impaired in mutants unable to produce either PEA or stripteamides (Supplementary Figure 12), and was fully restored only when both compounds were available (either via endogenous synthesis or exogenous supplementation) (Figure 4B). However, in contrast to *S. diaprepesi*, where PEA exhibited higher potency, stripteamide showed markedly stronger bioactivity than PEA in *S. hermaphroditum*. This was evidenced by the pronounced difference between PEA and stripteamide **11** treatments observed in both Δ*staS*_*griff*_ and Δ*DC*_*griff*_ Δ*staS*_*griff*_ mutants (Figure 4B), suggesting that the final amide product serves as the primary signaling molecule in this host. Collectively, these findings demonstrate that both stripteamides biosynthesis and its signaling function are conserved across selected *Xenorhabdus* species, pointing to a general mechanism by which these bacteria regulate developmental transitions in their *Steinernema* hosts.

### StaS is an NRPS-derived amide synthase that uses activated acyl donors

As mentioned above, StaS contains a truncated A domain region, which raises the question of whether this region retains catalytic activity or fulfils a structural role instead. To examine the functional importance of this A-like region, we separated the two domains of StaS at the conserved WNATE motif^36^ within the C–A interdomain linker and reassembled them using the SYNZIP (SZ) 17/18 leucine zipper pair^37^. Expression of either domain alone failed to support stripteamide production (Figure 5A, traces ii and iii), whereas co-expression of both fragments restored amide synthesis to levels comparable to full-length StaS (Figure 5A, traces i and iv).

**Figure 5.**
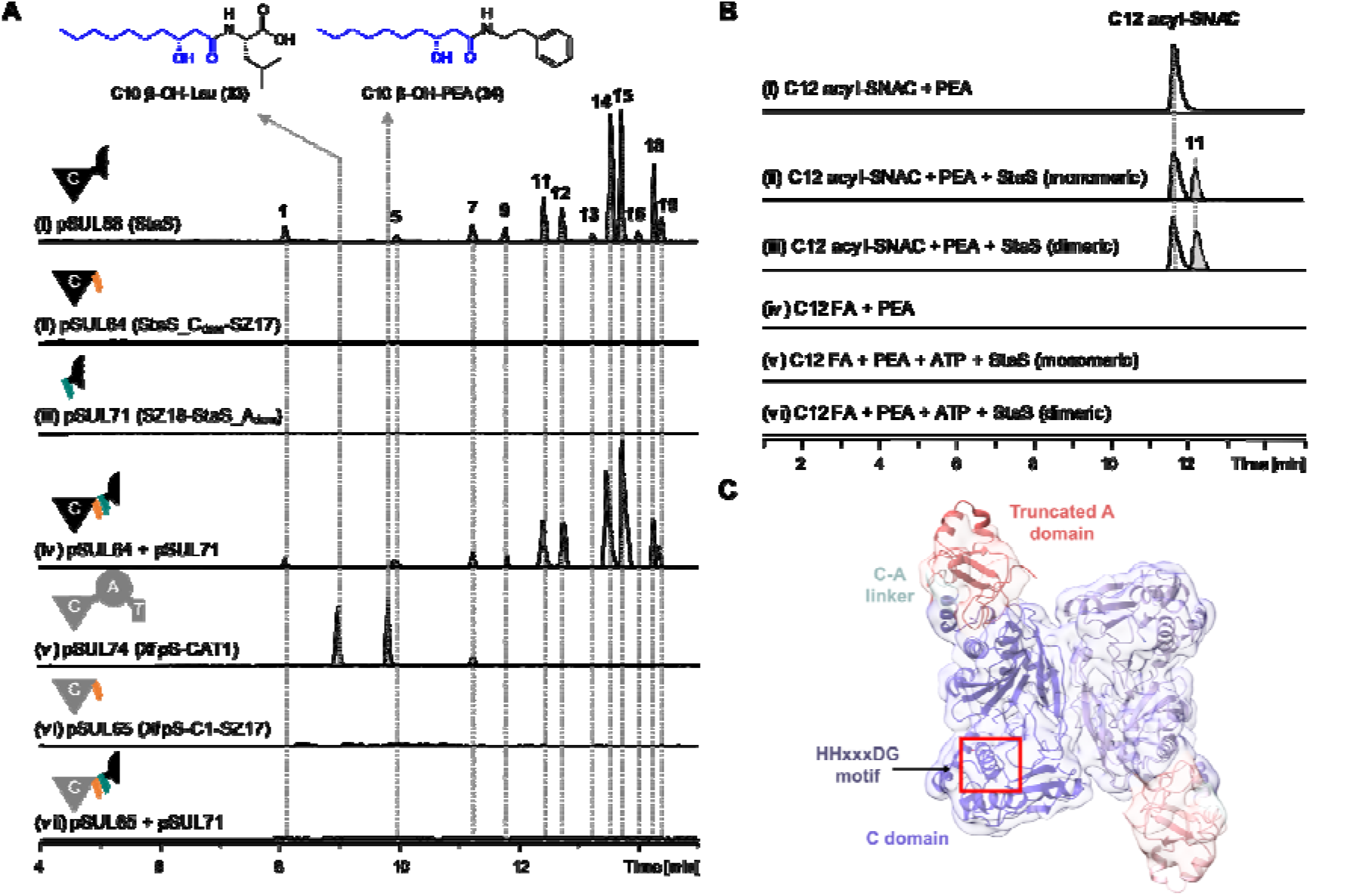
Structure and mechanistic characterization of StaS. **(A)**, Domain architecture of StaS and its domain-splitting, splicing and swapping experiments. Extracted ion chromatograms (EICs) show stripteamide production in *E. coli* DH10B strains harboring the indicated plasmids and PEA in the growth medium. For plasmid pSUL74, the host strain is *E. coli* DH10B::mtaA. For NRPS domain explanation see Fig. 4. **(B)**, *In vitro* enzymatic assays of monomeric and dimeric StaS using either an activated acyl thioester (C12 acyl-SNAC) or free fatty acid (C12 FA) as substrate. **(C)**, Cryo-EM structure of the StaS homodimer (PDB 31EM) at 4.01 Å with the conserved HHxxxDG motif of the condensation domain highlighted in the red box. The Coulomb potential map is shown as a transparent surface and colored according to the respective protein parts.

Prompted by this finding and the inability of XfpS to produce stripteamides (Supplementary Figures 13–16), we expressed the initiation module of XfpS (XfpS-CAT1) and the isolated C1 domain in *E. coli*. Expression of the full CAT1 module generated the expected product derived from its native β-hydroxy fatty acid and amino acid substrate (C10 β-OH-Leu, compound **33**) (data not shown). When PEA was supplied, the corresponding amide (C10 β-OH-PEA, compound **34**) was also detected (Figure 5A, trace v), indicating that amide-forming activity is present by XfpS but is only detectable when a defined portion is expressed. In contrast, the isolated C1 domain produced no detectable products (Figure 5A, trace vi). We then tested whether the StaS A-like region could functionally replace the missing domains of XfpS-C1. A chimeric construct linking the StaS A-like region to XfpS-C1 failed to restore activity (Figure 5A, trace vii), suggesting that StaS and XfpS rely on distinct structural contexts for catalysis.

We next investigated the chemical mechanism of StaS-catalyzed amide formation. In vitro assays revealed that StaS efficiently produced stripteamides when supplied with fatty acyl thioester (synthetic SNAC, N-acetylcysteamine, Supplementary Figures 18 and 19) and PEA (Figure 5B, traces i-iii), while no products were detected with free fatty acids and ATP (Figure 5B, traces iv-vi). Thus, StaS accepts activated acyl thioesters as substrates as other starter C domains but has no active A domain.

### Cryo-EM structure of StaS amide synthase homodimer

To place these biochemical findings into a structural context, we expressed and purified StaS and XfpS-CAT1 *in vitro* (Supplementary Figure 20) and determined the cryo-EM structure of StaS at 4.01 Å resolution with C2 symmetry (Figure 5C, Supplementary Figure 21). StaS eluted as two distinct species during size-exclusion chromatography, and we selected the dimer species for structural elucidation as the monomeric species was too small for reliable reconstruction using cryo-EM. SEC reinjection experiments showed that monomeric and dimeric StaS did not measurably interconvert (Supplementary Figure 22), and both forms displayed comparable *in vitro* activity for stripteamide generation (Figure 5B), indicating that dimerization is not required for catalysis under the assay conditions. The StaS structure revealed that each protomer is dominated by an NRPS condensation domain fold containing the conserved HHIIGDG motif, corresponding to the HHxxxDG motif characteristic of amide bond-forming C domains^38^. Rather than retaining a complete adenylation domain, StaS carries only a short N-terminal appendage homologous to part of the XfpS A1 domain, consistent with the loss of the canonical ATP-dependent substrate-activation machinery of NRPS A domains^39,40^. A local resolution map shows a compact dimer interface and low-resolution areas mainly around the truncated A domain as well as the N-terminal lobe of the C domain, which suggests that these regions are highly flexible (Supplementary Figure 23). The increased flexibility around the N-terminal lobe of the C domain is likely important for accommodating the activated acyl thioester substrates as this is also where the HHxxxDG motif is present. A comparison of the StaS structure with an AlphaFold3 model shows significant structural variation mainly at the N-terminal lobe of the C domain, with a 180° shift of secondary structure elements contained within residues 59-72 and 156-169 away from the dimer interface (Supplementary Figure 24), which also causes the rest of the N-terminal lobe to shift and deviate from a canonical condensation domain fold. It is unclear whether dimerization alone is causing this shift or whether this represents a novel type of fold within C domains that could explain its non-canonical substrate requirements. Higher resolution data will be required to further analyze these structural details.

Attempts to obtain a high-resolution structure of XfpS-CAT1 were not successful due to the presence of several different conformations of the protein, demonstrating inherent flexibility (Supplementary Figure 25). Therefore, we selected two distinct conformations and calculated initial low-resolution models with conformation 1 likely representing a more closed conformation relative to conformation 2. The presence of the full A domain and also the thiolation domain contributes to this flexibility, and future structural studies will require substrate analogues designed to lock XfpS-CAT1 into a more rigid conformation. We therefore compared the StaS structure to an AlphaFold3 model of XfpS-CAT1, which serves as a representative model for canonical NRPS enzymes, to determine whether these evolutionary related enzymes have conserved structural elements. The overall architecture of the two proteins is similar (Supplementary Figure 26A), but just as we observed with the StaS AF3 model, the N-terminal lobe portion of the C domain adopts a completely new conformation relative to the XfpS AF3 model (Supplementary Figure 26B). A closer look at the A domains shows a similar C-A domain interface and a highly conserved fold overall, but limited conclusions can be drawn due to the high flexibility in this region that has low resolution and some density lacking (Supplementary Figure 26C).

Overall, the StaS structure provides a molecular rationale for its biochemical behaviour: amide formation is centered on a retained C domain scaffold with altered conformation in a homodimeric state, whereas the A domain-derived segment appears to support the enzyme architecture rather than catalyze ATP-dependent fatty acid activation. Together with the split-and-reassembly experiment, these data support a model in which StaS evolved from an NRPS initiation module by reductive remodelling into a compact fatty acid amide synthase: the ancestral C domain retained amide-forming capacity, whereas the A-domain remnant was retained to stabilize the active conformation rather than to activate carboxylate substrates.

## Conclusion

Chemical signals produced by microbial symbionts frequently regulate key developmental transitions in their eukaryotic hosts, yet the identities and origins of these molecules remain poorly understood in many associations. In the entomopathogenic nematode (EPN) symbiosis relevant to agricultural insect pest control, bacterial metabolites have emerged as critical regulators of nematode host development, exemplified by the role of isopropylstilbene (IPS)^19^ in triggering recovery of *Heterorhabditis* infective juveniles. Our work reveals that some *Steinernema–Xenorhabdus* symbiosis rely on a fundamentally different signaling strategy based on a family of fatty acid amides, which we term stripteamides. This discovery expands the known symbiotic signals that regulate nematode development and provides new insight into host–microbe communication.

Beyond identifying the signaling molecules themselves, our findings provide insight into the evolutionary origin of this communication system. Mechanistic and structural analyses indicate that StaS represents a streamlined amide synthase derived from an ancestral nonribosomal peptide synthetase (NRPS) module. In contrast to canonical NRPS enzymes that activate substrates through ATP-dependent adenylation, StaS utilizes different preactivated acyl thioesters and aromatic amines, enabling the production of a library of chemically diverse family of fatty acid amides. These observations illustrate how conserved NRPS catalytic architectures can be repurposed via duplication and divergence through evolution to generate signaling molecules with distinct biological functions. The restriction of StaS homologs to *Xenorhabdus* species suggests that this enzymatic innovation emerged during the evolution of the *Xenorhabdus–Steinernema* partnership, providing the bacterium with a dedicated chemical mechanism to regulate its nematode host.

The identification of stripteamides also places fatty acid amides among a growing class of signaling molecules that regulate nematode physiology and behavior. In *Caenorhabditis elegans*, the endocannabinoid anandamide^41^ promotes preference for higher-quality food through NPR-19, a G-protein coupled receptor with functional similarity to mammalian CB1, demonstrating that fatty acid amides can influence complex behavioral decisions in nematodes. Although the molecular targets of stripteamides remain unknown, the developmental effects described here suggest that fatty acid amides may have been repeatedly recruited during evolution to regulate critical nematode life-history transitions. Whether stripteamides and endocannabinoids engage related signaling pathways remains an intriguing question for future investigation.

Our results further suggest that stripteamide signaling operates through a composite system involving both the amine precursor and the final amide products. While PEA strongly stimulates nematode development in *S. diaprepesi*, different nematode hosts may respond preferentially to specific stripteamide derivatives, as observed in the *Steinernema hermaphroditum–Xenorhabdus griffiniae* association. The structural diversity of stripteamides, together with their chain-length–dependent bioactivity, raises the possibility that this metabolite family might function as a tunable chemical signaling system coordinating bacterial metabolism with host developmental state.

Furthermore, modulating host development is already the third biological function described for stripteamides besides acting as quorum quenching^29^ and cytotoxic^30^ compounds. This might suggest a general biological principle where natural product function is actually context and interaction partner specific. Here, the generation of variant libraries that easily can adopt different functions might be especially beneficial to the producing strains.

Together, our findings establish stripteamides as conserved symbiotic signals that govern a central developmental transition in *Steinernema* nematodes. More broadly, this work highlights how bacterial secondary metabolism can be co-opted during evolution to mediate communication between symbiotic partners. Understanding such chemical signaling systems will be essential for deciphering the molecular logic of host–microbe interactions and may ultimately enable the rational manipulation of beneficial symbioses for applications such as biological pest control.

## Supporting information

PDB validation report

## Data and material availability

The StaS cryo-EM map and coordinates have been deposited into the Electron Microscopy Data Bank (EMDB) with accession code EMD-58340 and the Protein Data Bank (PDB) with accession code 31EM. All data is shown in the supplementary information and material is available from the corresponding author upon request.

## Acknowledgements

We thank Elmar Meyer for his help with preparing *Galleria* larvae and Tania Köbel for her expert instruction on high-throughput cloning workflows and automated colony handling. Work in the Bode lab was supported by the Max Planck Society and work in the Köhnke lab by Leibniz University Hannover and zukunft.niedersachsen by the Ministerium für Wissenschaft und Kultur. We acknowledge the use of the FBMH Glacios instrument funded by BBSRC grants (BB/T017643/1 & APP61766) and support from the Wellcome Trust for equipment. We also acknowledge access to eBIC KRIOS instruments via BAG BI-36408 sessions 11 & 12.

## Author contributions

H.B.B., L.S., and D.V.B. conceived the study. L.S. identified StaS, validated amide biosynthesis both in vivo and in vitro, generated *X. doucetiae* mutants for nematode development assays, and performed StaS biochemical characterization. D.V.B. conducted nematode development and infection bioassays, established the role of amides in the host nematode, and quantified amide production throughout the infection cycle in *Galleria mellonella*. M.A.D. generated *X. griffiniae* mutants and performed the corresponding nematode development assays. K.P.P. and R.F.C. purified proteins and solved cryo-EM structures together with A.K. and J.K. S.K. performed phylogenetic analyses. K.H. synthesized acyl-SNAC substrate and interpreted NMR data. E.B. established the foundational framework for this study through initial characterization of acyl amide biosynthesis in *X. doucetiae*. R.H.T. and M.T., together with D.V.B. and M.A.D., acquired microscopy movies. L.S., D.V.B., M.A.D., K.P.P., A.K., and S.K. analyzed the data and prepared all figures. L.S. and D.V.B. wrote the manuscript with input from K.P.P., S.K., J.K., and H.B.B. All authors reviewed and approved the final manuscript.

## Declaration of interests

The authors declare no competing interests.

## Notes

### Competing Interest Statement

The authors have declared no competing interest.

## References

1. Lee, K.A., Kim, S.H., Kim, E.K., Ha, E.M., You, H., Kim, B., Kim, M.J., Kwon, Y., Ryu, J.H., and Lee, W.J. (2013). Bacterial-derived uracil as a modulator of mucosal immunity and gut-microbe homeostasis in Drosophila. Cell 153, 797–811. 10.1016/j.cell.2013.04.009.

2. An, D., Oh, S.F., Olszak, T., Neves, J.F., Avci, F.Y., Erturk-Hasdemir, D., Lu, X., Zeissig, S., Blumberg, R.S., and Kasper, D.L. (2014). Sphingolipids from a symbiotic microbe regulate homeostasis of host intestinal natural killer T cells. Cell 156, 123–133. 10.1016/j.cell.2013.11.042.

3. Yang, T., Hu, X., Cao, F., Yun, F., Jia, K., Zhang, M., Kong, G., Nie, B., Liu, Y., Zhang, H., et al. (2025). Targeting symbionts by apolipoprotein L proteins modulates gut immunity. Nature 643, 210–218. 10.1038/s41586-025-08990-4.

4. Zhang, S., Schlabach, K., Perez Carrillo, V.H., Ibrahim, A., Nayem, S., Komor, A., Mukherji, R., Chowdhury, S., Reimer, L., Trottmann, F., et al. (2025). A chemical radar allows bacteria to detect and kill predators. Cell 188, 2495–2504 e2420. 10.1016/j.cell.2025.02.033.

5. Woo, V., and Alenghat, T. (2017). Host-microbiota interactions: epigenomic regulation. Curr Opin Immunol 44, 52–60. 10.1016/j.coi.2016.12.001.

6. Eshleman, E.M., and Alenghat, T. (2021). Epithelial sensing of microbiota-derived signals. Genes Immun 22, 237–246. 10.1038/s41435-021-00124-w.

7. Feng, M., Gao, B., Garcia, L.R., and Sun, Q. (2023). Microbiota-derived metabolites in regulating the development and physiology of Caenorhabditis elegans. Front Microbiol 14, 1035582. 10.3389/fmicb.2023.1035582.

8. He, Y.W., Deng, Y., Miao, Y., Chatterjee, S., Tran, T.M., Tian, J., and Lindow, S. (2023). DSF-family quorum sensing signal-mediated intraspecies, interspecies, and inter-kingdom communication. Trends Microbiol 31, 36–50. 10.1016/j.tim.2022.07.006.

9. Thomas, G.M., and Poinar, G.O. (1979). Xenorhabdus gen. nov., a Genus of Entomopathogenic, Nematophilic Bacteria of the Family Enterobacteriaceae. International Journal of Systematic and Evolutionary Microbiology 29, 352–360. 10.1099/00207713-29-4-352.

10. Cao, M., Patel, T., Rickman, T., Goodrich-Blair, H., and Hussa, E.A. (2017). High levels of the Xenorhabdus nematophila transcription factor Lrp promote mutualism with the Steinernema carpocapsae nematode host. Applied and environmental microbiology 83, e00276–00217.

11. Poinar, G.O. (1975). Description and Biology of a New Insect Parasitic Rhabditoid, Heterorhabditis Bacteriophora N. Gen., N. Sp. (Rhabditida; Heterorhabditidae N. Fam.). Nematologica 21, 463–470. 10.1163/187529275×00239.

12. Koppenhöfer, H.S., and Gaugler, R. (2009). Entomopathogenic nematode and bacteria mutualism. In Defensive mutualism in microbial symbiosis, (CRC Press), pp. 117–134.

13. Stock, S.P., and Blair, H.G. (2008). Entomopathogenic nematodes and their bacterial symbionts: the inside out of a mutualistic association. SYMBIOSIS-REHOVOT- 46, 65.

14. Liao, C.L., Zhang, S.J., Fu, T.T., Jin, X.S., Xu, W.W., Liu, D.X., Zhang, H.M., Liu, Z.M., Liu, X.B., Zhu, T., et al. (2025). Mutualistic bacteria of entomopathogenic nematodes as an insecticidal agent for sustainable agriculture. Chem Biol Technol Ag 12. 10.1186/s40538-025-00862-3.

15. Tarasco, E., Fanelli, E., Salvemini, C., El-Khoury, Y., Troccoli, A., Vovlas, A., and De Luca, F. (2023). Entomopathogenic nematodes and their symbiotic bacteria: from genes to field uses. Front Insect Sci 3, 1195254. 10.3389/finsc.2023.1195254.

16. Snyder, H., Stock, S.P., Kim, S.-K., Flores-Lara, Y., and Forst, S. (2007). New insights into the colonization and release processes of Xenorhabdus nematophila and the morphology and ultrastructure of the bacterial receptacle of its nematode host, Steinernema carpocapsae. Applied and environmental microbiology 73, 5338–5346.

17. Shi, Y.-M., and Bode, H.B. (2018). Chemical language and warfare of bacterial natural products in bacteria–nematode–insect interactions. Natural Product Reports 35, 309–335. 10.1039/C7NP00054E.

18. Dillman, A.R., and Sternberg, P.W. (2012). Entomopathogenic nematodes. Curr Biol 22, R430–431. 10.1016/j.cub.2012.03.047.

19. Joyce, S.A., Brachmann, A.O., Glazer, I., Lango, L., Schwar, G., Clarke, D.J., and Bode, H.B. (2008). Bacterial biosynthesis of a multipotent stilbene. Angew Chem Int Ed Engl 47, 1942–1945. 10.1002/anie.200705148.

20. Golden, J.W., and Riddle, D.L. (1982). A pheromone influences larval development in the nematode Caenorhabditis elegans. Science 218, 578–580.

21. Bode, E., Heinrich, A.K., Hirschmann, M., Abebew, D., Shi, Y.N., Vo, T.D., Wesche, F., Shi, Y.M., Grun, P., Simonyi, S., et al. (2019). Promoter Activation in Deltahfq Mutants as an Efficient Tool for Specialized Metabolite Production Enabling Direct Bioactivity Testing. Angew Chem Int Ed Engl 58, 18957–18963. 10.1002/anie.201910563.

22. Zhou, Q., Grundmann, F., Kaiser, M., Schiell, M., Gaudriault, S., Batzer, A., Kurz, M., and Bode, H.B. (2013). Structure and biosynthesis of xenoamicins from entomopathogenic Xenorhabdus. Chemistry 19, 16772–16779. 10.1002/chem.201302481.

23. Behsaz, B., Bode, E., Gurevich, A., Shi, Y.N., Grundmann, F., Acharya, D., Caraballo-Rodriguez, A.M., Bouslimani, A., Panitchpakdi, M., Linck, A., et al. (2021). Integrating genomics and metabolomics for scalable non-ribosomal peptide discovery. Nat Commun 12, 3225. 10.1038/s41467-021-23502-4.

24. Bode, E., Brachmann, A.O., Kegler, C., Simsek, R., Dauth, C., Zhou, Q., Kaiser, M., Klemmt, P., and Bode, H.B. (2015). Simple “on-demand” production of bioactive natural products. Chembiochem 16, 1115–1119. 10.1002/cbic.201500094.

25. Su, L., Huber, E.M., Westphalen, M., Gellner, J., Bode, E., Kobel, T., Grun, P., Alanjary, M.M., Glatter, T., Cirnski, K., et al. (2024). Isofunctional but Structurally Different Methyltransferases for Dithiolopyrrolone Diversification. Angew Chem Int Ed Engl 63, e202410799. 10.1002/anie.202410799.

26. Park, D., Ciezki, K., van der Hoeven, R., Singh, S., Reimer, D., Bode, H.B., and Forst, S. (2009). Genetic analysis of xenocoumacin antibiotic production in the mutualistic bacterium Xenorhabdus nematophila. Mol Microbiol 73, 938–949. 10.1111/j.1365-2958.2009.06817.x.

27. Bode, H.B., Reimer, D., Fuchs, S.W., Kirchner, F., Dauth, C., Kegler, C., Lorenzen, W., Brachmann, A.O., and Grun, P. (2012). Determination of the absolute configuration of peptide natural products by using stable isotope labeling and mass spectrometry. Chemistry 18, 2342–2348. 10.1002/chem.201103479.

28. Nollmann, F.I., Dauth, C., Mulley, G., Kegler, C., Kaiser, M., Waterfield, N.R., and Bode, H.B. (2015). Insect-specific production of new GameXPeptides in photorhabdus luminescens TTO1, widespread natural products in entomopathogenic bacteria. Chembiochem 16, 205–208. 10.1002/cbic.201402603.

29. Bode, E., He, Y., Vo, T.D., Schultz, R., Kaiser, M., and Bode, H.B. (2017). Biosynthesis and function of simple amides in Xenorhabdus doucetiae. Environ Microbiol 19, 4564–4575. 10.1111/1462-2920.13919.

30. Proschak, A., Schultz, K., Herrmann, J., Dowling, A.J., Brachmann, A.O., ffrench-Constant, R., Muller, R., and Bode, H.B. (2011). Cytotoxic fatty acid amides from Xenorhabdus. Chembiochem 12, 2011–2015. 10.1002/cbic.201100223.

31. Su, L., Huber, E.M., Westphalen, M., Gellner, J., Bode, E., Köbel, T., Grün, P., Alanjary, M.M., Glatter, T., and Cirnski, K. (2024). Isofunctional but structurally different methyltransferases for dithiolopyrrolone diversification. Angewandte Chemie 136, e202410799.

32. Wojda, I. (2017). Immunity of the greater wax moth Galleria mellonella. Insect Sci 24, 342–357. 10.1111/1744-7917.12325.

33. Tobias, N.J., Wolff, H., Djahanschiri, B., Grundmann, F., Kronenwerth, M., Shi, Y.M., Simonyi, S., Grun, P., Shapiro-Ilan, D., Pidot, S.J., et al. (2017). Natural product diversity associated with the nematode symbionts Photorhabdus and Xenorhabdus. Nat Microbiol 2, 1676–1685. 10.1038/s41564-017-0039-9.

34. Kegler, C., and Bode, H.B. (2020). Artificial Splitting of a Non-Ribosomal Peptide Synthetase by Inserting Natural Docking Domains. Angew Chem Int Ed Engl 59, 13463–13467. 10.1002/anie.201915989.

35. Cao, M., Schwartz, H.T., Tan, C.H., and Sternberg, P.W. (2022). The entomopathogenic nematode Steinernema hermaphroditum is a self-fertilizing hermaphrodite and a genetically tractable system for the study of parasitic and mutualistic symbiosis. Genetics 220. 10.1093/genetics/iyab170.

36. Bozhuyuk, K.A.J., Fleischhacker, F., Linck, A., Wesche, F., Tietze, A., Niesert, C.P., and Bode, H.B. (2018). De novo design and engineering of non-ribosomal peptide synthetases. Nat Chem 10, 275–281. 10.1038/nchem.2890.

37. Bozhueyuek, K.A.J., Watzel, J., Abbood, N., and Bode, H.B. (2021). Synthetic Zippers as an Enabling Tool for Engineering of Non-Ribosomal Peptide Synthetases**. Angewandte Chemie-International Edition 60, 17531–17538. 10.1002/anie.202102859.

38. Bloudoff, K., and Schmeing, T.M. (2017). Structural and functional aspects of the nonribosomal peptide synthetase condensation domain superfamily: discovery, dissection and diversity. Biochim Biophys Acta Proteins Proteom 1865, 1587–1604. 10.1016/j.bbapap.2017.05.010.

39. Tanovic, A., Samel, S.A., Essen, L.O., and Marahiel, M.A. (2008). Crystal structure of the termination module of a nonribosomal peptide synthetase. Science 321, 659–663. 10.1126/science.1159850.

40. Yonus, H., Neumann, P., Zimmermann, S., May, J.J., Marahiel, M.A., and Stubbs, M.T. (2008). Crystal structure of DltA. Implications for the reaction mechanism of non-ribosomal peptide synthetase adenylation domains. J Biol Chem 283, 32484–32491. 10.1074/jbc.M800557200.

41. Levichev, A., Faumont, S., Berner, R.Z., Purcell, Z., White, A.M., Chicas-Cruz, K., and Lockery, S.R. (2023). The conserved endocannabinoid anandamide modulates olfactory sensitivity to induce hedonic feeding in C. elegans. Curr Biol 33, 1625–1639 e1624. 10.1016/j.cub.2023.03.013.

