## Supplementary Figures & Tables for "Acyl amides derived from a bacterial symbiont control development of its nematode host"

**Contents**

**Supplementeray Figure 1.** The life cycle of entomopathogenic nematodes.

**Supplementeray Figure 2.** Methodology for entomopathogenic nematode rearing, generation of aposymbiotic and axenic infective juveniles, and in vitro developmental assays.

**Supplementeray Figure 3.** Nature Products synthesized by the wild type of *X. doucetiae.*

**Supplementeray Figure 4.** Stripteamides and StripTRAmides synthesized by the wild type of *X. doucetiae.*

**Supplementeray Figure 5.** Identification of AT38 via targeted AT expression library.

**Supplementeray Figure 6.** Identification of StaS via genomic expression library.

**Supplementeray Figure 7.** StripTRAmide production by StaS-expressing *E. coli*.

**Supplementeray Figure 8.** Production of stripteamides and stripTRAmides during insect infection.

**Supplementeray Figure 9.** Sequence conservation and domain architecture of StaS and related NRPS modules.

**Supplementeray Figure 10.** Biosynthetic pathway and chemical structure of Xefoampeptides (XFPs).

**Supplementeray Figure 11.** Phylogenetic distribution and genomic context of *staS* and *xfpS.*

**Supplementeray Figure 12.** Stripteamides synthesized by the wild type of *X. griffiniae* HGB2511.

**Supplementeray Figure 13.** Production of XFPs and stripteamides by wild type *X. hominickii*.

**Supplementeray Figure 14.** Production of XFPs and stripteamides by wild type *X. bovienii.*

**Supplementeray Figure 15.** Production of XFPs and stripteamides by wild type *X. poinarii.*

**Supplementeray Figure 16.** Production of XFPs and stripteamides by wild type *X. beddingii*.

**Supplementeray Figure 17.** Production of XFPs and stripteamides by wild type *X. vietnamensis*.

**Supplementeray Figure 18.** ^1^H NMR (500 MHz) spectrum of C12 acyl-SNAC in CDCl_3_.

**Supplementeray Figure 19.** ^13^C NMR (126 MHz) spectrum of C12 acyl-SNAC in CDCl_3_.

**Supplementeray Figure 20.** Purification of StaS and XfpS-CAT1.

**Supplementeray Figure 21.** Cryo-EM data processing workflow for StaS.

**Supplementeray Figure 22.** Overlay of StaS size-exclusion chromatograms after reinjecting the monomer and dimer peaks from the first SEC run.

**Supplementeray Figure 23.** Local resolution map of front and back views of StaS.

**Supplementeray Figure 24.** Comparison of StaS structure with AlphaFold3 predicted model.

**Supplementeray Figure 25.** Cryo-EM data processing workflow for XfpS-CAT1.

**Supplementeray Figure 26.** Compariosn of the StaS structure to an XfpS-CAT1 AlphaFold3 predicted model.

**Supplementeray Table 1.** Strains used in this study.

**Supplementeray Table 2.** Plasmides used in this study.

**Supplementeray Table 3.** Primers used in this study.

**Supplementeray Table 4.** Cryo-EM data collection and refinement satatistics.

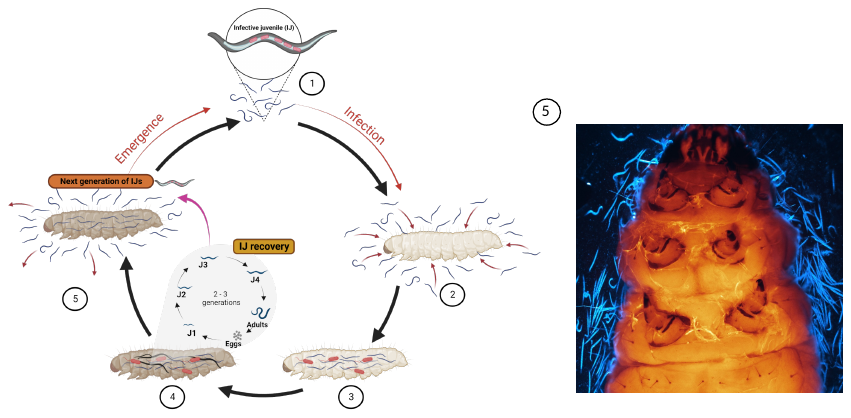

**Supplementary Figure 1**

**The life cycle of entomopathogenic nematodes**. 1). Free-living stage called infective juvenile (IJ) carrying the bacterial symbiont in a non-pathogenic phase within its intestinal tract. 2). IJs locate and enter the insect host through natural openings. 3). Releasing of the symbiont within the hemocoel. 4). Nematode development and bacterial proliferation. 5). The emergence of the new generation of infective juveniles (NG-IJs) from the host cadaver (Created with BioRender.com).

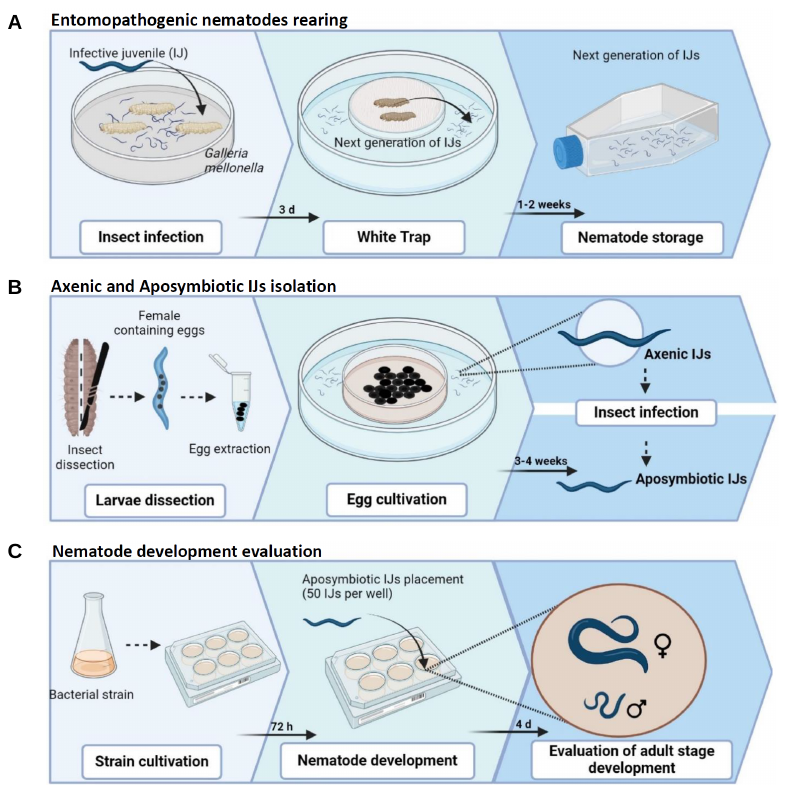

**Supplementary Figure 2**

**Methodology for entomopathogenic nematode rearing, generation of aposymbiotic and axenic infective juveniles, and in vitro developmental assays**. **(A),** Rearing of *Steinernema diaprepesi* and recovery of infective juveniles (IJs) from infected *Galleria mellonella* cadavers using a modified White trap. Insect cadavers displaying characteristic symptoms of *Steinernema* infection (ochre-brown coloration) were selected for IJ collection. **(B)**, Generation of axenic and aposymbiotic nematodes. Eggs were isolated from gravid females of *S. diaprepesi* recovered from infected *G. mellonella* cadavers and cultured on liver–kidney agar to obtain axenic nematodes. Axenic IJs collected using a modified White trap were subsequently used to reinfect *G. mellonella* larvae to generate aposymbiotic IJs. **(C)**, In vitro evaluation of *S. diaprepesi* development. Aposymbiotic IJs were exposed to different bacterial strains, and nematode development was assessed by quantifying progression to first-generation adults.

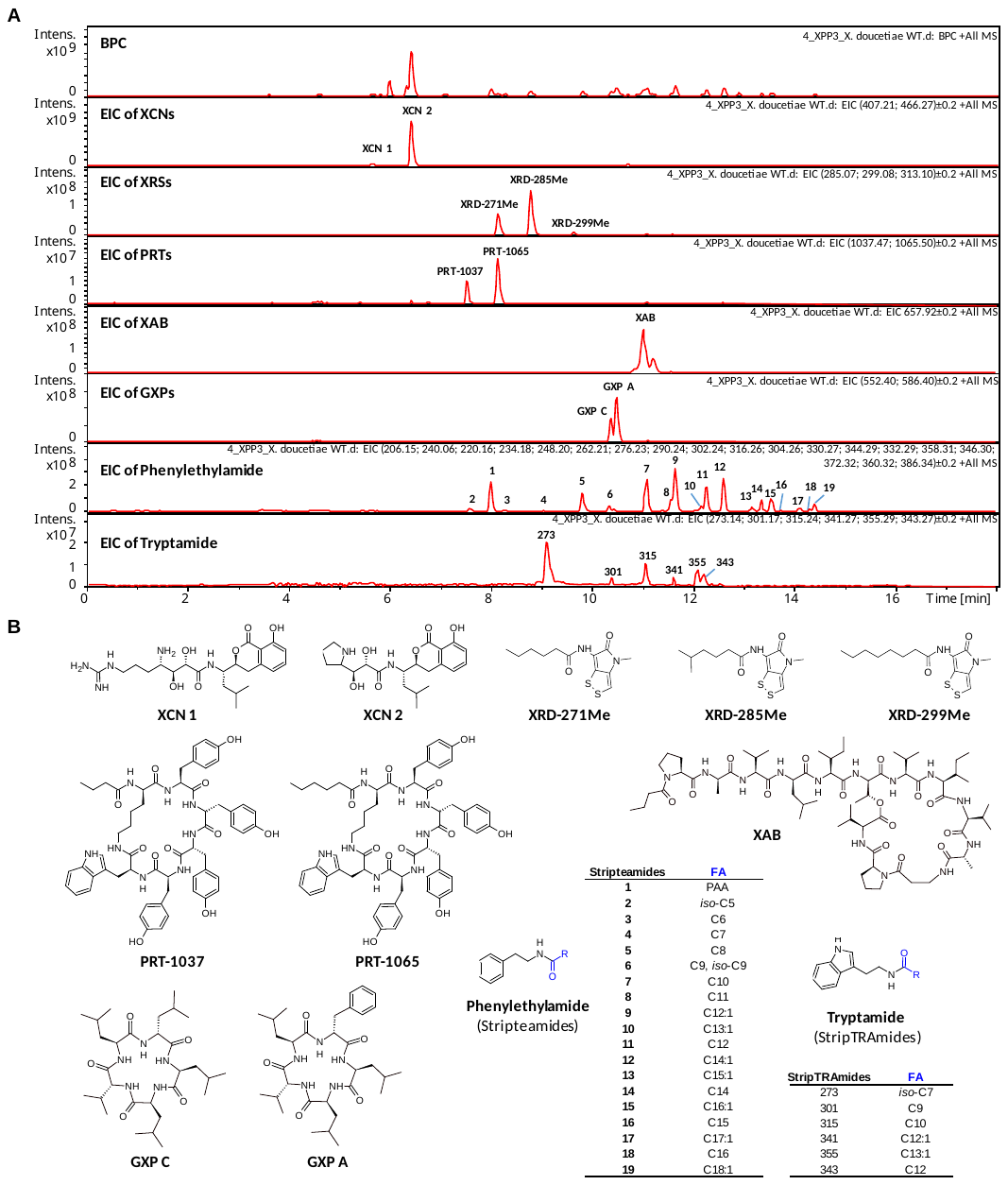

**Supplementary Figure 3**

**Nature Products synthesized by the wild type of *X. doucetiae****.* **(A)**, Base peak chromatogram (BPC) of *X. doucetiae* WT, cultured in XPP3 medium and analyzed by HPLC/MS (amaZon ion trap), together with the extracted ion chromatograms (EICs) for all known natural products (NPs). **(B)**, Structures of these nature products: XCN (Xenocoumacin), XRD (Xenorhabdin), PRT (Protegomycin), XAB (Xenoamicin), GXP (GameXPeptide), and two types of fatty acid amides (Phenylethylamide and Tryptamide).

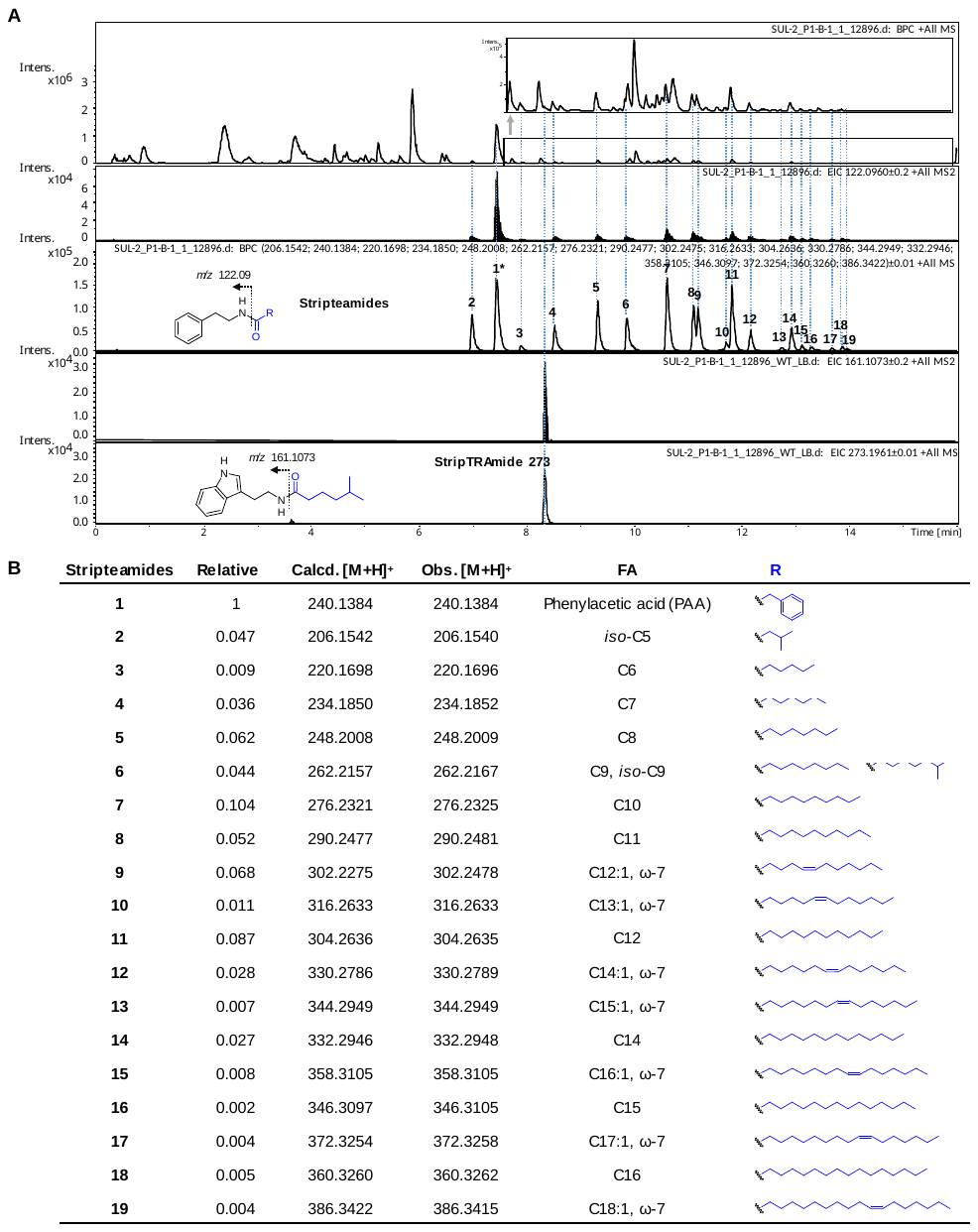

**Supplementary Figure 4**

**Stripteamides and StripTRAmides synthesized by the wild type of *X. doucetiae.*** (**A**) Production of amides by WT of *X. doucetiae* in LB medium analyzed by HPLC/MS using a high-resolution timsTOF mass spectrometer. Compared to the production culture in XPP3 medium, here only stripTRAmide-273 was observed in LB medium condition. (**B**) The structure of stripteamide with the different fatty acid chains is shown in the figure, which has been validated and confirmed by previous studies^1,2^.

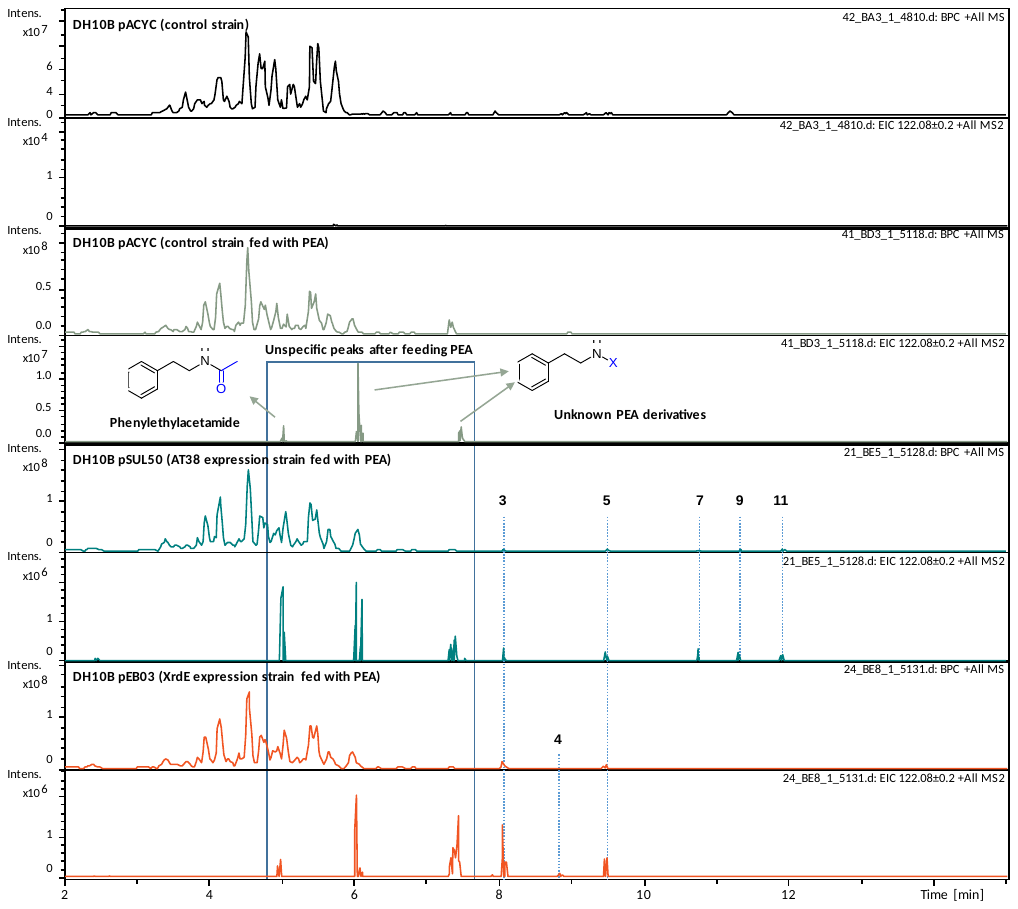

**Supplementary Figure 5**

**Identification of AT38 via targeted AT expression library.** Production of stripteamides by the AT-expression library strains in LB medium analyzed by HPLC/MS (amaZon ion trap). AT candidates (AT1–39) were cloned into the pACYC vector and transformed into *E. coli* DH10B, respectively. The negative control pACYC (empty plasmid) was also transformed into *E. coli* DH10B, which produces phenylethylacetamide and two PEA derivatives of unknown structure. These three compounds were consistently observed in all the AT-expression library strains. The XrdE expression strain DH10B pEB03 serves as a positive control, producing stripteamides **3** (C6), **4** (C7), and **5** (C8). Of the 39 AT candidates, the AT38 expression strain was the only one found to be able to produce stripteamides, including **3** (C6), **5** (C8), **7** (C10), **9** (C12:1), and **11** (C12).

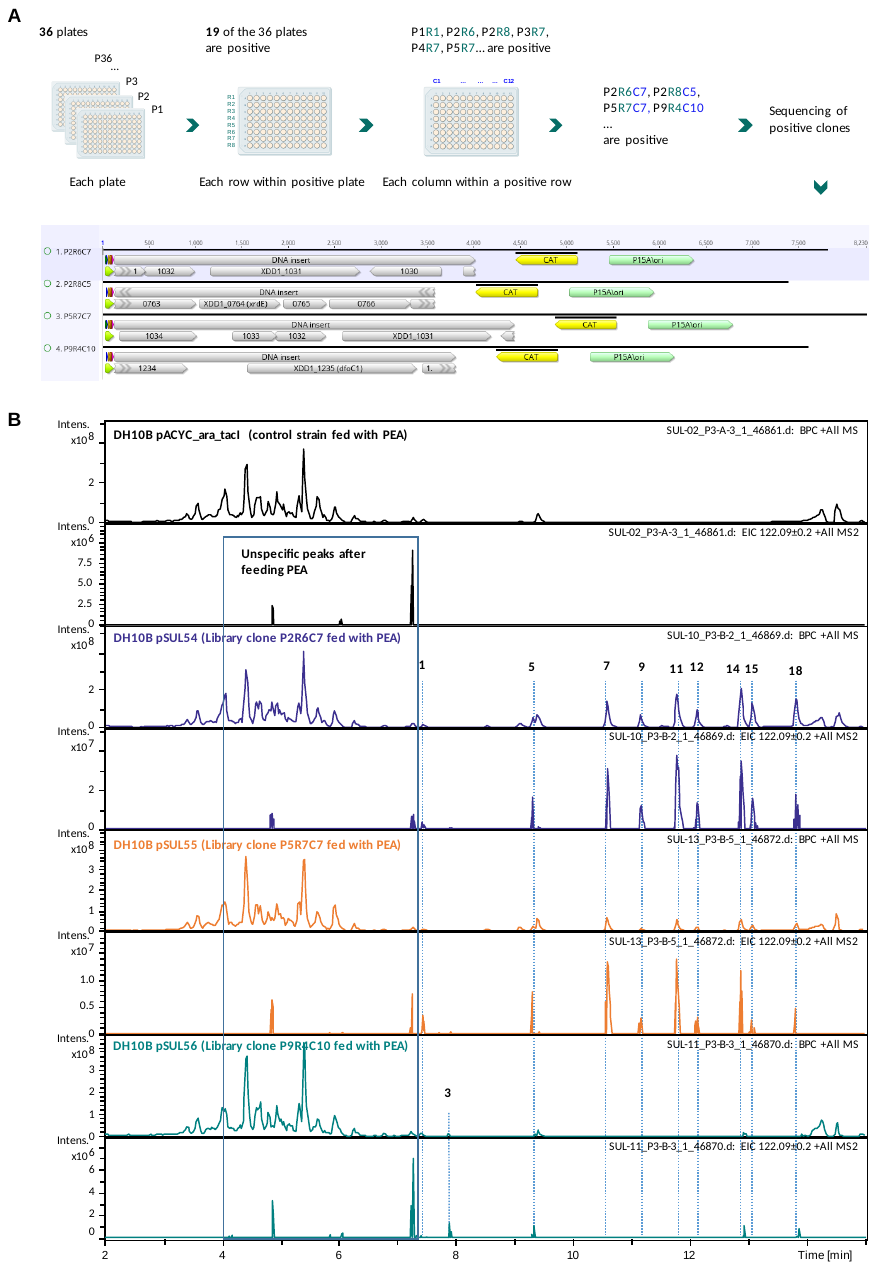

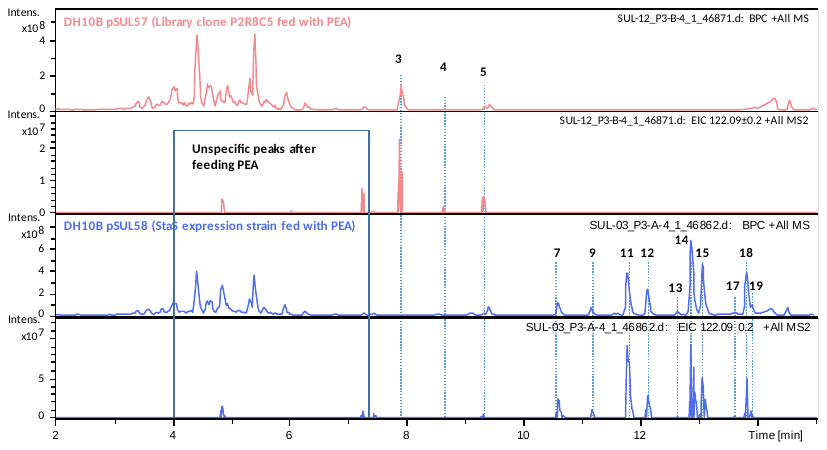

**Supplementary Figure 6**

**Identification of StaS via genomic expression library.** (**A**) The expression library was screened by treating each 96-well plate, each row within a positive plate, and each column within a positive row as a single sample in order to identify positive clones. Four positive clones were identified in the first ten plates. Sequencing revealed that clones P2R6C7 and P5R7C7 contained an identical pair of genes, *XDD1_1031* and *XDD1_1032*; clone P2R8C5 contained XrdE; and clone P9R4C10 contained the gene *XDD1_1235*, which encodes a putative siderophore synthetase (DfoC1) involved in the biosynthesis of the siderophores putrebactin, avaroferrin, and bisucaberin, as reported in our previous study^3^. (**B**) Production of stripteamides by the negative control (empty plasmid pACYC_araC_tacI) clone and four selected positive clones in LB medium and was measured by HPLC/MS amaZon ion trap. Further clone characterization confirmed that the *XDD1_1031* gene encodes protein involved in the biosynthetic pathway for stripteamides, and the protein was then named StaS.

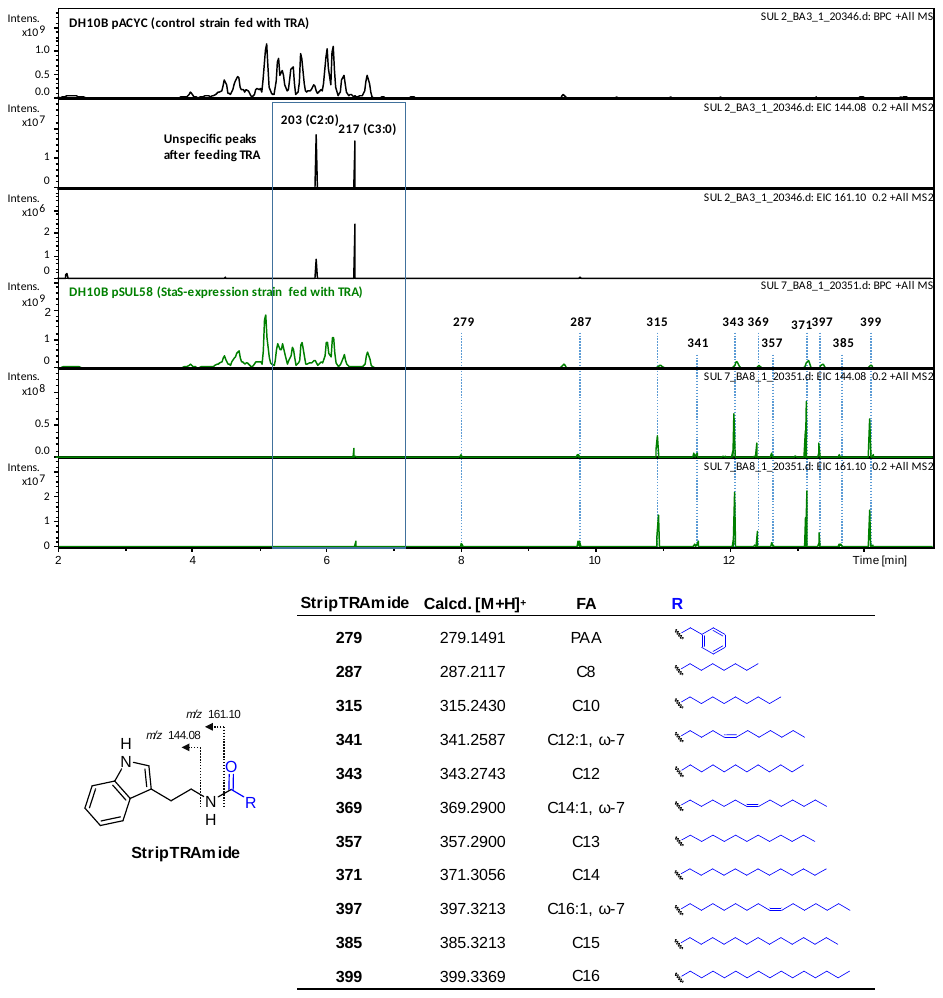

**Supplementary Figure 7**

**StripTRAmide production by StaS-expressing *E. coli*.** Production of stripTRAmides by StaS-expression strain DH10B pSUL58 and the control strain DH10B pACYC in LB medium with amine substrate TRA. Samples were measured by HPLC/MS (amaZon ion trap). According to the previous work^1,2^, structure of these amides is shown with details of the fatty acid chains.

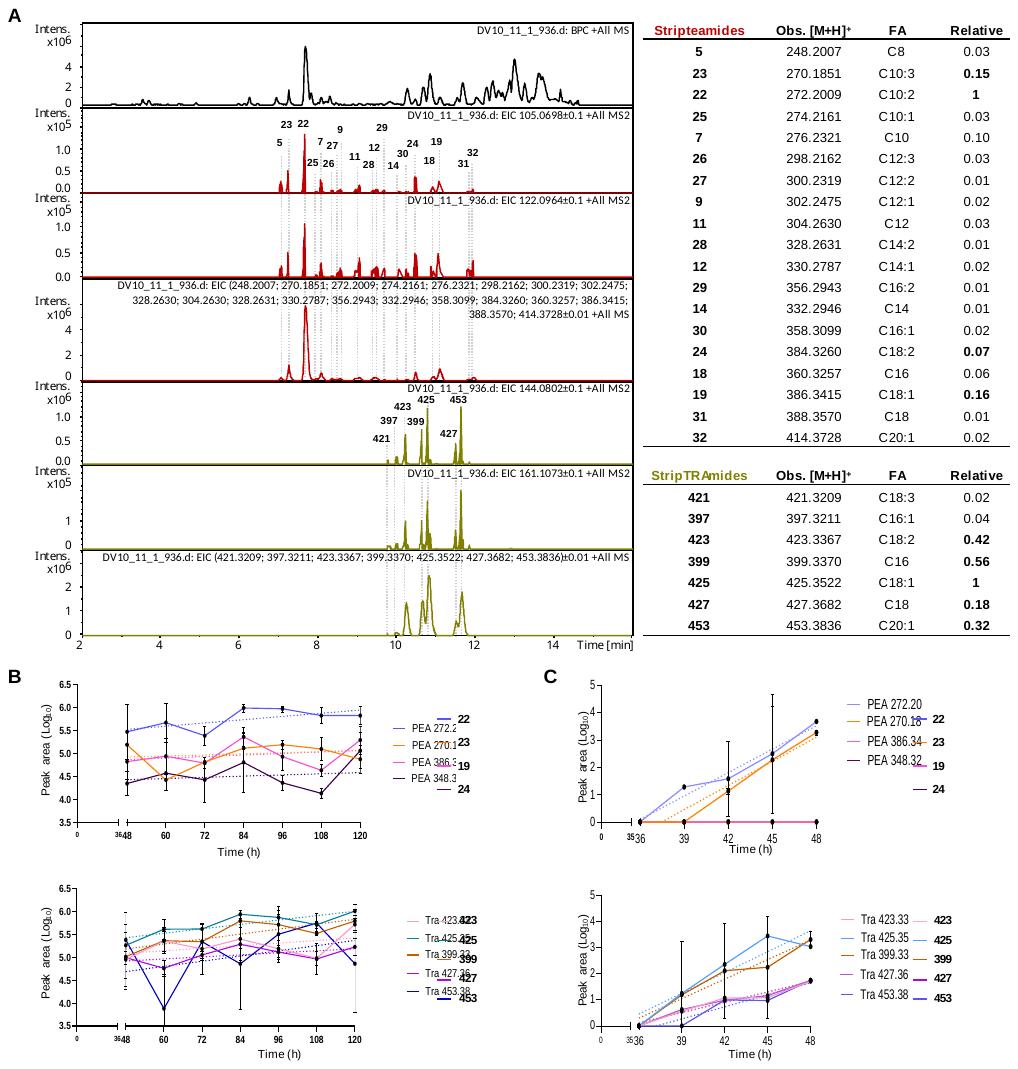

**Supplementary Figure 8**

**Production of stripteamides and stripTRAmides during insect infection.** (**A**), Production profile of stripteamides and stripTRAmides in the hemolymph samples collected from *Galleria mellonella* larvae infected with *S. diaprepesi* IJs carrying the symbiont *X. doucetiae*. Stripteamide is dominated by **22** (C10:2, *m/z* 272.20), **23** (C10:3, *m/z* 270.18), **19** (C18:1, *m/z* 386.34) and **24** (C18:2, *m/z* 384.32). StripTRAmide is dominated by **425** (C18:1), **453** (C20:1), **399** (C16), **423** (C18:2), **427** (C18). Representative chromatogram of a 48h sample (n=3), ratios reflect the mean of three biological replicates. (**B**), Accumulation of the most abundant derivatives of stripteamides and stripTRAmides during insect infection by *S. diaprepesi* over 120 hours. (**C**), Temporal analysis of stripteamide and TRA-amide accumulation during the critical 36–48 h window post-infection, sampled at 3 h intervals. Data in **b** and **c** represent mean peak areas from three independent biological replicates (*n* = 3 per time point). Dashed lines indicate simple linear regression fits.

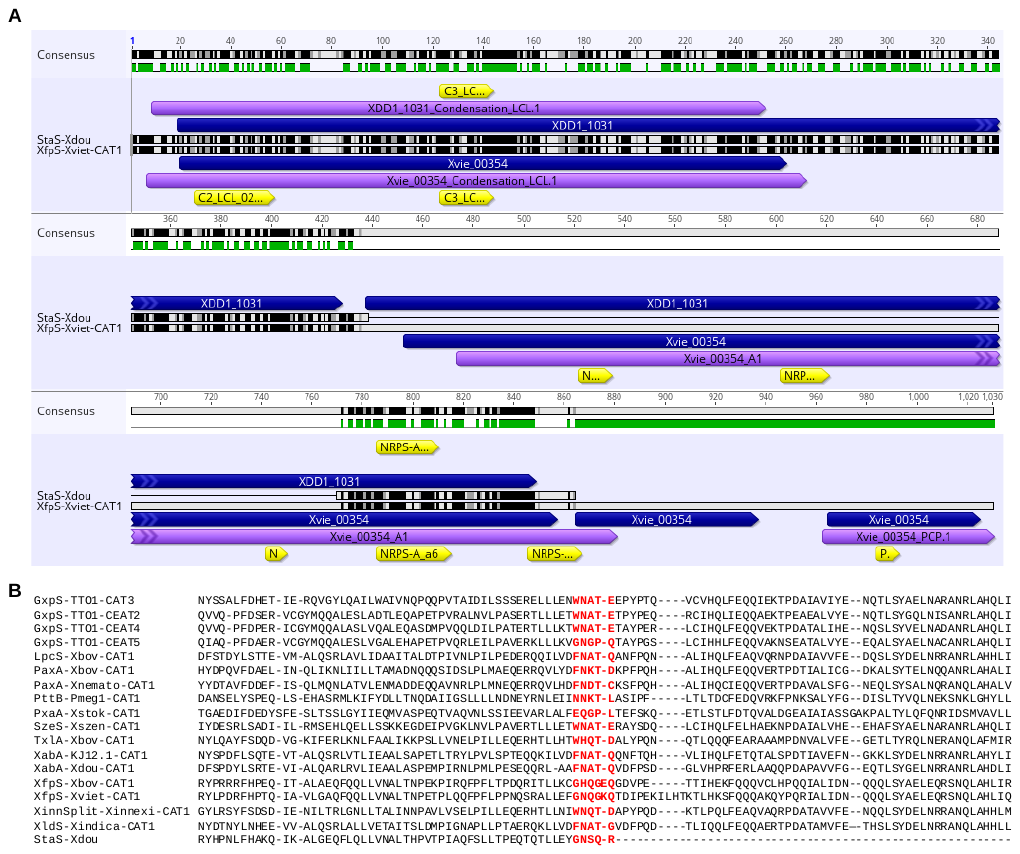

**Supplementary Figure 9**

**Sequence conservation and domain architecture of StaS and related NRPS modules.** (**A**) Sequence alignments of StaS-Xdou and XfpS-Xviet-CAT1 suggest that StaS comprises an NRPS C domain and a partial A domain. (**B**) Sequence alignments of 17 selected NRPS C-A-T and C/E-A-T module sequences from *Photorhabdus* and *Xenorhabdus*. Shown here is part of the alignment including the end of C or E/C domains, the C-A or E/C-A linkers (in bold), and the beginning of the A domains. Second part of the linkers colored in red are the WNATE consensus motif used for C-A splitting. TTO1: *Photorhabdus laumondii* TTO1, Xbov: *Xenorhabdus bovienii* SS-2004, Xnemato: *Xenorhabdus nematophila* ATCC 19061, Pmeg1: *Photorhabdus temperata* Meg1, Xstock: *Xenorhabdus stockiae* DSM 17904, Xszen: *Xenorhabdus szentirmaii* DSM 16338, KJ12.1: *Xenorhabdus sp.* KJ12.1, Xdou: *Xenorhabdus doucetiae* DSM 17909, Xviet: *Xenorhabdus vietnamensis* DSM 22392, Xinnexi: *Xenorhabdus innexi* DSM 16336, Xindica: *Xenorhabdus indica* DSM 17382.

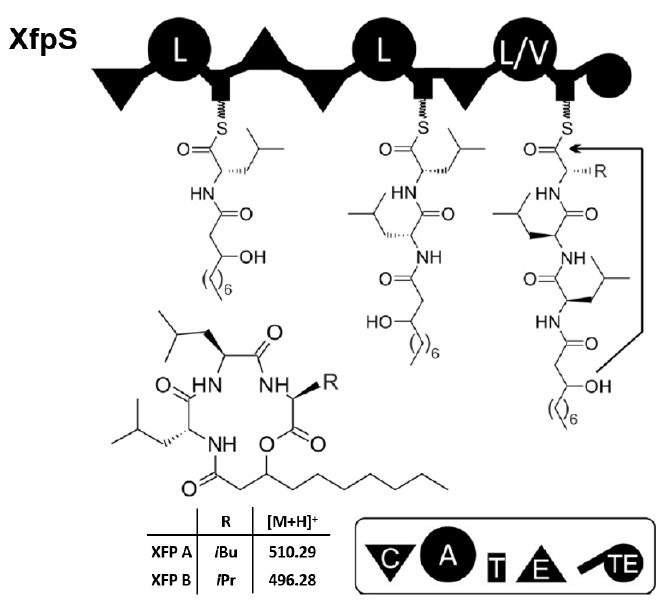

**Supplementary Figure 10**

Biosynthetic pathway and chemical structure of Xefoampeptides (XFPs).^4^

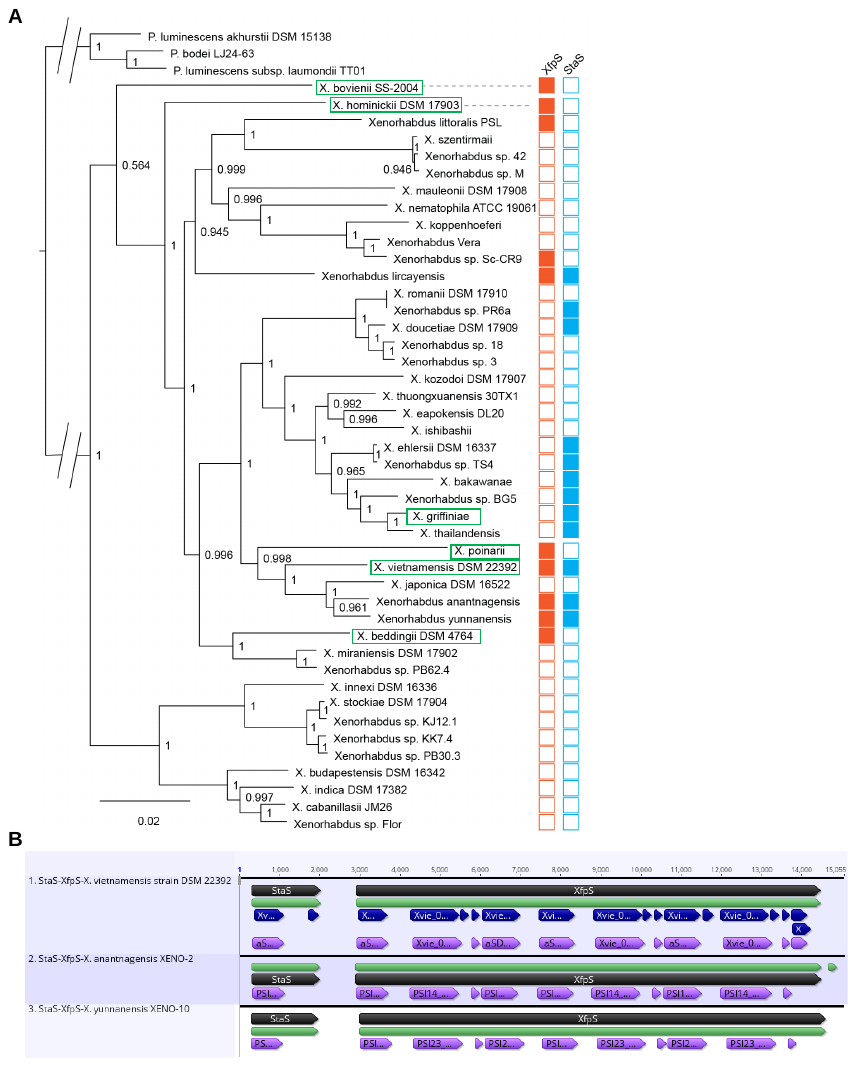

**Supplementary Figure 11**

**Phylogenetic distribution and genomic context of *staS* and *xfpS***. (**A**) Examination of staS and XfpS distribution across the *Xenorhabdus* phylogeny. Strains marked in green boxes were subjected to fermentation to verify the production of stripteamides and xefoampeptides. (**B**) Scheme showing the organization of the *staS* and *xfpS* are located adjacent to each other in strain *X. vietnamensis* DSM 22392 (GenBank: GCA_002127535.1; StaS: Xvie_00355; XfpS: Xvie_00354), *X. anantnagensis* XENO-2 (GenBank: GCA_028598555.1; StaS: PSI14_07380; XfpS: PSI14_07385), and *X. yunnanensis* XENO-10 (GenBank: GCA_028598805.1; StaS: PSI23_15345; XfpS: PSI23_15350).

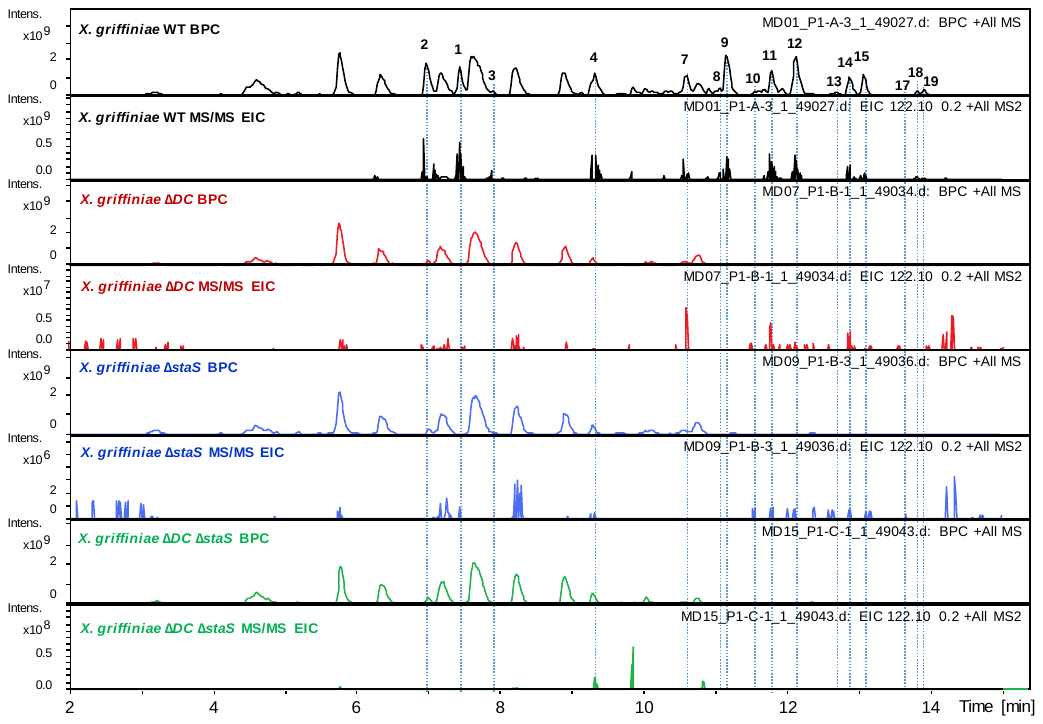

**Supplementary Figure 12**

**Stripteamides synthesized by the wild type of *X. griffiniae* HGB2511***.* WT and thereof mutants were cultured in XPP3 medium and analyzed by HPLC/MS (amaZon io trap). Base peak chromatogram (BPC) and extracted ion chromatograms (EICs) of the diagnostic fragment *m/z* 122 ([M+H]⁺) were checked to confirm the production of stripteamides.

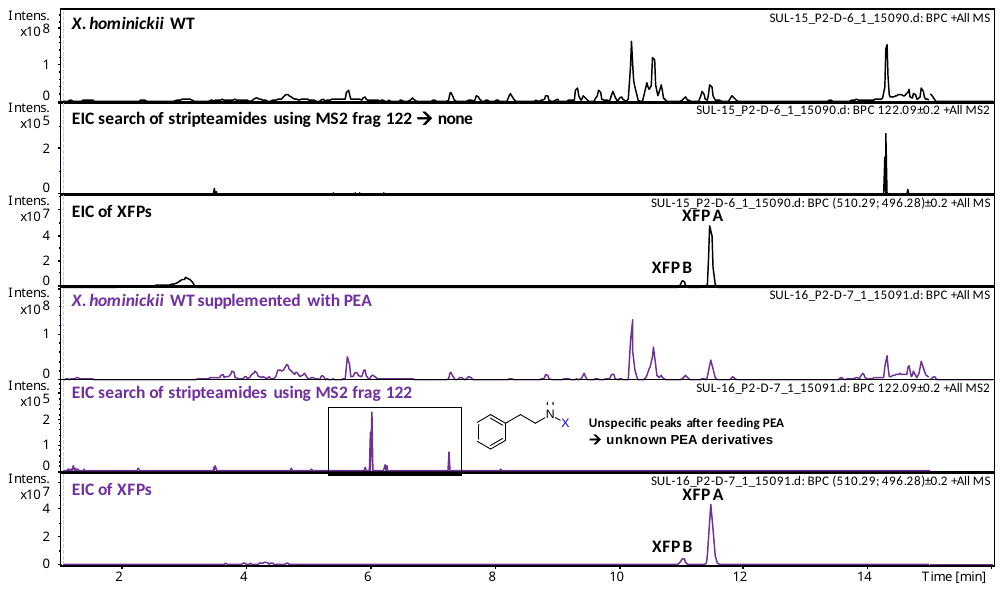

**Supplementary Figure 13**

**Production of XFPs and stripteamides by wild type *X. hominickii*.** Cultivation of *X. hominickii* in XPP3 medium with or without 1 mM PEA supplementation. Data were analyzed by HPLC/MS (amaZon ion trap).

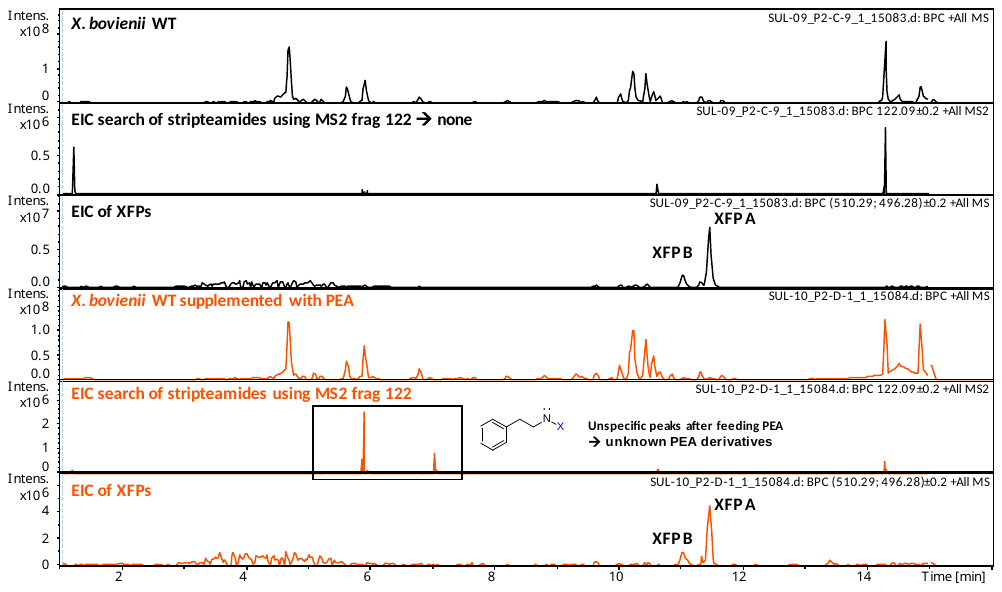

**Supplementary Figure 14**

**Production of XFPs and stripteamides by wild type *X. bovienii*.** Cultivation of *X. bovienii* in XPP3 medium with or without 1 mM PEA supplementation. Data were analyzed by HPLC/MS (amaZon ion trap).

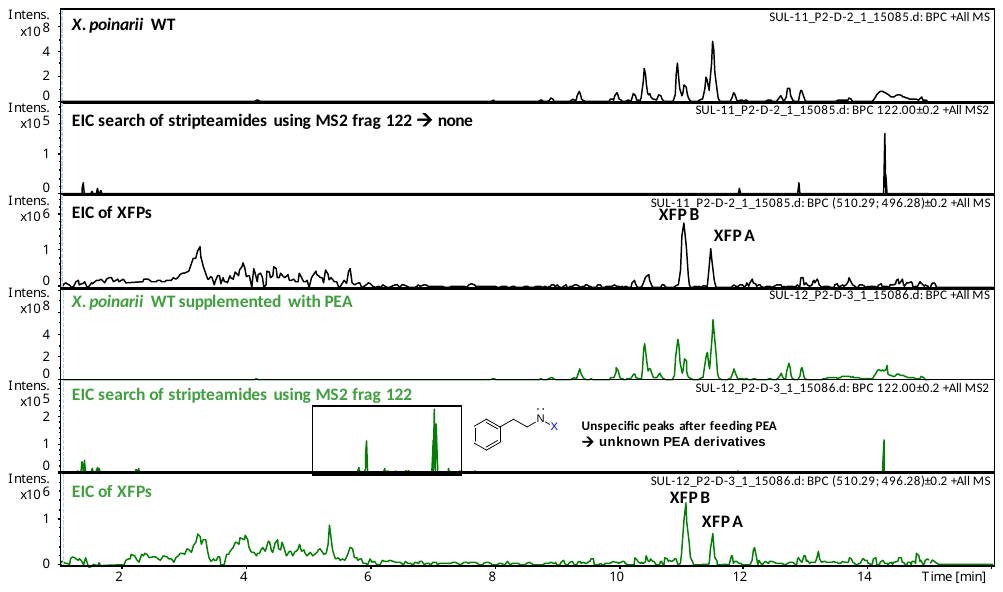

**Supplementary Figure 15**

**Production of XFPs and stripteamides by wild type *X. poinarii*.** Cultivation of *X. poinarii* in XPP3 medium with or without 1 mM PEA supplementation. Data were analyzed by HPLC/MS (amaZon ion trap).

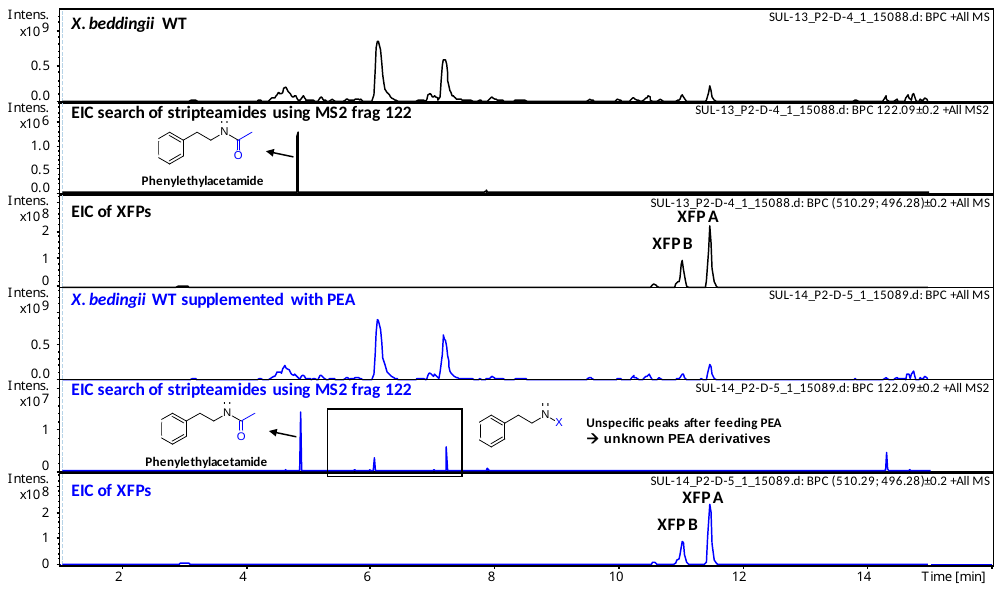

**Supplementary Figure 16**

**Production of XFPs and stripteamides by wild type *X. beddingii*.** Cultivation of *X. beddingii* in XPP3 medium with or without 1 mM PEA supplementation. Data were analyzed by HPLC/MS (amaZon ion trap).

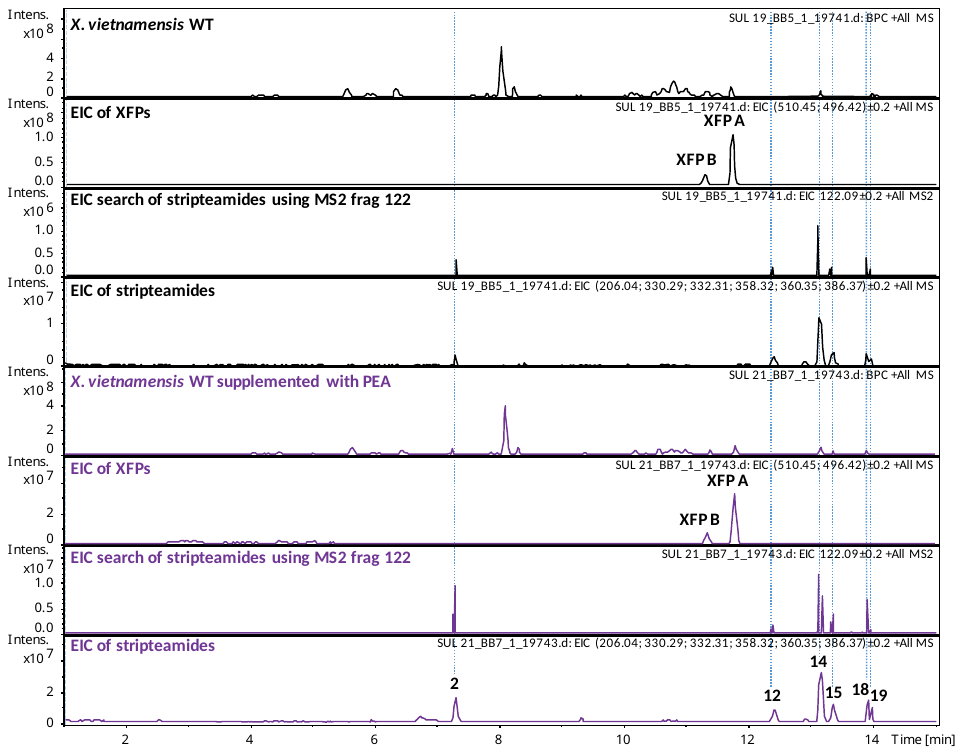

**Supplementary Figure 17**

**Production of XFPs and stripteamides by wild type *X. vietnamensis*.** Cultivation of *X. vietnamensis* in XPP3 medium with or without 1 mM PEA supplementation. Data were analyzed by HPLC/MS (amaZon ion trap).

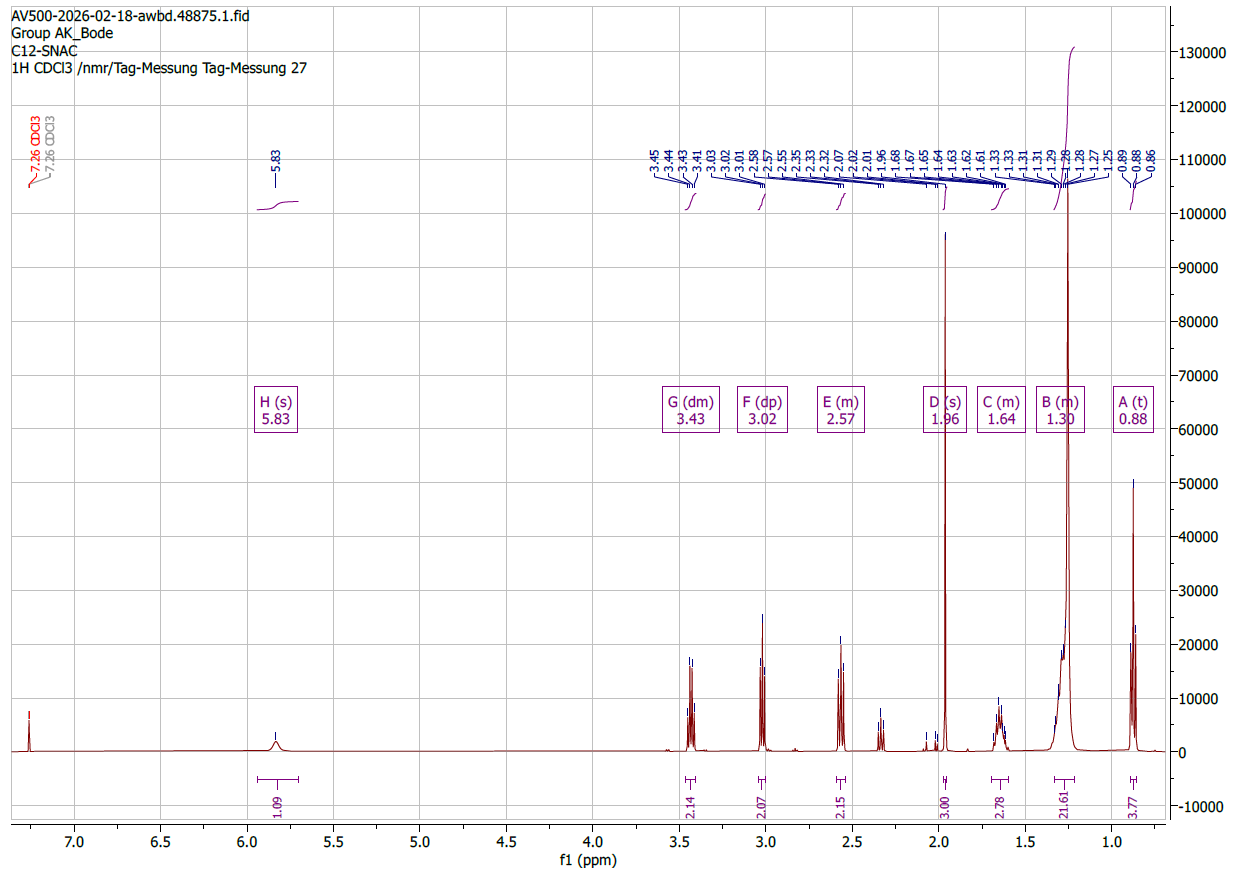
**Supplementary Figure 18**

^1^H NMR (500 MHz) spectrum of C12 acyl-SNAC in CDCl_3_.

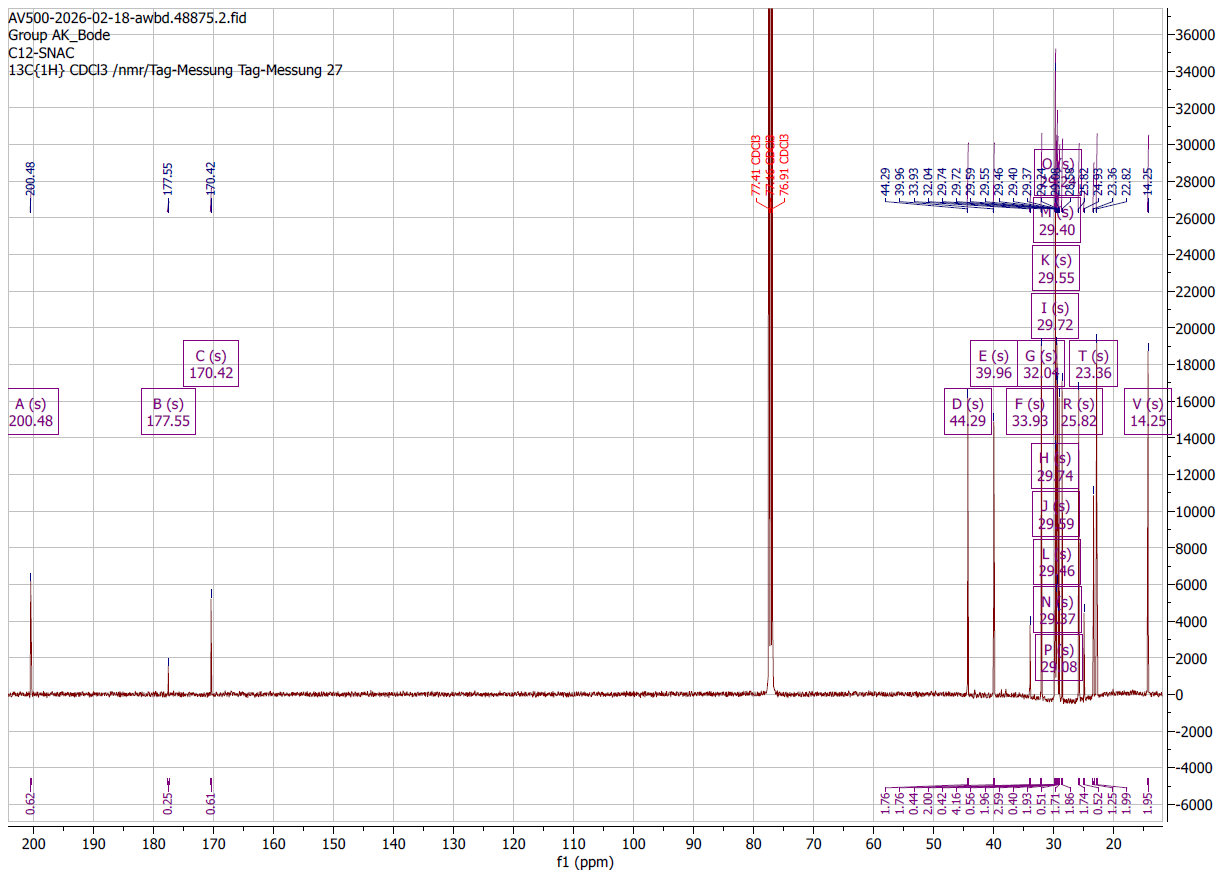

**Supplementary Figure 19**

^13^C NMR (126 MHz) spectrum of C12 acyl-SNAC in CDCl_3_.

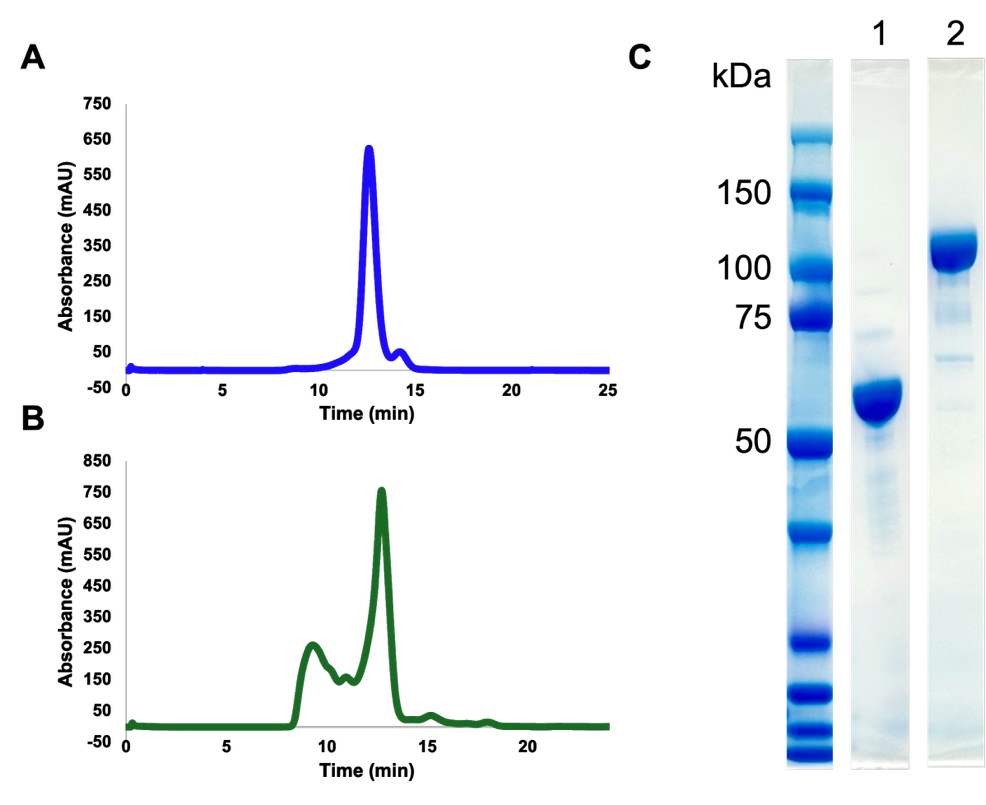

**Supplementary Figure 20**

**Purification of StaS and XfpS-CAT1.** (**A**) Size-exclusion chromatogram of StaS dimer. (**B**) Size-exclusion chromatogram of XfpS. (**C**) SDS-PAGE analysis of (1) StaS and (2) XfpS after size-exclusion chromatography.

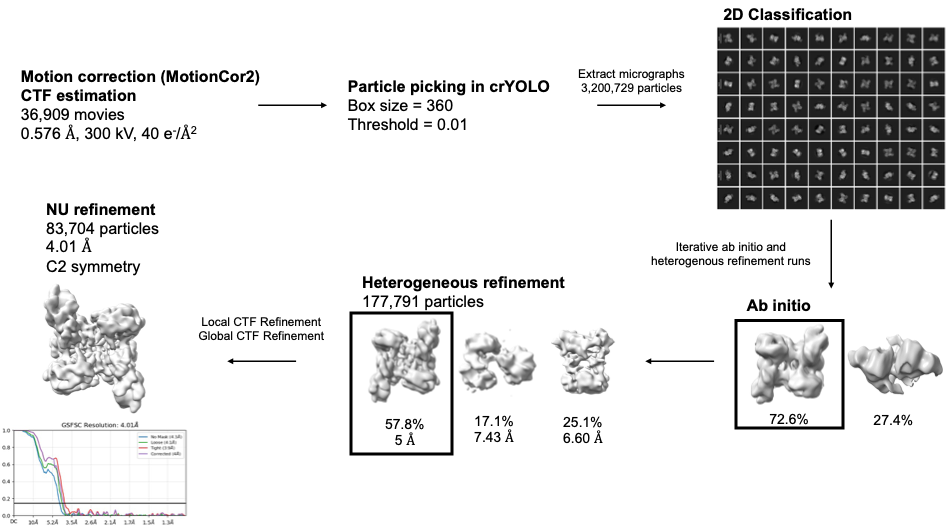

**Supplementary Figure 21**

**Cryo-EM data processing workflow for StaS.** Processing was carried out using crYOLO and cryoSPARC with a total of 36,909 movies collected. The final model contains 83,704 particles at 4.01 Å.

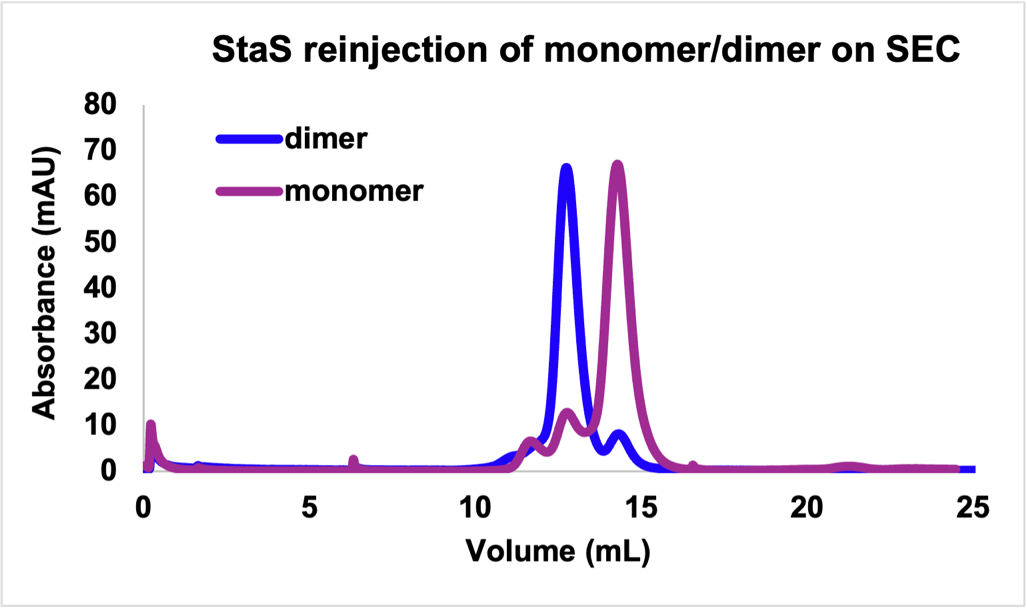

**Supplementary Figure 22**

Overlay of StaS size-exclusion chromatograms after reinjecting the monomer and dimer peaks from the first SEC run.

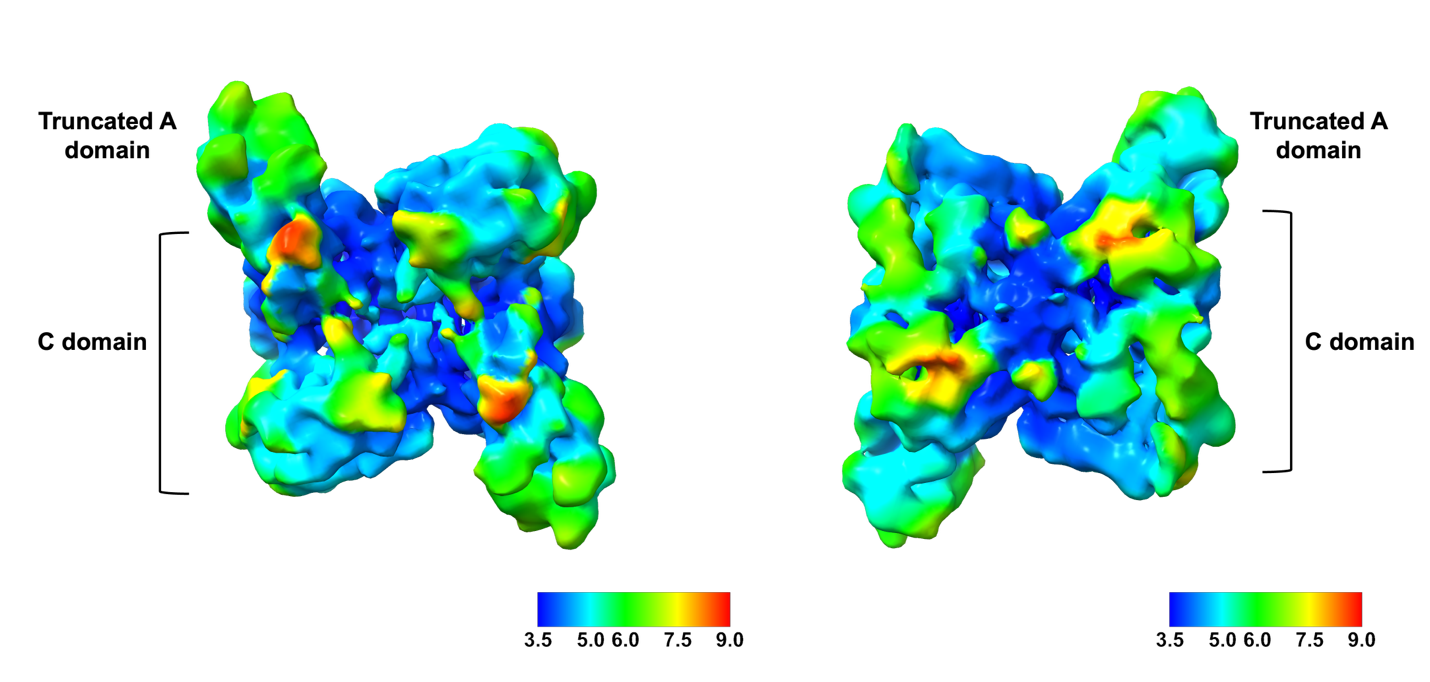

**Supplementary Figure 23**

Local resolution map of front and back views of StaS. Lower resolution areas are observed in the more flexible A domain and N-terminal lobe of the C domain. The color key represents resolution in Å.

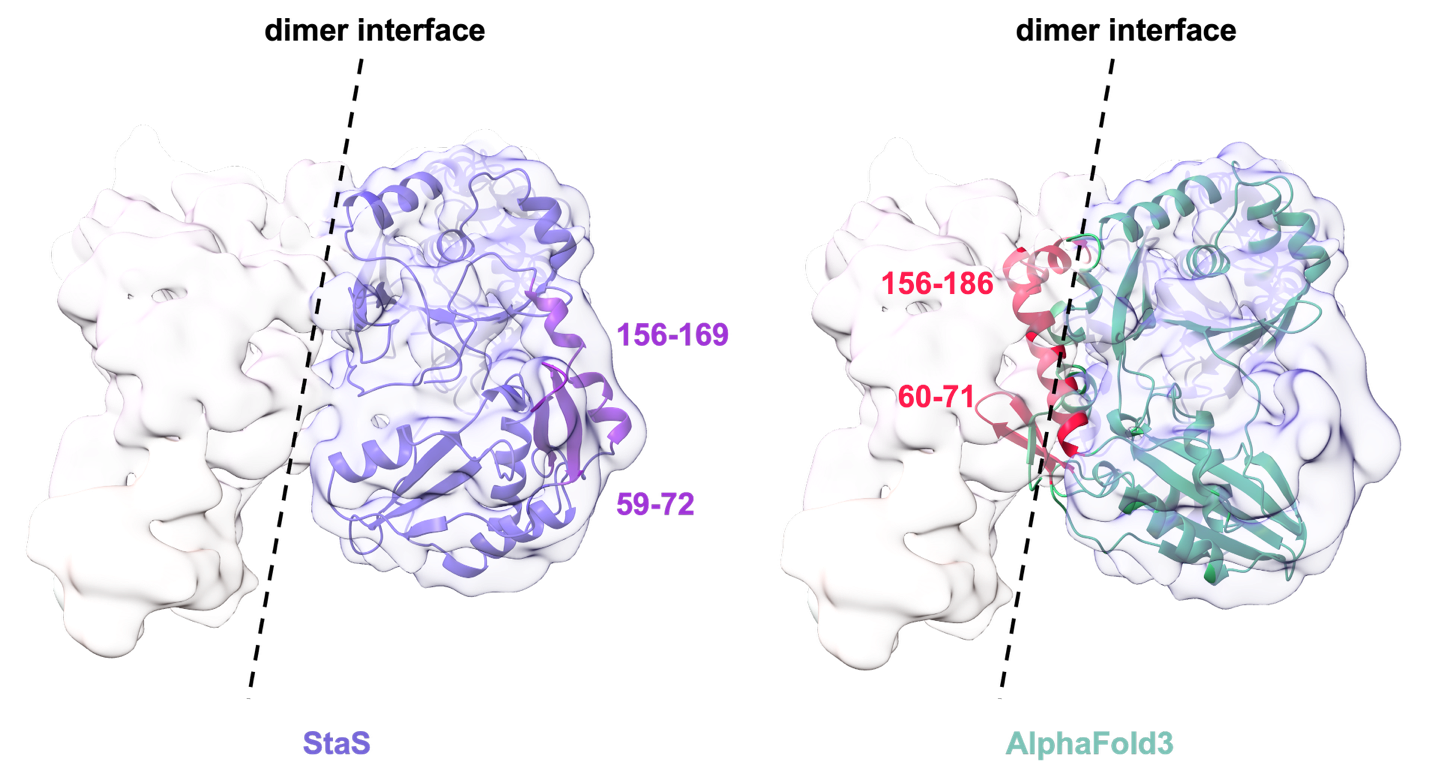

**Supplementary Figure 24**

**Comparison of StaS structure with AlphaFold3 predicted model.** A key difference is observed in residues 59-72 and 156-169, where these secondary structure elements are shifted away from the dimer interface in the solved StaS structure.

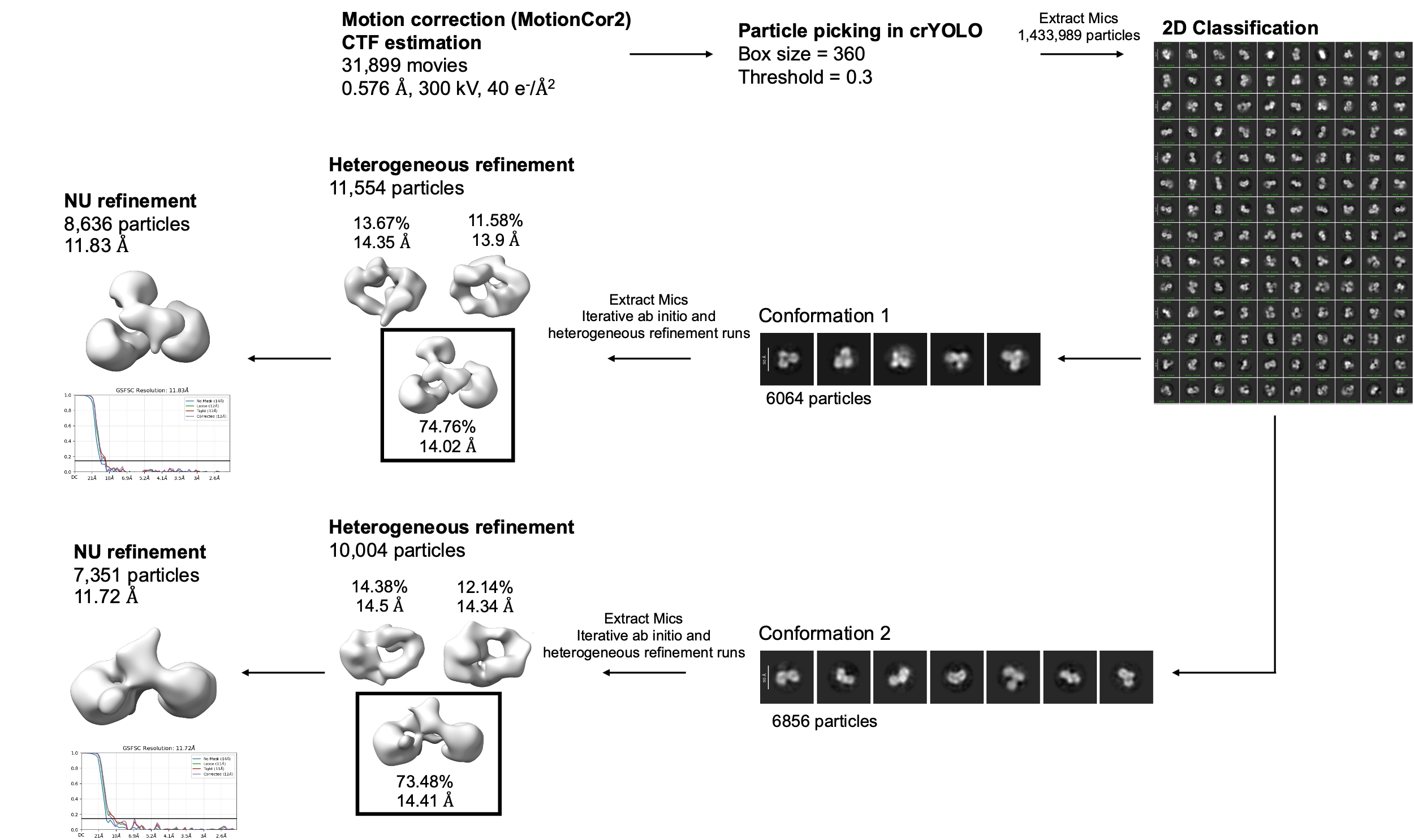

**Supplementary Figure 25**

**Cryo-EM data processing workflow for XfpS-CAT1.** Processing was carried out using crYOLO and cryoSPARC with a total of 31,899 movies. Two conformations of the protein were chosen to generate initial low-resolution models.

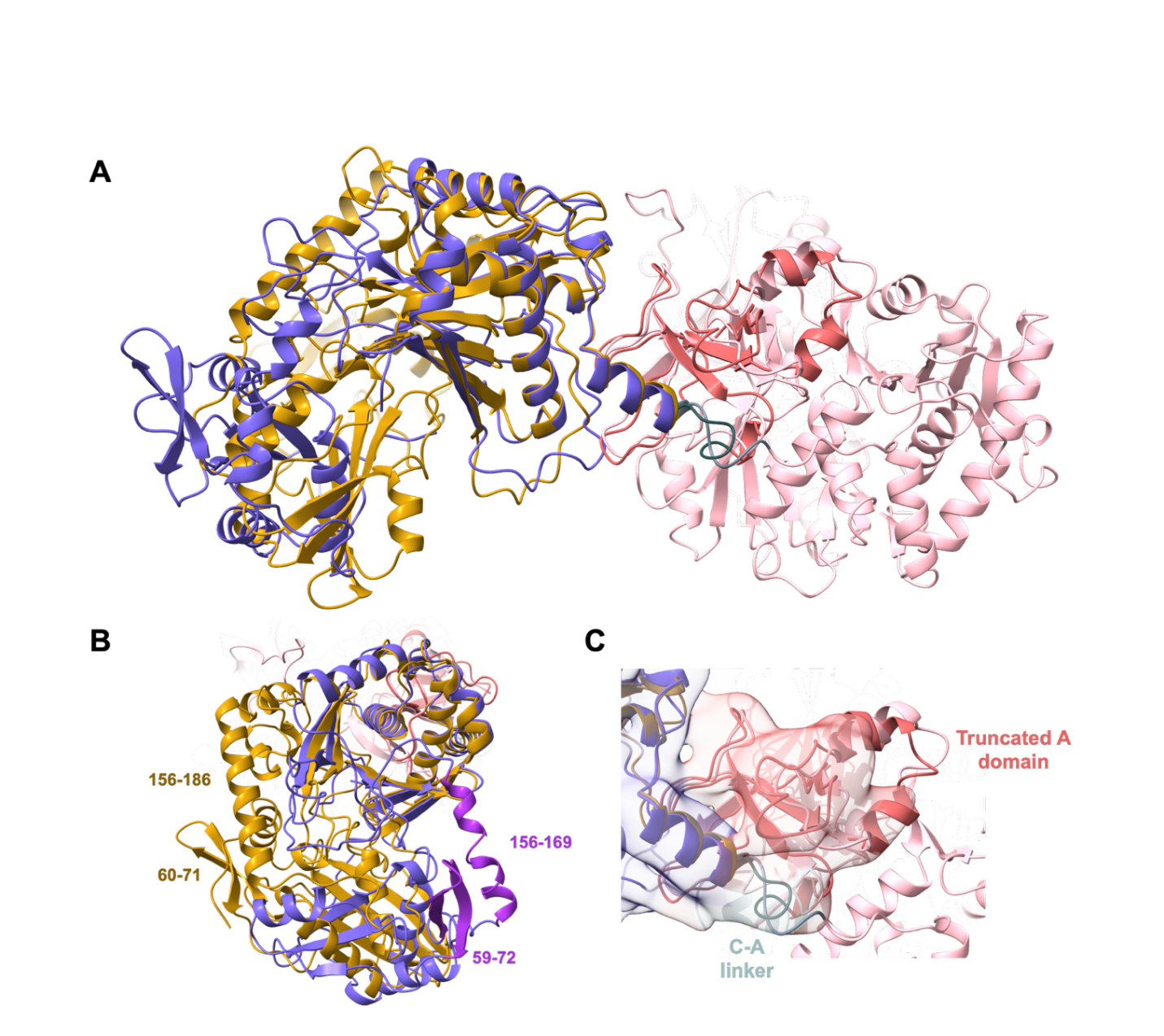

**Supplementary Figure 26**

**Comparison of the StaS structure to an XfpS-CAT1 AlphaFold3 predicted model. (A)**, Comparison of StaS condensation domain (slate blue) and truncated A domain (dark pink) with XfpS AF3 model showing similar architecture overall within the A domain (light pink) and key differences in the C domain (gold). **(B)**, Comparison of the C domains shows a similar rotation of residues 59-72 and 156-169 away from the dimer interface in StaS (purple). **(C)**, Comparison of the truncated A domain of StaS (dark pink) to the full A domain of XfpS (light pink) shows overall similar folding.

**Supplementary Table 1.** Strains used in this study.

| **Strain** | **Genotype/Description** | **Reference** |
| --- | --- | --- |
| ***Xenorhabdus doucetiae* DSM 17909**  **(NCBI reference sequence)** |  |  |
| WT | *X. doucetiae* DSM 17909 wild type | Ref^2^ |
| WT pCEP_XAB | X. doucetiae DSM 17909 with the P*_BAD_* promoter exchanged in front of gene *xabA* (*XDv3_70458*) | Ref^5^ |
| WT pCEP_PRT | X. doucetiae DSM 17909 with the P*_BAD_* promoter exchanged in front of gene *prtA* *(XDv3_70082)* | Ref^5^ |
| WT pCEP_XRD | X. doucetiae DSM 17909 with the P*_BAD_* promoter exchanged in front of gene *xrdA* *(XDv3_10495)* | Ref^5^ |
| WT pCEP_XCN | X. doucetiae DSM 17909 with the P*_BAD_* promoter exchanged in front of gene *xcnA* (*XDD1_RS08255)* | Ref^6^ |
| WT pCEP_GXP | X. doucetiae DSM 17909 with the P*_BAD_* promoter exchanged in front of gene *gxpS* (*XDv3_110151)* | Ref^7^ |
| WT pCEP_DC | X. doucetiae DSM 17909 with the PBAD promoter exchanged in front of gene *DC* (*XDD1_2132*) | Ref^2^ |
| Δ*DC* (WP_045970772.1) | *X. doucetiae* DSM 17909 with the deletion of gene encoding DC (*XDD1_2132*) | Ref^2^ |
| Δ*xrdE* (WP_323857116.1) | *X. doucetiae* DSM 17909 with the deletion of *xrdE* (*XDD1_0766*) | Ref^2^ |
| Δ*AT38* (WP_045969499.1) | *X. doucetiae* DSM 17909 with the deletion of *AT38* (*XDD1_1245*) | This study |
| Δ*xrdE* Δ*AT38* | Δ*xrdE* with the deletion of *AT38* | This study |
| Δ*staS* (WP_045969165.1) | *X. doucetiae* DSM 17909 with the deletion of gene encoding *staS* (*XDD1_1031*) | This study |
| Δ*xrdE* Δ*staS* | Δ*xrdE* with the deletion of *staS* | This study |
| Δ*AT3*8 Δ*staS* | Δ*AT38* with the deletion of *staS* | This study |
| Δ*triple* | Δ*xrdE* Δ*AT38* with the deletion of *staS* | This study |
| Δ*tetra* | Δ*xrdE* Δ*AT38* Δ*staS* with the deletion of gene encoding DC | This study |
| ***Xenorhabdus griffiniae* HGB2511** |  |  |
| WT | *X. griffiniae* HGB2511 wild type | This study |
| Δ*DC_griff_* (WXG15772.1) | *X. griffiniae* HGB2511 with the deletion of gene encoding DC (*WDV76_10420*) | This study |
| Δ*staS_griff_* (WXG12962.1) | *X. griffiniae* HGB2511 with the deletion of gene encoding amide synthetase (*WDV76_15415*) | This study |
| Δ*DC_griff_* Δ*staS_griff_* | Δ*DC* with the deletion of *staS* | This study |
| ***Other Xenorhabdus species*** |  |  |
| *X. hominickii* | *Xenorhabdus hominickii* DSM 17903 | Ref^8^ |
| *X. bovienii* | *Xenorhabdus bovienii* SS-2004 | Ref^8^ |
| *X. poinarii* | *Xenorhabdus poinarii* DSM 4768 | Ref^9^ |
| *X. beddingii* | *Xenorhabdus beddingii* DSM 4764 | Ref^8^ |
| *X. vietnamensis* | *Xenorhabdus vietnamensis* DSM22392 | Ref^8^ |
| ***E. coli*** |  |  |
| DH10B | Cloning and expression strain; F^–^ *mcr*A, Δ(*mrr-hsdRMS-mcrBC*) Φ80*lacZ* Δ*M15* Δ*lac*X74 *rec*A1 *end*A1,araD139, Δ(*ara-leu*)7697 *gal*U, *gal*K, *rps*L,*nup*G, λ- | Invitrogen |
| DH10B::*mtaA* | Expression strain; DH10B with *mtaA* from pCK_*mtaA* Δ*entD* | Ref^10^ |
| DH5α | Cloning strain; *fhuA2::IS2 Δ(mmuP-mhpD)169 ΔphoA8 glnX44* *ϕ80d[ΔlacZ58(M15)] rfbD1 gyrA96 luxS11 recA1 endA1 rphWT thiE1 hsdR17* | NEB |
| BL21(DE3) | Expression strain; F– *omp*T *hsdS_B_* (r_B_–, m_B_–) *gal dcm* (DE3) | Novagen |
| ST18 | Conjugation strain; *E. coli* S17-1 *λpir* Δ*hem*A | Ref^11^ |
| ST18 pEB_de-AT38 | ST18 contains the gene *AT38* deletion plasmid pEB_de-AT38 | This study |
| ST18 pEB_de-staS | ST18 contains the gene *staS* deletion plasmid pEB_de-staS | This study |
| ST18 pEB_de-DC | ST18 contains the gene *DC* deletion plasmid pEB_de-DC | Ref^2^ |
| ST18 pEB_de-staS_griff_ | ST18 contains the gene *staS* deletion plasmid pEB_de-staS_griff_ | This study |
| ST18 pEB_de-DC_griff_ | ST18 contains the gene *DC* deletion plasmid pEB_de-DC_griff_ | This study |

**Supplementary Table 2.** Plasmids used in this study.

| **Plasmid** | **Genotype/Description** | **Reference** |
| --- | --- | --- |
| pACYC | p15A ori, *araC*-P*_BAD_*, *tacI*, Cm^R^ | Ref^12^ |
| pACYC_SZ17 | pACYC derivative with the SYNZIP17 sequence attached after the target gene entry site | Ref^13^ |
| pCOLA | ColA ori, *araC*-P*_BAD_*, *tacI*, Kana^R^ | Ref^12^ |
| pCOLA_SZ18 | pACYC derivative with the SYNZIP18 sequence attached ahead of the target gene entry site | Ref^13^ |
| pCDF | CloDF13 ori, *araC*-P*_BAD_*, *tacI*, Spec^R^ | Ref^12^ |
| pEB17 | pDS132 based, R6K ori, oriT, *cipB* derivative with additional *Bgl*II site, *sacB,* kan^R^ | Ref^14^ |
| pEB02 | pEB17 backbone with and two homologous arms (HAL and HAR) of gene *xrdE* were ligated by Gibson assembly | Ref^2^ |
| pEB_de-AT38 | pEB17 backbone with and two homologous arms (HAL and HAR) of gene *AT38* were ligated by Gibson assembly | This study |
| pEB_de*-*DC | pEB17 backbone with and two homologous arms (HAL and HAR) of gene *DC* were ligated by Gibson assembly | This study |
| pEB_de-staS | pEB17 backbone with and two homologous arms (HAL and HAR) of gene *staS* were ligated by Gibson assembly | This study |
| pEB_de-DC_griff_ | pEB17 backbone with and two homologous arms (HAL and HAR) of gene *DC_griff_* were ligated by Gibson assembly | This study |
| pEB_de-staS_griff_ | pEB17 backbone with and two homologous arms (HAL and HAR) of gene *staS_griff_* were ligated by Gibson assembly | This study |
| pCEP_Km | pDS132 based, R6K ori, oriT, *araC*-P*_BAD_*, Kana^R^ | Ref^7^ |
| pEB06 (pCEP_DC) | pCEP-Km with the first 793 bp region of *XDD1_RS09835* (*DC*) was inserted under araBAD promoter | Ref^7^ |
| pCEP_GXP | pCEP-Km, *XDv3_11015* | Ref^7^ |
| pCEP_XCN | pCEP-Km, *XDD1_RS08255* | Ref^6^ |
| pCEP_PRT | pCEP-Km, *XDv3_70082* | Ref^5^ |
| pCEP_XAB | pCEP-Km, *XDv3_70458* | Ref^7^ |
| pCEP_XRD | pCEP-Km, *XDv3_10499* | Ref^7^ |
| pSUL14 | pACYC with gene *AT2* (*XDD1_2637*, PCR amplified using sul03+04) inserted under P*_BAD_* control | This study |
| pSUL15 | pACYC with gene *AT3* (*XDD1_1046*, PCR amplified using sul05+06) inserted under P*_BAD_* control | This study |
| pSUL16 | pACYC with gene *AT4* (*XDD1*_*0723*, PCR amplified using sul07+08) inserted under P*_BAD_* control | This study |
| pSUL17 | pACYC with gene *AT5* (*XDD1*_*1664*, PCR amplified using sul09+10) inserted under P*_BAD_* control | This study |
| pSUL18 | pACYC with gene *AT6* (*XDD1*_*3811*, PCR amplified using sul11+12) inserted under P*_BAD_* control | This study |
| pSUL19 | pACYC with gene *AT7* (*XDD1*_*2739*, PCR amplified using sul13+14) inserted under P*_BAD_* control | This study |
| pSUL20 | pACYC with gene *AT8* (*XDD1*_*3477*, PCR amplified using sul15+16) inserted under P*_BAD_* control | This study |
| pSUL21 | pACYC with gene *AT9* (*XDD1*_*2896*, PCR amplified using sul17+18) inserted under P*_BAD_* control | This study |
| pSUL22 | pACYC with gene *AT10* (*XDD1*_*2147*, PCR amplified using sul19+20) inserted under P*_BAD_* control | This study |
| pSUL23 | pACYC with gene *AT11* (*XDD1*_*0494*, PCR amplified using sul21+22) inserted under P*_BAD_* control | This study |
| pSUL24 | pACYC with gene *AT12* (*XDD1*_*0352*, PCR amplified using sul23+24) inserted under P*_BAD_* control | This study |
| pSUL25 | pACYC with gene *AT13* (*XDD1*_*3888*, PCR amplified using sul25+26) inserted under P*_BAD_* control | This study |
| pSUL26 | pACYC with gene *AT14* (*XDD1*_*1432*, PCR amplified using sul27+28) inserted under P*_BAD_* control | This study |
| pSUL27 | pACYC with gene *AT15* (*XDD1*_*0748*, PCR amplified using sul29+30) inserted under P*_BAD_* control | This study |
| pSUL28 | pACYC with gene *AT16* (*XDD1*_*3418*, PCR amplified using sul31+32) inserted under P*_BAD_* control | This study |
| pSUL29 | pACYC with gene *AT17* (*XDD1*_*0537*, PCR amplified using sul33+34) inserted under P*_BAD_* control | This study |
| pSUL30 | pACYC with gene *AT18* (*XDD1*_*2430*, PCR amplified using sul35+36) inserted under P*_BAD_* control | This study |
| pSUL31 | pACYC with gene *AT19* (*XDD1*_*0160*, PCR amplified using sul37+38) inserted under P*_BAD_* control | This study |
| pSUL32 | pACYC with gene *AT20* (*XDD1*_*0083*, PCR amplified using sul39+40) inserted under P*_BAD_* control | This study |
| pSUL33 | pACYC with gene *AT21* (*XDD1*_*1613*, PCR amplified using sul41+42) inserted under P*_BAD_* control | This study |
| pSUL34 | pACYC with gene *AT22* (*XDD1*_*0504*, PCR amplified using sul43+44) inserted under P*_BAD_* control | This study |
| pSUL35 | pACYC with gene *AT23* (*XDD1*_*0527*, PCR amplified using sul45+46) inserted under P*_BAD_* control | This study |
| pSUL36 | pACYC with gene *AT24* (*XDD1*_*2993*, PCR amplified using sul47+48) inserted under P*_BAD_* control | This study |
| pSUL37 | pACYC with gene *AT25* (*XDD1*_*0706*, PCR amplified using sul49+50) inserted under P*_BAD_* control | This study |
| pSUL38 | pACYC with gene *AT26* (*XDD1*_*3359*, PCR amplified using sul51+52) inserted under P*_BAD_* control | This study |
| pSUL39 | pACYC with gene *AT27* (*XDD1*_*0017*, PCR amplified using sul53+54) inserted under P*_BAD_* control | This study |
| pSUL40 | pACYC with gene *AT28* (*XDD1*_*2992*, PCR amplified using sul55+56) inserted under P*_BAD_* control | This study |
| pSUL41 | pACYC with gene *AT29* (*XDD1*_*2412*, PCR amplified using sul57+58) inserted under P*_BAD_* control | This study |
| pSUL42 | pACYC with gene *AT30* (*XDD1*_*2120*, PCR amplified using sul59+60) inserted under P*_BAD_* control | This study |
| pSUL43 | pACYC with gene *AT31* (*XDD1*_*2027*, PCR amplified using sul61+62) inserted under P*_BAD_* control | This study |
| pSUL44 | pACYC with gene *AT32* (*XDD1*_*2709*, PCR amplified using sul63+64) inserted under P*_BAD_* control | This study |
| pSUL45 | pACYC with gene *AT33* (*XDD1*_*1232*, PCR amplified using sul65+66) inserted under P*_BAD_* control | This study |
| pSUL46 | pACYC with gene *AT34* (*XDD1*_*3898*, PCR amplified using sul67+68) inserted under P*_BAD_* control | This study |
| pSUL47 | pACYC with gene *AT35* (*XDD1*_*3480*, PCR amplified using sul69+70) inserted under P*_BAD_* control | This study |
| pSUL48 | pACYC with gene *AT36* (*XDD1*_*0143*, PCR amplified using sul71+72) inserted under P*_BAD_* control | This study |
| pSUL49 | pACYC with gene *AT37* (*XDD1*_*1214*, PCR amplified using sul73+74) inserted under P*_BAD_* control | This study |
| pSUL50 | pACYC with gene *AT38* (XDD1_*1245*, PCR amplified using sul75+76) inserted under P*_BAD_* control | This study |
| pSUL51 | pACYC with gene *AT39* (*XDD1*_*2923*, PCR amplified using sul77+78) inserted under P*_BAD_* control | This study |
| pEB03 | pACYC with gene *xrdE* inserted under P*_BAD_* control | Ref^2^ |
| pSUL54 (P2R6C7) | Library clone with DNA fragment contains *XDD1*_*1029* -*1030-1031-1032-1033* inserted to pACYC_ara_tacI (*Bgl*II digested) under P*_tacI_* control | This study |
| pSUL55 (P5R7C7) | Library clone with DNA fragment contains *XDD1*_*1030*-*1031*-*1032-1033*-*1034* inserted to pACYC_ara_tacI (*Bgl*II digested) under P*_tacI_* control | This study |
| pSUL56 (P9R4C10) | Library clone with DNA fragment contains *XDD1*_*1234*-*1235-1236* inserted to pACYC_ara_tacI (*Bgl*II digested) under P*_tacI_* control | This study |
| pSUL57 (P2R8C5) | Library clone with DNA fragment contains *XDD1*_*0763*-*0764-0765-0766-0767* inserted to pACYC_ara_tacI (*Bgl*II digested) under P*_tacI_* control | This study |
| pSUL58 | pACYC with gene *staS* (*XDD1*_*1031*, PCR amplified using sul199+200) inserted under P*_BAD_* control | This study |
| pSUL64 | pACYC_SZ17 with the Condensation domain (Cdom) of StaS (PCR amplified using sul199+256) inserted under P*_BAD_* control | This study |
| pSUL65 | pACYC_SZ17 with the first Condensation domain (C1) of XfpS from *X. vietnamensis* DSM 22392 (*Xvie_00354,* PCR amplified using sul259+260) inserted under P*_BAD_* control | This study |
| pSUL71 | pCOLA_SZ18 with the partial A domain (Adom) of StaS (PCR amplified using sul257+258) inserted under P*_BAD_* control | This study |
| pSUL74 | pACYC with the Xviet-XfpS-CAT1 (*Xvie_00354*, PCR amplified using sul279+280) inserted under P*_BAD_* control | This study |
| StaS_pET-28b | pET-28b with gene *staS* inserted under T7 promoter control | This study |
| XfpS-CAT1_pET-28b | pET-28b with gene *xfpS-CAT1* inserted under T7 promoter control | This study |

**Supplementary Table 3.** Primers used in this study.

| **Primer** | **Sequence** | **usage** |
| --- | --- | --- |
| sul251 (ck1007) | CATGGAATTCCTCCTGTTAGCCCAAAAAAAC | Backbone amplification of pACYC, pCOLA and pCDF |
| sul254 (ck1010) | TTAATTAACCTAGGCTGCTGCCACCG |  |
| sul252 | GGGTCTGGATCCAACGAGAAGGAGGAATTAAAATCGAAAAAGG | Backbone amplification of pACYC_SZ17, pair with  sul251 |
| sul253 | TGATCCCGAACCTGAGATAGCTGCAGTC | Backbone amplification of pCOLA_SZ18, pair with  sul254 |
| sul142 | AGATCTGAGCTCTCCCGGGAATTCCAC | Backbone amplification of pEB17 |
| sul143 | CTGCAGGTCGACTCTAGAGGATCGATCC |  |
| sul213 | TCTAGAGGTACCGCATGCGATATCG | Backbone amplification of pCEP_Km |
| sul214 | CATATGCTAGCCTCCTGTTAGCCC |  |
| sul79 | CACACTTTGCTATGCCATAG | Verification of all constructs based on pACYC, pCOLA and pCDF |
| sul80 | CTACCTTAGGACCGTTATAG |  |
| sul124 | GCTATGCCATAGCATTTTTATCCATAAG | Verification of all constructs based on pEB17 |
| sul125 | ACATGTGGAATTGTGAGCGG |  |
| sul146 | GATCGATCCTCTAGAGTCGACCT | Verification of all constructs based on pCEP_Km, pair with sul125 |
| sul01 | GTTTTTTTGGGCTAACAGGAGGAATTCCATGAATACGCTGTTTTCAATTCGGTTAAC | AT1 amplification for cloning into pACYC |
| sul02 | CGGTGGCAGCAGCCTAGGTTAATTAATTATATCGTCAGGCAACGTCTCATTG |  |
| sul03 | GTTTTTTTGGGCTAACAGGAGGAATTCCATGAAAATGCACATTACCGATACTCC | AT2 amplification for cloning into pACYC |
| sul04 | CGGTGGCAGCAGCCTAGGTTAATTAATTAGATATGTTGAATACTTTTCGTGAG |  |
| sul05 | GTTTTTTTGGGCTAACAGGAGGAATTCCATGAAGAATATTTCTCTATTAACTCCCG | AT3 amplification for cloning into pACYC |
| sul06 | CGGTGGCAGCAGCCTAGGTTAATTAACTAAATACACAACGGCAACGCCATAATAATAG |  |
| sul07 | GTTTTTTTGGGCTAACAGGAGGAATTCCATGACAAAATCCCTCCCTACATCTG | AT4 amplification for cloning into pACYC |
| sul08 | CGGTGGCAGCAGCCTAGGTTAATTAATCAAATCAACTTATAAAAATAGGTTG |  |
| sul09 | GTTTTTTTGGGCTAACAGGAGGAATTCCATGGAAATTATAATTAGGGCAACTGAGC | AT5 amplification for cloning into pACYC |
| sul10 | CGGTGGCAGCAGCCTAGGTTAATTAATCATGCAAGCCGGCTATTGATTAAAG |  |
| sul11 | GTTTTTTTGGGCTAACAGGAGGAATTCCATGTACCATCTGAGAGTACCTAAAACAG | AT6 amplification for cloning into pACYC |
| sul12 | CGGTGGCAGCAGCCTAGGTTAATTAATCAGGCTGTAGCCAAACGTTCGGGAGGC |  |
| sul13 | GTTTTTTTGGGCTAACAGGAGGAATTCCATGGAAATTCGGGTATTTCGGCAAGACG | AT7 amplification for cloning into pACYC |
| sul14 | CGGTGGCAGCAGCCTAGGTTAATTAATTAGTCAATAATCAAGCGCTTGCTAAAC |  |
| sul15 | GTTTTTTTGGGCTAACAGGAGGAATTCCATGCAAATACAACCAGCCACCGAAGCGG | AT8 amplification for cloning into pACYC |
| sul16 | CGGTGGCAGCAGCCTAGGTTAATTAATTACTGGATGGGCGAAGATTGGAGGGC |  |
| sul17 | GTTTTTTTGGGCTAACAGGAGGAATTCCATGGCTTTTTTGATCCTTTTTTATGGTAG | AT9 amplification for cloning into pACYC |
| sul18 | CGGTGGCAGCAGCCTAGGTTAATTAATTATTCAGATGTTCCAGGTATTTTCGTTC |  |
| sul19 | GTTTTTTTGGGCTAACAGGAGGAATTCCATGATCAGATCATTTACCGAAGCAGATATG | AT10 amplification for cloning into pACYC |
| sul20 | CGGTGGCAGCAGCCTAGGTTAATTAACTATCTTTTTAAACACATAAAATACTC |  |
| sul21 | GTTTTTTTGGGCTAACAGGAGGAATTCCATGTTAATCAGAGTCGAAATTCCTGTTG | AT11 amplification for cloning into pACYC |
| sul22 | CGGTGGCAGCAGCCTAGGTTAATTAACTATGACCACAGGTGATCAAAATAAGCAG |  |
| sul23 | GTTTTTTTGGGCTAACAGGAGGAATTCCATGTCTATACACGCCAACCTTGAACCTTTG | AT12 amplification for cloning into pACYC |
| sul24 | CGGTGGCAGCAGCCTAGGTTAATTAATTAAATGGAATCATTTCGTCCTCTGTATAAC |  |
| sul25 | GTTTTTTTGGGCTAACAGGAGGAATTCCATGATAATTTCGGAACCAAAACTTCTGGCTG | AT13 amplification for cloning into pACYC |
| sul26 | CGGTGGCAGCAGCCTAGGTTAATTAATCAGCTATCTGTACAGGCGCTTTTTAAATCC |  |
| sul27 | GTTTTTTTGGGCTAACAGGAGGAATTCCATGCTAATAAGAACGGCTTCCACGCAAG | AT14 amplification for cloning into pACYC |
| sul28 | CGGTGGCAGCAGCCTAGGTTAATTAATCACAGCGTAATCGCCATCGTTCTGTTC |  |
| sul29 | TTTTTTGGGCTAACAGGAGGAATTCCATGTTTAGATCAAAATCTTTTAATAATATTAATC | AT15 amplification for cloning into pACYC |
| sul30 | CGGTGGCAGCAGCCTAGGTTAATTAATTAGTTTTTATATCGAAAATGATTAGGATTG |  |
| sul31 | GTTTTTTTGGGCTAACAGGAGGAATTCCATGAAAAAATCCCCTTTATCAGCCGCTG | AT16 amplification for cloning into pACYC |
| sul32 | CGGTGGCAGCAGCCTAGGTTAATTAATTATTGGCGGATAAATGATTGATATTCAAAC |  |
| sul33 | GTTTTTTTGGGCTAACAGGAGGAATTCCATGGGAATAAATGCACCTGAAATATTAACAG | AT17 amplification for cloning into pACYC |
| sul34 | CGGTGGCAGCAGCCTAGGTTAATTAATCAGTTCTTCCTGCCTAATGGCAGCATC |  |
| sul35 | GTTTTTTTGGGCTAACAGGAGGAATTCCATGTCCAGTGGTTCAGAAAATCCATCCG | AT18 amplification for cloning into pACYC |
| sul36 | CGGTGGCAGCAGCCTAGGTTAATTAATTATTGGCAAAATTTATTAGCAAAATAG |  |
| sul37 | GTTTTTTTGGGCTAACAGGAGGAATTCCATGTCTATAGAAAAAAGAGTTAAATG | AT19 amplification for cloning into pACYC |
| sul38 | CGGTGGCAGCAGCCTAGGTTAATTAACTATCTATCTTTTATGTGTCGATAAACC |  |
| sul39 | GTTTTTTTGGGCTAACAGGAGGAATTCCATGACTATCTTCCAATCCGCGATTCAGATCC | AT20 amplification for cloning into pACYC |
| sul40 | CGGTGGCAGCAGCCTAGGTTAATTAATTATGTTATCAGCTCACCCCCCAATTCC |  |
| sul41 | GTTTTTTTGGGCTAACAGGAGGAATTCCATGTTAGAACTCAAACAAGTTACCCCTC | AT21 amplification for cloning into pACYC |
| sul42 | CGGTGGCAGCAGCCTAGGTTAATTAATCATAAAGGAATATTTACATTTTCTATATC |  |
| sul43 | TTTTTTGGGCTAACAGGAGGAATTCCATGAAAAATAACCTAGATATCATTATGTTAATC | AT22 amplification for cloning into pACYC |
| sul44 | CGGTGGCAGCAGCCTAGGTTAATTAATTAATTTAATTCTTTTTTGAAAAAAACG |  |
| sul45 | GTTTTTTTGGGCTAACAGGAGGAATTCCATGACAACAAAAACAACACACGACTACG | AT23 amplification for cloning into pACYC |
| sul46 | CGGTGGCAGCAGCCTAGGTTAATTAATTACAGTTCTTTGACCATCCTGATTTCAC |  |
| sul47 | GTTTTTTTGGGCTAACAGGAGGAATTCCATGAAAACGACCAGAGAGCTATTTCATAATAT | AT24 amplification for cloning into pACYC |
| sul48 | CGGTGGCAGCAGCCTAGGTTAATTAATTAAGGTGATATGTATCTTCTGAGGCGATAAG |  |
| sul49 | GTTTTTTTGGGCTAACAGGAGGAATTCCATGTCCATGATAATTGAAGAAATTACTC | AT25 amplification for cloning into pACYC |
| sul50 | CGGTGGCAGCAGCCTAGGTTAATTAACTATAACTCCCCCACAACTTCCATACAAG |  |
| sul51 | GTTTTTTTGGGCTAACAGGAGGAATTCCATGAAAGAGCGCAGTATCGAATTGGTTGAAG | AT26 amplification for cloning into pACYC |
| sul52 | CGGTGGCAGCAGCCTAGGTTAATTAACTATAAATTCAGCATCAAGATCTTGGAAC |  |
| sul53 | GTTTTTTTGGGCTAACAGGAGGAATTCCATGGAAAAGTCACTAAATGAAATTAGGGTTG | AT27 amplification for cloning into pACYC |
| sul54 | CGGTGGCAGCAGCCTAGGTTAATTAATCAATAAGGTATAGCCGCCAATAAATC |  |
| sul55 | GTTTTTTTGGGCTAACAGGAGGAATTCCATGCCTAATAGAGAAAGAGATAACGCTATCG | AT28 amplification for cloning into pACYC |
| sul56 | CGGTGGCAGCAGCCTAGGTTAATTAACTACAGTATATGACATTGCCAATTTG |  |
| sul57 | GTTTTTTTGGGCTAACAGGAGGAATTCCATGTATAGCTTATGGCACAGCGCATGGG | AT29 amplification for cloning into pACYC |
| sul58 | CGGTGGCAGCAGCCTAGGTTAATTAATTAACGATCAATGGGATGATGAGGATG |  |
| sul59 | GTTTTTTTGGGCTAACAGGAGGAATTCCATGATTTGGCGGTTTATGGTGGATAAAAACC | AT30 amplification for cloning into pACYC |
| sul60 | CGGTGGCAGCAGCCTAGGTTAATTAACTAAATCTTAATAACAGCGATCATCTC |  |
| sul61 | GTTTTTTTGGGCTAACAGGAGGAATTCCATGTTTGGTTATCGTTCTACATCTCCCAAG | AT31 amplification for cloning into pACYC |
| sul62 | CGGTGGCAGCAGCCTAGGTTAATTAATCAAGCATCTTCAATGGCGATTGGTAT |  |
| sul63 | GTTTTTTTGGGCTAACAGGAGGAATTCCATGCCTTTATTTTCAAAAACGATAACTG | AT32 amplification for cloning into pACYC |
| sul64 | CGGTGGCAGCAGCCTAGGTTAATTAATCAATGGGTTGAATCAGGAGAAACCAGC |  |
| sul65 | GTTTTTTTGGGCTAACAGGAGGAATTCCATGAGTCAGCGTGGACTTGAAGCATTATTG | AT33 amplification for cloning into pACYC |
| sul66 | CGGTGGCAGCAGCCTAGGTTAATTAACTAGAGCGTCAACATCAGATTAACAATG |  |
| sul67 | GTTTTTTTGGGCTAACAGGAGGAATTCCATGTTAACGATCCGGGATGCAACGATAG | AT34 amplification for cloning into pACYC |
| sul68 | CGGTGGCAGCAGCCTAGGTTAATTAACTACTCGGATAAATTTTTAGCCATGATATG |  |
| sul69 | GTTTTTTTGGGCTAACAGGAGGAATTCCATGGAAAAAATAACAGATTGGGTTACATTG | AT35 amplification for cloning into pACYC |
| sul70 | CGGTGGCAGCAGCCTAGGTTAATTAATCAACGTGGCTTCGTCGTTAACCATATAAG |  |
| sul71 | GTTTTTTTGGGCTAACAGGAGGAATTCCATGAAAATAAAAACACTCCTTTCAGGATG | AT36 amplification for cloning into pACYC |
| sul72 | CGGTGGCAGCAGCCTAGGTTAATTAATTAAAAAAGCACTTTGCCTAATGGC |  |
| sul73 | GTTTTTTTGGGCTAACAGGAGGAATTCCATGAATAAATCTGTTAATTTTACTCATGATG | AT37 amplification for cloning into pACYC |
| sul74 | CGGTGGCAGCAGCCTAGGTTAATTAATTATCTGTTTTGTTGTAGTAACCATGTTAG |  |
| sul75 | GTTTTTTTGGGCTAACAGGAGGAATTCCATGTCTACTGTTGCTACTGTATTATTTG | AT38 amplification for cloning into pACYC |
| sul76 | CGGTGGCAGCAGCCTAGGTTAATTAATCATTTTGAAACGCTATTTAGGTTCTTAG |  |
| sul77 | GTTTTTTTGGGCTAACAGGAGGAATTCCATGATATCTAAAAATACATTAAATATAG | AT39 amplification for cloning into pACYC |
| sul78 | CGGTGGCAGCAGCCTAGGTTAATTAATCACACAAAATCGGAACGCCGCAGCCGATAC |  |
| sul136 | CGATCCTCTAGAGTCGACCTGCAGATTCTTTTGCAAAAAGTTAGTAC | Amplification of HAL and HAR for *AT38* gene deletion |
| sul137 | CCGGGAGAGCTCAGATCTGATTATTGTCCTTGCTATTACTGCG |  |
| sul138 | GTTGGGGACATTTTCCTTGCGTCCGCTATGAAAATTTATTTAGAAAGATTATG |  |
| sul139 | CATAATCTTTCTAAATAAATTTTCATAGCGGACGCAAGGAAAATGTCCCCAAC |  |
| sul140 | GTCTAATCTGCGTTTTAATACCCGTG | Verification of *AT38* gene deletion |
| sul141 | GTTTTTGCTGGTAATTGGCGTAGGC |  |
| sul155 | CACCAATCGACAATATTTCGGTGGGG | Verification of *xrdE* gene deletion |
| sul156 | CCGTGATCCTGACCGAAGAGCTGATC |  |
| sul201 | GGATCGATCCTCTAGAGTCGACCTGCAGGCTGGCAACGCTCAAGTAAGATTGGC | Amplification of HAL and HAR for *staS* gene deletion |
| sul202 | CATCATAGAGGAGTATTCACATGTAAGCGGGAAATCGGTGAAATTATG |  |
| sul203 | CATAATTTCACCGATTTCCCGCTTACATGTGAATACTCCTCTATGATG |  |
| sul204 | GTGGAATTCCCGGGAGAGCTCAGATCTGCGGGATTATTAGGCGGTCATGC |  |
| sul205 | CTTCCCAGCGTTTTTTTCCGGCTCCGTG | Verification of *staS* gene deletion |
| sul206 | CTTTGAATCATCCCCACCGGGATATC |  |
| sul161 | CACTCTTTGCTGCCATCTAT | Verification of DC gene deletion |
| sul164 | TGAGCAACCGTAATATGAGG |  |
| sul199 | GTTTTTTTGGGCTAACAGGAGGAATTCCATGGATAAAATTCTGGCTTCTCCATTTATTG | StaS amplification for cloning into pACYC |
| sul200 | CGGTGGCAGCAGCCTAGGTTAATTAATTAACTCGCCTCTTTCAGTGAATCATTG |  |
| sul256 | CTCGTTGGATCCAGACCCTCCGTATTCCAGCAATGTCTGTGTC | StaS_Cdom amplification for cloning into pACYC_SZ17, pair with sul199 |
| sul257 | CTATCTCAGGTTCGGGATCAAACTCACAACGACATAACAGTCTGG | StaS_Adom amplification for cloning into pCOLA_SZ18 |
| sul258 | CAGCCTAGGTTAATTAATTAACTCGCCTCTTTCAGTGAATCATTG |  |
| sul259 | GGCTAACAGGAGGAATTCCATGGATAACATTTTGGCCTCTCCATTTAC | Xviet-XfpS-C1 domain amplification for cloning into pACYC_SZ17 |
| sul260 | CTCGTTGGATCCAGACCCCCCGAATTCCAATAGTGCCCGGC |  |
| sul279 | GTTTTTTTGGGCTAACAGGAGGAATTCCATGGATAACATTTTGGCCTCTCC | Xviet-XfpS-CAT1 amplification for cloning into pACYC |
| sul280 | CGGTGGCAGCAGCCTAGGTTAATTAATCATGTTTCTGTCAACTGTGTTGCC |  |
| MD05_fw | CTTAGCGGAATATTGCCTAT | Verification of the *X. griffiniae* Δ*DC* mutant. |
| MD06_rv | CTTCATCAACAATCTGGTTC |  |
| MD32_fw | TACTCCCGCAGAGGATTTGGATA | Verification of the *X. griffiniae* Δ*staS* mutant. |
| MD33_rv | CACGATCACGCCGTTGTTTTAAC |  |
| StaS_NcoI_fw | CGACTCCATGGATAAAATTCTGGCTTCTC | Subcloning StaS into pET-28b vector |
| StaS_XhoI_rv | CTCACTCGAGACTCGCCTCTTTCAGTGAATC |  |
| XfpS-CAT1_28_FF | CTTTAAGAAGGAGATATACCATGGATAACATTCTGGCC | Subcloning XfpS into pET-28b vector |
| XfpS-CAT1_28_FR | TGGTGCTCGAGTGCGGCCGCAGTGGTAGCCAGTTG |  |
| XfpS-CAT1_28_VF | CGGTACAACTGGCTACCACTGCGGCCGCACTC |  |
| XfpS-CAT1_28_VR | GAGGCCAGAATGTTATCCATGGTATATCTCCTTCTTAAAG |  |

**Supplementary Table 4.** Cryo-EM data collection and refinement statistics.

|  | StaS (PDB 31EM, EMD-58340) |
| --- | --- |
| **Data collection and processing** |  |
| Microscope | Titan Krios III |
| Magnification | 215,000x |
| Voltage (kV) | 300 |
| Electron exposure (e^-^/Å^2^) | 40 |
| Defocus range (μM) | -0.5 to -1.75 |
| Pixel size (Å) | 0.576 |
| Micrographs | 36,909 |
| Initial particles | 3,200,729 |
| Final particles | 83,704 |
| Symmetry | C2 |
| Map resolution (Å) | 4.01 |
| Box size (pixels) | 360 |
| FSC threshold | 0.143 |
| **Refinement** |  |
| MolProbity score | 2.05 |
| Clash score | 13.71 |
| Rotamer outliers | 0 |
| RMSD |  |
| Bond lengths (Å) | 0.002 |
| Bond angles (°) | 0.571 |
| Ramachandran plot |  |
| Favored (%) | 93.99 |
| Allowed (%) | 5.68 |
| Outliers (%) | 0.33 |
| Model composition |  |
| Chains | 2 |
| Non-hydrogen atoms | 7482 |
| Protein residues | 910 |

**Supplementary References**

1.	Proschak, A., Schultz, K., Herrmann, J., Dowling, A.J., Brachmann, A.O., ffrench-Constant, R., Muller, R., and Bode, H.B. (2011). Cytotoxic fatty acid amides from Xenorhabdus. Chembiochem *12*, 2011-2015. 10.1002/cbic.201100223.

2.	Bode, E., He, Y., Vo, T.D., Schultz, R., Kaiser, M., and Bode, H.B. (2017). Biosynthesis and function of simple amides in Xenorhabdus doucetiae. Environ Microbiol *19*, 4564-4575. 10.1111/1462-2920.13919.

3.	Hirschmann, M., Grundmann, F., and Bode, H.B. (2017). Identification and occurrence of the hydroxamate siderophores aerobactin, putrebactin, avaroferrin and ochrobactin C as virulence factors from entomopathogenic bacteria. Environ Microbiol *19*, 4080-4090. 10.1111/1462-2920.13845.

4.	Kegler, C., and Bode, H.B. (2020). Artificial Splitting of a Non-Ribosomal Peptide Synthetase by Inserting Natural Docking Domains. Angew Chem Int Ed Engl *59*, 13463-13467. 10.1002/anie.201915989.

5.	Bode, E. (2017). Structure, function and biosynthesis of natural products from *Xenorhabdus doucetiae.* Doctorate (Johann Wolfgang Goethe-Universität in Frankfurt am Main).

6.	Bode, E., Heinrich, A.K., Hirschmann, M., Abebew, D., Shi, Y.N., Vo, T.D., Wesche, F., Shi, Y.M., Grun, P., Simonyi, S., et al. (2019). Promoter Activation in Deltahfq Mutants as an Efficient Tool for Specialized Metabolite Production Enabling Direct Bioactivity Testing. Angew Chem Int Ed Engl *58*, 18957-18963. 10.1002/anie.201910563.

7.	Bode, E., Brachmann, A.O., Kegler, C., Simsek, R., Dauth, C., Zhou, Q., Kaiser, M., Klemmt, P., and Bode, H.B. (2015). Simple "on-demand" production of bioactive natural products. Chembiochem *16*, 1115-1119. 10.1002/cbic.201500094.

8.	Shi, Y.M., Hirschmann, M., Shi, Y.N., Ahmed, S., Abebew, D., Tobias, N.J., Grun, P., Crames, J.J., Poschel, L., Kuttenlochner, W., et al. (2022). Global analysis of biosynthetic gene clusters reveals conserved and unique natural products in entomopathogenic nematode-symbiotic bacteria. Nat Chem *14*, 701-712. 10.1038/s41557-022-00923-2.

9.	Tobias, N.J., Wolff, H., Djahanschiri, B., Grundmann, F., Kronenwerth, M., Shi, Y.M., Simonyi, S., Grun, P., Shapiro-Ilan, D., Pidot, S.J., et al. (2017). Natural product diversity associated with the nematode symbionts Photorhabdus and Xenorhabdus. Nat Microbiol *2*, 1676-1685. 10.1038/s41564-017-0039-9.

10.	Schimming, O., Fleischhacker, F., Nollmann, F.I., and Bode, H.B. (2014). Yeast homologous recombination cloning leading to the novel peptides ambactin and xenolindicin. Chembiochem *15*, 1290-1294. 10.1002/cbic.201402065.

11.	Thoma, S., and Schobert, M. (2009). An improved Escherichia coli donor strain for diparental mating. FEMS Microbiol Lett *294*, 127-132. 10.1111/j.1574-6968.2009.01556.x.

12.	Lorenzen, W., Ahrendt, T., Bozhuyuk, K.A., and Bode, H.B. (2014). A multifunctional enzyme is involved in bacterial ether lipid biosynthesis. Nat Chem Biol *10*, 425-427. 10.1038/nchembio.1526.

13.	Abbood, N., Effert, J., Bozhueyuek, K.A.J., and Bode, H.B. (2023). Guidelines for Optimizing Type S Nonribosomal Peptide Synthetases. ACS Synth Biol *12*, 2432-2443. 10.1021/acssynbio.3c00295.

14.	Bode, E., Heinrich, A.K., Hirschmann, M., Abebew, D., Shi, Y.N., Vo, T.D., Wesche, F., Shi, Y.M., Grün, P., Simonyi, S., et al. (2019). Promoter Activation in Δhfq Mutants as an Efficient Tool for Specialized Metabolite Production Enabling Direct Bioactivity Testing. Angew Chem Int Ed Engl *58*, 18957-18963. 10.1002/anie.201910563.
