## Supplementary Methods for "Acyl amides derived from a bacterial symbiont control development of its nematode host"

**Materials and methods**

**General methods**

All reagents and chemicals were obtained from Sigma-Aldrich, ROTH, VWR, and BD. Genomic DNA was isolated using the Monarch^®^ Genomic DNA Purification Kit. DNA purification was done from 1% (w/v) TAE agarose gels using Monarch^®^ DNA Gel Extraction Kit. Plasmids were isolated from *E. coli* by using Monarch^®^ Plasmid Miniprep Kit. DNA primers were purchased from Sigma-Aldrich. DNA polymerases (Taq and Q5) were purchased from New England Biolabs. Polymerases were used according to the manufacturer’s instructions. Plasmids used for gene expression and deletion were assembled using the NEB Gibson assembly^®^ Kit.

HPLC-UV-MS analysis was conducted on an Agilent Infinity II 1290 system coupled to an Bruker amaZon speed mass spectrometer (ESI-MS spectra were recorded in positive-ion-mode with the mass range from 100-1200 *m*/*z,* ultraviolet at 190–640 nm) with an ACQUITY UPLC BEH C18 column (130 Å, 2.1 mm × 50 mm, Waters) at a flow of 0.4 mL/min (5–95% acetonitrile/water with 0.1% formic acid, v/v, 14 min). HPLC-HR-MS analysis was conducted on an Agilent Infinity III 1290 system coupled to a Bruker timsTOF flex mass spectrometer (VIP-HESI spectra were recorded in positive-ion-mode with the mass range from 100-2000 *m*/*z*) with an ACQUITY UPLC BEH C18 column (130 Å, 2.1 mm × 50 mm, Waters) at a flow of 0.4 mL/min (5–95% acetonitrile/water with 0.1% formic acid, v/v, 14 min). Data processing and analysis were conducted using Bruker Compass DataAnalysis version 5.3.

**Bacteria cultivation**

Unless otherwise specified, all strains were cultured in LB medium (10 g/L tryptone, 5 g/L yeast extract, and 5 g/L NaCl) or on an LB agar plate (1.5 % (w/v) agar was added) at 37 ºC (for *E. coli* strains) or 28 ºC (for *Xenorhabdus* strains). The antibiotics kanamycin (50 µg/mL), chloramphenicol (34 µg/mL) and spectinomycin (50 µg/mL) were added when appropriate. For the production culture of *X. doucetiae* and its gene deletion and promoter exchange mutants, the overnight LB culture was transferred into 5 mL LB/XPP3 medium^1^ with a starting OD_600_ of 0.1. It was incubated at 28 ºC with shaking at 200 revolutions per minute (rpm) for 3 days. For induction of promoter exchange strains or for induction of heterologous expression in *E. coli*, 0.2% (w/v) L-arabinose was added to the culture at the start of cultivation, while, IPTG (100 µM) was added for induction of library clones in *E. coli*.

**Galleria mellonella rearing**

*Galleria mellonella* (Lepidoptera: *Pyralidae*) were reared in tall, wide-neck glass jars with ultra-thin stainless-steel wire mesh at the top, containing a filter paper on the top (sieve size of 0.1 mm^2^). The jars were closed by a perforated hose clamp. The media or insect diet was composed of 22% glycerol, 22% organic honey, 4% water, 48% cereal, and 4% instant dry baker’s yeast. The mixture was incubated overnight at 30°C and then stored at 4°C for further use. The larvae were placed in jars containing the culture media and incubated at 28°C. Three days after the larvae molted into adults, the filter paper containing eggs was removed and cut into small pieces, which were placed in a new jar incubated under the same conditions^2^.

**Entomopathogenic nematodes (EPNs) rearing**

A Whatman filter paper of 10 cm was placed on the lid of a plastic Petri dish (100 mm diameter), where 1 mL of the infective juvenile (IJ) suspension (1500 IJs/mL) was distributed. Six last instar larvae of the insect *Galleria mellonella*, previously sprayed with ethanol 70% followed by distilled water, were placed in the Petri dish, which was covered with the bottom part. The larvae were incubated at 24°C. After three days, the insect cadavers exhibiting signs of entomopathogenic nematode infection were removed and placed in a modified White Trap.^3^ A White Trap is a technique employed to obtain the next generation of IJs from infected insects in a clean manner, without debris from the corpse. This method involves placing the lid of a Petri dish (60 mm) inside the bottom of a larger one (100 mm). A Whatman filter paper measuring 5 cm was placed above the lid of the smaller Petri dish, into which the removed insect cadavers were placed. The larger Petri dish was filled with Ringer’s solution ([NaCl](https://en.wikipedia.org/wiki/Sodium_chloride) 9 g/L, [KCl](https://en.wikipedia.org/wiki/Potassium_chloride" \o "Potassium chloride) 0.4 g/L, [CaCl_2_](https://en.wikipedia.org/wiki/Calcium_chloride) 0.4 g/L, and [NaH_2_CO_3_](https://en.wikipedia.org/wiki/Sodium_bicarbonate) 0.2 g/L in distilled water), ensuring that the cadaver was not in direct contact with it. The larger Petri dish was covered with the corresponding lid. The White Traps were incubated at 24°C, and after 1–2 weeks, the IJs contained in the ringer solution were harvested. The nematodes were rinsed three times through a filter with a vacuum pump and subsequently placed into a tissue culture flask (175 mL) containing 75 mL of Ringer’s solution. The tissue culture flasks were stored at 18°C for subsequent use^4^.

**Axenic and Aposymbiotic IJs isolation**

Ten *G. mellonella* larvae previously sprayed with ethanol (70%) followed by distilled water were placed on a 100 × 15 mm Petri dish containing a Whatman filter paper. A 1 mL suspension of infective juveniles (120 IJs/mL) was distributed, and the Petri dish was incubated at 24°C. Four days post-infection, which is the period during which the first generation of adults should be fully developed according to the nematode development cycle, the infected larvae were dissected in a Petri dish containing 20 mL of M9 buffer (KH_2_PO_4_ 15 g/L, Na_2_HPO_4_ 30 g/L, NaCl 25 g/L, and MgSO_4_ (1 M) 5 mL/L in distilled water). Gravid females were collected (20-30) and transferred to a watch glass containing M9 buffer, where they were washed three times in order to remove debris from the insect cadaver. An axenic solution (6.75 mL distilled water, 1.25 mL 5M NaOH, and 2 mL of 3% (v/v) chlorine) was prepared and the M9 buffer was removed by pipetting and replaced with the axenic solution in a watch glass. The solution was transferred to a 15 mL Eppendorf tube, which was slightly agitated for 10 minutes and vortexed for 10 seconds every 2 minutes during that period. At the same time, the female cuticular disintegration was constantly checked. After 10 minutes, the tube was centrifuged at 7,000 rpm for two minutes. The supernatant was removed, and the egg pellet was rinsed three times with sterile Ringer's solution.

At the final washing step, the egg pellets were resuspended in 300 μL of sterilized Ringer’s solution by gentle pipetting and then transferred to a sterilized petri dish for examination. The egg solution (100 μL) was plated on liver kidney agar (ground beef kidney 100g/L, ground beef liver 100g/L, NaCl 5g/L and agar 15g/L in distilled water), thereby enabling EPNs to grow without a bacterial symbiont. Subsequently, the plates were incubated at 24°C for 24 hours. Following this period, the plate was placed in a White Trap and continuously monitored for egg hatching and developmental stages. Approximately two to three weeks after inoculation, axenic infective juveniles emerged. These juveniles were then used to infect *G. mellonella* larvae, as previously described, in order to obtain aposymbiotic IJs^3^.

**Nematode development evaluation**

To analyze the effect of NPs on the development of nematodes, the recovery process of IJ of *Steinernema diaprepesi* was evaluated in the presence of the WT strain of *X. doucetiae* as control, deletion or promoter exchange mutants for the synthesis of natural products (NPs) such as xenoamicin (XAB), protegomycin (PRT), xenorhabdin (XRD), xenocoumacin (XCN), GameXPeptide (GXP), and amides (phenylethylamide and tryptamide). Bacterial strains were cultivated overnight in LB medium. For nematode recovery analysis, a volume of 100 μL of the bacterial culture (OD600 5.0) was plated in a 6-well plate containing 5 mL of NGM (Nematode Growth Medium) agar (NaCl 3 g/L, tryptone 2.4 g/L, yeast extract 0.8 g/L, 1 mM CaCl_2_ 1 mL/L, 1 mM MgSO_4_ 1 mL/L and Agar 20 g/L in distilled water) and plates were incubated for 72 h at 30°C. After the incubation period, approximately 50 aposymbiotic IJs surface-sterilized with a solution of hyamine (0.4% (w/v)) were added to the plates containing bacterial strains to be tested. After four days, the number of recovered nematodes in the 6-well plates was counted under a dissecting microscope at 8 x magnifications. When it was required, pure compounds were added to the media. PEA and stripteamide were used to supplement the media to a final concentration of 1 mM.

**Hemolymph samples from EPN infected Galleria mellonella**

*Galleria mellonella* instar larvae were surface-sterilized with 70% (v/v) ethanol and individually placed in 1.5 mL microcentrifuge tubes containing filter paper. Each larva was inoculated with 50 infective juveniles (IJs) of *Steinernema diaprepesi*. The larvae were incubated at 24 °C throughout the experimental period. The entire experiment was conducted in triplicate using independent biological replicates.

At 12-hour intervals post-infection, three larvae were randomly selected until 120 hours post-infection. From each larva, 30 µL of hemolymph was collected and stored at −20 °C until analysis. At each time point, larvae were dissected to monitor nematode development by assessing the presence of distinct life cycle stages. Following completion of the sampling period, hemolymph samples were diluted 1:1 (v/v) with methanol (MeOH) and centrifuged at 13,000 rpm for 20 minutes. Subsequently, 50 µL of the resulting supernatant was transferred to analysis vials. Metabolomic analysis was performed using a timsTOF mass spectrometer.

**Construction and screening of AT expression strains**

To identify additional acyltransferases (ATs) potentially involved in stripteamide biosynthesis, we performed a systematic hidden Markov model (HMM) search against the *X. doucetiae* FRM16 genome. HMM profiles for 50 members of the acetyltransferase clan (CL0257) were downloaded from the InterPro database (https://www.ebi.ac.uk/interpro/set/pfam) and used as queries for HMMER searches (https://www.ebi.ac.uk /Tools/hmmer/search/hmmsearch). After merging all matches and removing redundancies, we obtained 40 AT candidate genes (including the previously identified XrdE) for subsequent expression and screening. Expression constructs for each candidate were generated via Gibson assembly. Candidate genes were amplified from wild-type *X. doucetiae* FRM16 genomic DNA using primers with overhangs homologous to the pACYC vector backbone (primer sequences listed in Supplementary Table 3). The vector backbone was amplified with primers sul251 (ck1007) and sul254 (ck1010). Purified PCR products were assembled with the vector backbone using Gibson assembly, and the resulting plasmids (pSUL13–pSUL51; Supplementary Table 2) were transformed into *E. coli* DH10B. Correct clones were verified by colony PCR and sequencing prior to screening.

The 40 AT expression strains were cultivated in LB medium supplemented with 34 μg/mL chloramphenicol at 25°C. Four hours post-inoculation, expression was induced with 0.2% L-arabinose (ARA), and 1 mM phenylethylamine (PEA) was added as a substrate. After shaking at 200 rpm for 2 days, 20 μL of culture was mixed with 180 μL methanol and vortexed thoroughly for 1 min. The mixture was centrifuged at 13,000 ×*g* for 20 min, and 100 μL of supernatant was subjected to LC-MS analysis. Stripteamide production was monitored by the presence of characteristic fragment ions at *m/z* 105.06 and 122.09 ([M+H]⁺), as previously reported^5^.

**Construction and screening of genomic expression library**

Genomic DNA of *X. doucetiae* was digested with *Sau3*AI and fractionated on a 1% agarose gel. Fragments ranging from 3–6 kb were excised, purified, and ligated into *Bgl*II-digested and dephosphorylated pACYC_ara_tacI vector using T4 DNA ligase. The ligation products were transformed into *E. coli* DH10B, and individual colonies were picked using a PIXL colony picker (Singer Instruments) into 36 × 96-well microplates to generate the genomic library.

For library screening, each 96-well plate was pooled as a single sample and cultured in LB medium containing chloramphenicol. After 4 h of incubation at 37°C, expression was induced with 100 µM IPTG, and 1 mM PEA was added as substrate. Cultures were shaken at 25°C, 200 rpm for 2 days, then processed and analyzed by LC-MS as described for the AT expression strains to identify plates producing stripteamides. Positive plates were sequentially combined in rows and columns followed by LC-MS analysis, ultimately identifying four individual clones: P2R6C7, P2R8C5, P5R7C7, and P9R4C10, capable of stripteamide production.

**Construction of gene deletion mutants**

Gene deletion mutants, including ∆*AT38*, ∆*xrdE* ∆*AT38,* ∆*staS,* ∆*xrdE* ∆*staS,* ∆*AT38* ∆*staS,* ∆*triple* (∆*xrdE* ∆*AT38* ∆*staS*), ∆*tertra* (∆*xrdE* ∆*AT38* ∆*staS* ∆*DC*), ∆*DC_griff_*, ∆*staS_griff_*, and ∆*DC_griff_* ∆*staS_griff_* were constructed following the protocol described below for ∆*AT38*. For deletion of *AT38*, left and right homology arms (HAL and HAR) flanking the target gene were amplified using primer pairs sul136/sul137 and sul138/sul139, respectively. The pEB17 vector backbone was amplified with primers sul142/sul143. The three fragments were assembled via Gibson assembly to generate plasmid pEB_de-AT38. This plasmid was electroporated into *E. coli* ST18 and subsequently transferred into *X. doucetiae* FRM16 by intergeneric conjugation. Double-crossover recombinants were screened and verified by colony PCR using primers sul140/sul141. The pEB_de-AT38 plasmid was then erased by counter-selection on medium containing 6% (w/v) sucrose. Final deletion mutants were confirmed by PCR and sequencing.

**Cloning of StaS and XfpS into expression vectors**

The *staS* gene was amplified by PCR from the pSUL58 construct using primers StaS_NcoI_fw/StaS_XhoI_rv. The amplified gene as well as pET-28b vector was then digested with NcoI and XhoI restriction enzymes and then gel extracted. Ligation of the digested products was performed using T4 DNA ligase and transformed into *E. coli* DH5α cells. Single colonies were inoculated into LB media with 50 µg/mL of kanamycin at 37 °C overnight. Plasmid purification was performed and sequences were verified by Sanger sequencing.

The *xfpS-CAT1* gene was amplified by PCR from the pSUL74 construct using primers XfpS_28_FF/XfpS_28_FR. The pET-28b vector was amplified using primers XfpS_28_VF/XfpS_28_VR. The fragments were gel extracted and ligated using Gibson assembly. The products were transformed into *E. coli* DH5α cells. Single colonies were inoculated into LB media with 50 µg/mL of kanamycin at 37 °C overnight. Plasmid purification was performed and sequences were verified by Sanger sequencing.

**Expression and purification of StaS and XfpS**

StaS-pET-28b and XfpS-CAT1-pET-28b plasmids were transformed into chemically competent *E. coli* BL21(DE3) cells. Single colonies were used to inoculate overnight cultures containing LB media containing 50 µg/mL of kanamycin and incubated at 37 °C overnight. 1 L cultures of LB media were inoculated using the overnight cultures and grown at 37 °C until OD600 0.6-0.8, after which cells were cooled to 16 °C and induced with 0.3 mM IPTG for overnight expression. Cells were harvested by centrifugation for 10 minutes at 5000 rpm at 4 °C. For StaS, cell pellets were resuspended in lysis buffer (20 mM Tris pH 8.0, 500 mM NaCl, 20 mM imidazole) supplemented with 1x homemade protease inhibitor cocktail (100 µM AEBSF, 1 µM E-64, 1 µM leupeptin, 0.15 µM aprotinin) and lysed using a cell disruptor (4°C, Constant Systems). Cell debris was removed by centrifugation at 40,000×*g* for 30 min at 4 °C. The supernatant was used for purification on a gravity column containing Ni-NTA beads equilibrated with lysis buffer. The beads were washed with lysis buffer and the protein was eluted with 10 mL elution buffer (20 mM Tris pH 8.0, 500 mM NaCl, 250 mM imidazole). The protein was desalted using a desalting column (HiPrep 26/10, Cytiva) in SEC buffer (20 mM Tris pH 8.0, 150 mM NaCl). Desalted proteins were then purified using a HiTrapQ column at a flow rate of 2.5 mL min-1 using a gradient from A to B over 20 CV (Buffer A = 20 mM Tris pH 8.0, Buffer B = 20 mM Tris pH 8.0, 1 M NaCl). Protein fractions were pooled, concentrated, and further purified by size-exclusion chromatography using a Superdex 200 10/300 column (Cytiva) pre-equilibrated in SEC buffer at a flow rate of 1 mL/min. The monomer and dimer fractions were collected separately and run again by size-exclusion chromatography. Proteins fractions were pooled and concentrated and aliquots were flash frozen in liquid nitrogen and stored at -80 °C until further use. Protein concentration was measured using a Nanodrop One (Thermo Fisher Scientific) and purity was assessed by SDS-PAGE. XfpS-CAT1 was purified following the same protocol but with a different lysis buffer (20 mM Tris pH 8.0, 300 mM NaCl, 10% glycerol, 20 mM imidazole, 2 mM BME) and elution buffer (20 mM Tris pH 8.0, 300 mM NaCl, 10% glycerol, 250 mM imidazole, 2 mM BME).

**Cryo-EM grid preparation and data collection**

Purified StaS was concentrated to 0.31 mg/mL. Quantifoil 0.6/1 200 mesh Au grids were washed with chloroform, glow discharged for 90 s at 25 mA, and treated with 43 µM DDM and blot dried immediately prior to use. 3 µL StaS were added to the grid and plunge-frozen in liquid ethane using a FEI Vitrobot Mark IV with 2.5 s blot time at 95% humidity at 22 °C. Grids were screened on a 200 kV Glacios cryo-TEM, and high-resolution data collection was performed on a 300 kV Titan Krios III cryo-TEM equipped with a Falcon 4i detector. 36,909 movies were collected with a pixel size of 0.576 Å, total electron dose of 40 e^-^/Å^2^, and defocus range of -0.5 to -1.75 µm.

Purified XfpS-CAT1 was concentrated to 0.32 mg/mL. Quantifoil 0.6/1 200 mesh Au grids were washed with chloroform and glow discharged for 30 s at 20 mA. 3 µL XfpS-CAT1 were added to the grid with 1.5 s blot time at 95% humidity at 22 °C. Grids were screened and high-resolution data collection was performed as mentioned above for StaS, with a total of 31,899 movies collected.

**Map reconstruction and model building**

Data were processed using crYOLO v 1.9.9^6^and cryoSPARC v 4.7.1^7^. Processing workflows are shown in Supplementary Figures 21 and 22. Both datasets were processed using motion corrected micrographs from Diamond Light Source (MotionCor2) for Patch CTF Estimation. For StaS, 3,200,729 particles were picked using crYOLO and then extracted 2x binned. Several rounds of 2D classification were performed to remove junk particles. Iterative ab initio, heterogeneous refinement, and non-uniform (NU) refinement runs were performed, the best set of particles were re-extracted unbinned, and the final model was obtained after a final NU refinement run. For XfpS-CAT1, 1,433,989 particles were picked using crYOLO and then extracted 2x binned. Several rounds of 2D classification were performed, and selected classes were split into two groups based on their orientations. The best set of particles were re-extracted unbinned for each group and one round of ab initio and several rounds of heterogeneous refinement and NU refinement were run. The final models for XfpS-CAT1 were obtained after a final NU refinement run. For both datasets, resolution was estimated based on gold-standard Fourier shell correlation (GSFSC) at the 0.143 criterion. A local resolution map of StaS was generated in cryoSPARC, and structure figures were generated using ChimeraX^8^.

Attempts to dock an AlphaFold3 (AF3) model^9^ of the whole StaS protein into the cryo-EM map failed due to a rotation of the N-terminal portion of StaS between the map and model. We therefore split the AF3 model at residue 205 and used VESPER(S2M) to dock each part of the StaS protein individually. The placed fragments (residues 1-205 and 206-520) were then connected in Coot, followed by iterative cycles of real-space refinement in Phenix^10^. Dock and rebuild in Phenix was used on the resulting model followed by manual building in Coot^11^ and real-space refinement in Phenix. Final structure statistics are shown in Supplementary Table 4.

**SYNZIP mediated splitting and splicing of StaS**

The splitting site for StaS was selected based on the conserved WNATE motif located in the C–A interdomain linker of NRPS modules^12^. Sequence alignment with several NRPS CAT and C/EAT modules identified the corresponding motif in StaS as GNSQR. StaS was split between the glycine (G) and asparagine (N) residues of this motif, yielding two fragments: StaS_C_dom_ (N-terminal to the cleavage site) and StaS_A_part_ (C-terminal to the cleavage site). These fragments were PCR-amplified using primer pairs sul199/sul256 and sul257/sul258, respectively, and ligated into the backbones of pACYC_SZ17 and pCOLA_SZ18, generating pSUL64 and pSUL71.

**Chemical synthesis C12 acyl-SNAC**

250mg *N*-Acetylcysteamine (SNAC) (119.19g/mol, 2mmol, 1.00eq) and 376mg of lauric acid (200,32 g/mol, 1,88mmol, 0,95eq) were weighed into a 50mL round-bottom-flask, dissolved in 10mL of DCM and cooled to 0°C in an ice bath. 50mg of DMAP (122,08g/mol, 0,43mmol, 0,21eq) and 399mg of EDCI (155,25g/mol, 2,2mmol, 1,10eq) were added to the cold, stirring mixture. The reaction was allowed to warm up to room temperature and quenched after stirring at room temperature for 2 days by adding 20mL of 2N HCL. The emulsion was extracted 3 times with 10mL DCM. The combined organic phases were subsequently washed with a saturated solution of NaHCO_3_ and NaCl. The crude organic phases were combined and dried with NaSO_4_ . Lastly, the solvent was removed under reduced pressure to afford a white, crystalline substance which weighed out to be 451mg (301.49g/mol, 1.4mmol, 79.5%). The identity was further confirmed by LC-MS with a gradient elution from 5-95% Acetonitrile/water with 0.1% of formic acid over 4min and NMR spectroscopy.

**C12 acyl-SNAC:** (ESI)-MS [M+H]^+^ calculated for C_16_H_32_NO_2_S: 302.21 m/z; found: 302.10m/z with t_R_ =3.4 min. ^1^H- NMR (CDCl_3_, 500 MHz): δ 5.83 (bs, 1H), 3.43 (q, J = 6,03Hz, 2H), 3.02 (t, J = 6.41 Hz, 2H), 2.57 (t, J = 7,39 Hz, 2H), 1.97 (s, 3H), 1.64 (m, 2H), 1.30 (m, 16H), 0.88 (t, J = 6.97 Hz, 3H). ^13^C NMR (126 MHz, CDCl_3_): δ 200.48, 170.42, 44.29, 39.96, 32.04, 29.74-29.46, 29.40, 29.37, 29.24, 29.08, 28.58, 25.82, 23.36, 22.82, 14.25.

**In vitro assays of StaS**

Enzyme assays were carried out in a 50 μL reaction containing 20 mM Tris-HCl (pH 8.0), 5 mM ATP (use only when C12 fatty acid as substrate), 10 mM MgCl_2_, 2 mM amine substrate (PEA) and 2 mM acyl donor (C12 fatty acid or C12 acyl-SNAC). Reactions were initiated by addition of purified StaS (monomeric or dimeric form), incubated at 30°C for 30 min, and quenched by 50 μL cold MeOH prior to analysis. Amide formation was monitored by HPLC-MS using an amaZon speed mass spectrometer.

**Phylogenetic analysis of StaS**

352 C-domains containing ^L^C_L_, ^D^C_L_, dual C/E domains and starter C-domains were extracted from various NRPS. This dataset (consisting of amino acids) was aligned using mafft^13^ and gaps were manually trimmed. IQ-TREE^14^ multicore version 2.0.7 was used for maximum likelihood tree calculation. JTT^15^ was used as the best fit model with a 7-category free rate distribution of among site variation.

**Species tree inference**

GToTree^16^ v1.7.0050 was used to construct the species tree. Single-copy gene (SCG) set for proteobacteria targeting 119 target genes were used. Genomes were provided as fasta files or gene bank files and prodigal^17^ was used to predict the SCGs. The target genes were then extracted using HMMER3^18^ v3.3.252. The alignment was subsequently aligned and trimmed using muscle v5^19^ and trimAI v1.4.rev^20^. FastTree2 v2.1^21^ concatenated all SCGs and inferred the species tree. As input, we used genomes from the Boce in house dataset as well as genomes from NCBI^22^.

**Supplementary References**

1.	Bode, E., Heinrich, A.K., Hirschmann, M., Abebew, D., Shi, Y.N., Vo, T.D., Wesche, F., Shi, Y.M., Grun, P., Simonyi, S., et al. (2019). Promoter Activation in Deltahfq Mutants as an Efficient Tool for Specialized Metabolite Production Enabling Direct Bioactivity Testing. Angew Chem Int Ed Engl *58*, 18957-18963. 10.1002/anie.201910563.

2.	Firacative, C., Khan, A., Duan, S., Ferreira-Paim, K., Leemon, D., and Meyer, W. (2020). Rearing and Maintenance of Galleria mellonella and Its Application to Study Fungal Virulence. J Fungi (Basel) *6*. 10.3390/jof6030130.

3.	McMullen, J.G., 2nd, and Stock, S.P. (2014). In vivo and in vitro rearing of entomopathogenic nematodes (Steinernematidae and Heterorhabditidae). J Vis Exp, 52096. 10.3791/52096.

4.	Orozco, R.A., Lee, M.M., and Stock, S.P. (2014). Soil sampling and isolation of entomopathogenic nematodes (Steinernematidae, Heterorhabditidae). J Vis Exp. 10.3791/52083.

5.	Bode, E., He, Y., Vo, T.D., Schultz, R., Kaiser, M., and Bode, H.B. (2017). Biosynthesis and function of simple amides in Xenorhabdus doucetiae. Environ Microbiol *19*, 4564-4575. 10.1111/1462-2920.13919.

6.	Wagner, T., Merino, F., Stabrin, M., Moriya, T., Antoni, C., Apelbaum, A., Hagel, P., Sitsel, O., Raisch, T., Prumbaum, D., et al. (2019). SPHIRE-crYOLO is a fast and accurate fully automated particle picker for cryo-EM. Commun Biol *2*, 218. 10.1038/s42003-019-0437-z.

7.	Punjani, A., Rubinstein, J.L., Fleet, D.J., and Brubaker, M.A. (2017). cryoSPARC: algorithms for rapid unsupervised cryo-EM structure determination. Nat Methods *14*, 290-296. 10.1038/nmeth.4169.

8.	Goddard, T.D., Huang, C.C., Meng, E.C., Pettersen, E.F., Couch, G.S., Morris, J.H., and Ferrin, T.E. (2018). UCSF ChimeraX: Meeting modern challenges in visualization and analysis. Protein Sci *27*, 14-25. 10.1002/pro.3235.

9.	Abramson, J., Adler, J., Dunger, J., Evans, R., Green, T., Pritzel, A., Ronneberger, O., Willmore, L., Ballard, A.J., Bambrick, J., et al. (2024). Accurate structure prediction of biomolecular interactions with AlphaFold 3. Nature *630*, 493-500. 10.1038/s41586-024-07487-w.

10.	Adams, P.D., Afonine, P.V., Bunkoczi, G., Chen, V.B., Davis, I.W., Echols, N., Headd, J.J., Hung, L.W., Kapral, G.J., Grosse-Kunstleve, R.W., et al. (2010). PHENIX: a comprehensive Python-based system for macromolecular structure solution. Acta Crystallogr D Biol Crystallogr *66*, 213-221. 10.1107/S0907444909052925.

11.	Emsley, P., Lohkamp, B., Scott, W.G., and Cowtan, K. (2010). Features and development of Coot. Acta Crystallogr D Biol Crystallogr *66*, 486-501. 10.1107/S0907444910007493.

12.	Bozhuyuk, K.A.J., Fleischhacker, F., Linck, A., Wesche, F., Tietze, A., Niesert, C.P., and Bode, H.B. (2018). De novo design and engineering of non-ribosomal peptide synthetases. Nat Chem *10*, 275-281. 10.1038/nchem.2890.

13.	Katoh, K., and Standley, D.M. (2013). MAFFT multiple sequence alignment software version 7: improvements in performance and usability. Mol Biol Evol *30*, 772-780. 10.1093/molbev/mst010.

14.	Minh, B.Q., Schmidt, H.A., Chernomor, O., Schrempf, D., Woodhams, M.D., von Haeseler, A., and Lanfear, R. (2020). IQ-TREE 2: New Models and Efficient Methods for Phylogenetic Inference in the Genomic Era. Mol Biol Evol *37*, 1530-1534. 10.1093/molbev/msaa015.

15.	Jones, D.T., Taylor, W.R., and Thornton, J.M. (1992). The rapid generation of mutation data matrices from protein sequences. Comput Appl Biosci *8*, 275-282. 10.1093/bioinformatics/8.3.275.

16.	Lee, M.D. (2019). GToTree: a user-friendly workflow for phylogenomics. Bioinformatics *35*, 4162-4164. 10.1093/bioinformatics/btz188.

17.	Hyatt, D., LoCascio, P.F., Hauser, L.J., and Uberbacher, E.C. (2012). Gene and translation initiation site prediction in metagenomic sequences. Bioinformatics *28*, 2223-2230. 10.1093/bioinformatics/bts429.

18.	Eddy, S.R. (2011). Accelerated Profile HMM Searches. PLoS Comput Biol *7*, e1002195. 10.1371/journal.pcbi.1002195.

19.	Edgar, R.C. (2022). Muscle5: High-accuracy alignment ensembles enable unbiased assessments of sequence homology and phylogeny. Nat Commun *13*, 6968. 10.1038/s41467-022-34630-w.

20.	Capella-Gutierrez, S., Silla-Martinez, J.M., and Gabaldon, T. (2009). trimAl: a tool for automated alignment trimming in large-scale phylogenetic analyses. Bioinformatics *25*, 1972-1973. 10.1093/bioinformatics/btp348.

21.	Price, M.N., Dehal, P.S., and Arkin, A.P. (2010). FastTree 2--approximately maximum-likelihood trees for large alignments. PLoS One *5*, e9490. 10.1371/journal.pone.0009490.

22.	Goldfarb, T., Kodali, V.K., Pujar, S., Brover, V., Robbertse, B., Farrell, C.M., Oh, D.H., Astashyn, A., Ermolaeva, O., Haddad, D., et al. (2025). NCBI RefSeq: reference sequence standards through 25 years of curation and annotation. Nucleic Acids Res *53*, D243-D257. 10.1093/nar/gkae1038.
