## Supplementary Movies 1-3 for "Acyl amides derived from a bacterial symbiont control development of its nematode host"

**Supplementary Videos 1–3.**

Representative phase-contrast microscopy videos of amide-treated *S. diaprepesi* infective juveniles undergoing exsheathment by shedding the retained second-stage cuticle associated with the free-living stage, preceding transition to the parasitic stage.
